# Prolific S-layer shedding and associated proteins from the methanotroph *Methylomicrobium album* BG8

**DOI:** 10.1101/2025.01.06.631565

**Authors:** Mariah K. Hermary, Maria C. Rodriguez Gallo, Rachael Rieberger, Aurelija M. Grigonyte, Kieran McDonald, R. Glen Uhrig, Dominic Sauvageau, Lisa Y. Stein

## Abstract

Some methanotrophs synthesize S-layers that overlay their outer membrane. TEM imaging revealed that *Methylomicrobium album* BG8 constitutively sheds abundant S-layer units, a phenotype not found in 7 other methanotrophs, even though *Methylotuvimicrobium buryatense* 5GB1 produces a similar structure. Release of S-layer units occurred regardless of carbon (methane or methanol) or nitrogen (ammonium or nitrate) source, with 50X trace metals, under copper deprivation, and at all growth phases. The released S-layer units were isolated from the culture medium of *M. album* BG8 by density gradient centrifugation for proteome analysis. The proteome revealed the S-layer protein subunits, transporters for calcium uptake including TolC and Repeats-in-Toxin (RTX) proteins, transporters for uptake for cobalamin and siderophores, cell wall biogenesis proteins, and proteins with Type I secretion system (T1SS) target domains. *M. album* BG8 adapted to grow at pH 4 lost its S-layer and genome analysis revealed a frameshift mutation plus reduced expression of the S-layer unit gene plus the deletion and almost no expression of an S-layer-associated porin gene. Together, the results suggest that the biogenesis and secretion of *M. album* BG8 S-layer is mediated by its associated T1SS, the S-layer possesses metal acquisition functions, and low pH adaptation of *M. album* BG8 results in loss of S-layer, likely due to reduced, or incomplete, expression of S-layer units and loss of an associated porin. The involvement of the T1SS and shedding phenotype of the S-layer in *M. album* BG8 could be applied towards selective secretion of proteins and other factors of bioindustrial interest.

**Importance:** The methanotrophic bacterium *M. album* BG8 produces and sheds large quantities of S-layer units into the culture medium regardless of carbon or nitrogen source, metal availability or growth phase. Of the 8 methanotrophic bacteria screened, only *M. album* BG8 possessed this phenotype. Proteomics analysis of density gradient purified culture supernatant identified the S-layer protein units and proteins involved in metal uptake and S-layer biogenesis, some with secretion signals for the T1SS. *M. album* BG8 adapted to grow at low pH lost production of its S-layer due to mutations in the genes encoding S-layer units and an associated porin. Better understanding of *M. album* BG8 S-layer production and its shedding phenotype could be harnessed for exporting expressed proteins and bioproducts of industrial interest for ease of collection and downstream processing.

## Introduction

Aerobic methanotrophic bacteria are clustered within the Gammaproteobacteria, Alphaproteobacteria and Verrucomicrobia phyla and are metabolically differentiated by their discrete pathways of carbon assimilation (1). Collectively, methanotrophs are of industrial interest for their capacity to convert methane, a potent greenhouse gas, into valuable molecules, from biopolymers to proteins to biofuels (2, 3). The Gammaproteobacterium *Methylomicrobium album* BG8 is a high priority strain for industrialization due to its metabolic flexibility for rapid growth on methane or methanol, with nitrate or ammonium as its nitrogen source (4, 5), and availability of a well-curated genome-scale metabolic model (6). Development of methanotrophs as industrial chassis relies on their ability to express materials of commercial interest at scale, requiring optimization of biomass production and possibly engineering strains to secrete valuable metabolites for ease of downstream processing (3). The diversity of substrates secreted by Type 1 secretion systems (T1SS) makes this system attractive for production and extracellular collection of heterologous proteins, biologics (e.g. growth factors), biosensors, and other molecules (7). Methanotrophs with an active T1SS could potentially be engineered to consume methane for the synthesis and secretion of valuable products that promote biodegradation, metal-scavenging, plant growth, and more, enabling simultaneous greenhouse gas removal and its valorization.

Structural analysis of methanotroph cell surfaces has revealed that some strains possess unique S-layers comprised of tightly packed cup-shaped protein units with p6 hexagonal symmetry (8–11). It was hypothesized that these structures could provide rigidity to the cell wall and play a role in osmoregulation and tolerance to extremes of salinity and alkalinity (11). A recent study of S-layers from two alkaliphilic *Methylotuvimicrobium* strains validated this hypothesis (12). However, non-halotolerant and neutralophilic methanotrophs, such as *Methylomicrobium album* BG8 (13) and *Methylomicrobium* sp. HG-1 (14), express similar arrays of these cup-like structures on their surfaces, which brings their physiological role and mechanism of biogenesis into question. *Caulobacter crescentus* forms similarly structured S-layers from cup-shaped RsaA proteins with p6 symmetry (15). These RsaA proteins are secreted through a T1SS and contain a span of six repeat-in-toxin (RTX) domains that bind calcium to enable proper folding and stabilization of the crystalline S-layer lattice on the outer membrane (16). Mutations in the N-terminus membrane attachment site of the RsaA protein results in a prolific shedding phenotype (17). Comparison of the *M. album* BG8 RsaA homolog to that of the methanotrophs *Methylotuvimicrobium buryatense* 5GB1 and *Methylotuvimicrobium alcaliphilum* 20Z showed only ca. 20% sequence similarity (12). However, all three methanotroph genomes possess a complete T1SS operon along with their RsaA gene, suggesting that S-layer biogenesis relies on the T1SS similar to that of *C. crescentus*.

The present study shows that *M. album* BG8 constitutively sheds its cup-shaped S-layer units under a variety of nutrient conditions. The cup-shaped S-layer units from the supernatant of *M. album* BG8 cultures were purified using density gradient centrifugation, and proteomic analysis by LC-MS/MS was performed. *M. album* BG8 was adapted to grow at acidic pH, which caused the loss of its S-layer altogether. The observations from this study provide understanding of the morphology, biogenesis, and associated protein functions for released S-layer units of *M. album* BG8, including involvement of a T1SS and calcium binding, plus its complete loss when challenged with growth at low pH. Insights into the biogenesis of S-layer protein units and their prolific release can be used as tools in the production and secretion of valuable industrial proteins using single-carbon molecules as feedstocks.

## Results

### *M. album* BG8 constitutively releases abundant cup-shaped S-layer units

Transmission electron microscopy of *M. album* BG8 displayed cup-shaped particles (Fig. 1) with diameters ranging from 49.5 to 75.2 nm, with an average of 60.9 ± 5.7 nm (n=50; Supp. Table S1). Released S-layer units were observed when *M. album* BG8 was grown in NMS or AMS media containing methane or methanol as a sole carbon source, media devoid of copper, and media containing 50X trace elements solution (examples of TEM images for various conditions are shown in Fig.1). Thus, release of S-layer protein units was considered constitutive, as it did not vary with changes in carbon source, nitrogen source, or metal availability. Released S-layer protein units were also present immediately after culture inoculation, in mid-log growth and at stationary phase (images not shown). However, released S-layer protein units were not detected in the medium of 7 other methanotrophic strains examined (Fig. 2), including cultures of *Methylotuvimicrobium buryatense* 5GB1, which possess similar cup-shaped S-layer lattices on its cell surface (11).

**Figure 1.**
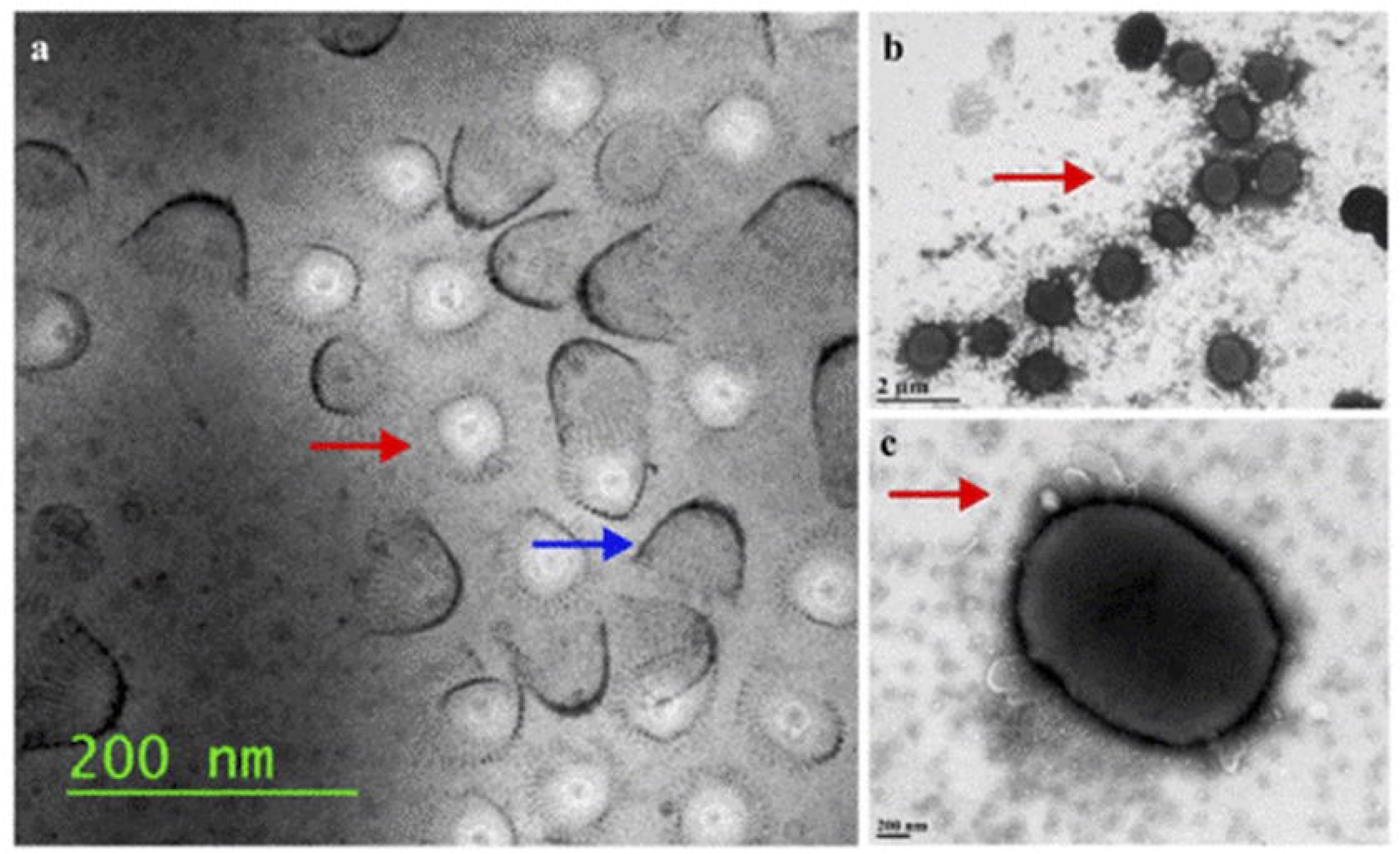
Transmission electron micrographs of *M. album* BG8 with released S-layer protein units (as depicted with arrows) when cultured with methane in various media conditions. (a) NMS media (b) AMS media without copper (c) NMS media with 50X trace elements solution. Micrographs contain a wide range of scales to highlight membrane morphology and adjacent surrounding areas.

**Figure 2.**
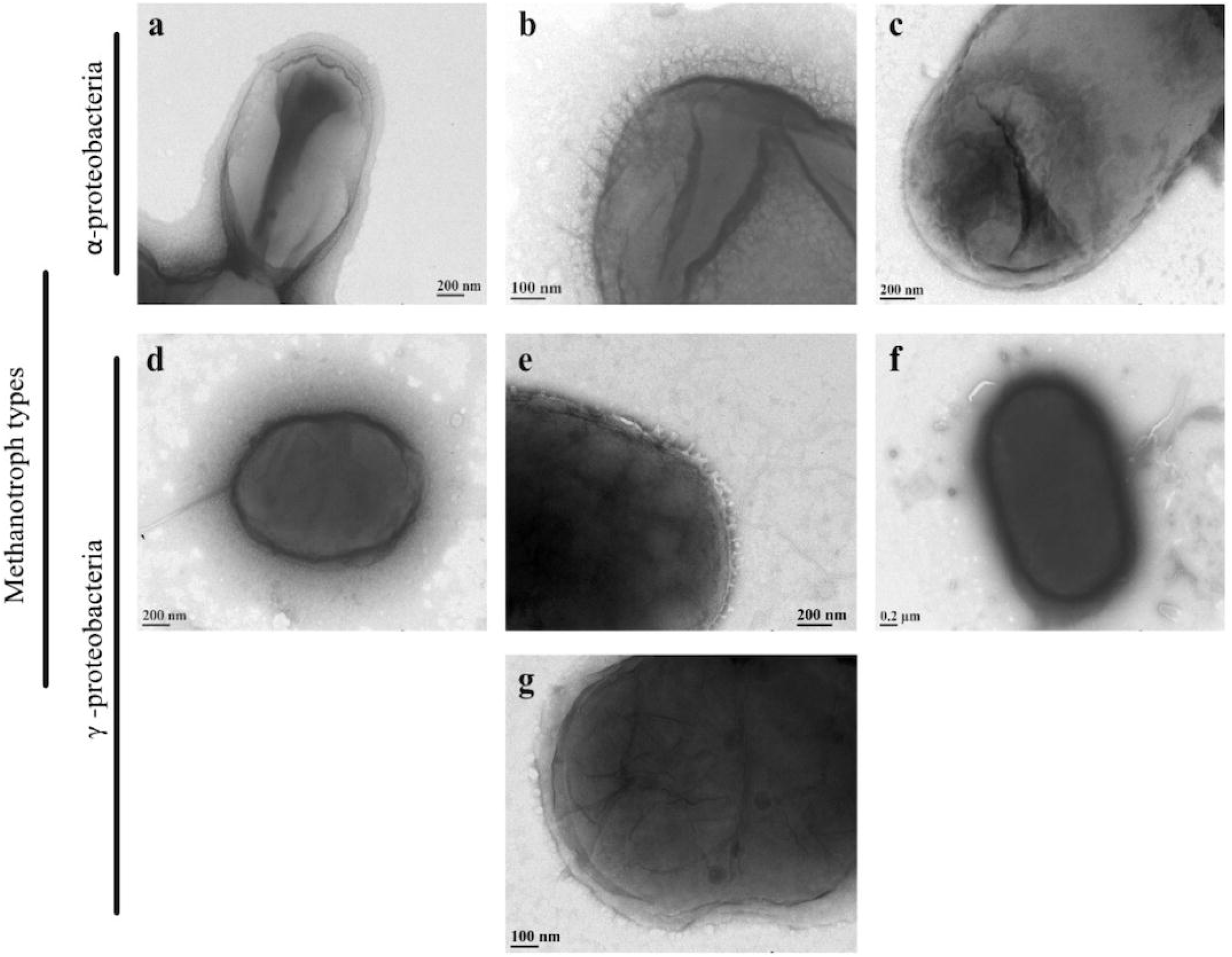
Transmission electron micrographs of selected alpha- and gammaproteobacterial methanotrophs cultivated on methane and NMS. (a) *Methylocystis* sp. WRRC1 (b) *Methylocystis* sp. Rockwell (c) *Methylosinus trichosporium* OB3b (d) *Methylomonas denitrificans* FJG1 (e) *Methylicorpusculum oleiharenae* XLMV4^T^ (f) *Methylotuvimicrobium buryatense* 5GB1 (g) *Methylococcus capsulatus* Bath.

### S-layer unit proteome suggests associated functions of T1SS, calcium uptake, metal acquisition, and membrane assembly

From the total of 310 peptides identified by LC-MS/MS from purified S-layer units released into the medium of *M. album* BG8 cultures (Supp. Table S2), 14 had high statistical significance across triplicate samples (ranking PSM >4, peptides >5, and score>20). Of note were the S-layer protein itself (H8GFV3), proteins related to binding and uptake of calcium and S-layer assembly (Ca^2^-binding RTX protein, PtrC metalloprotease, pre-pilin N-terminal cleavage) (16, 18, 19), proteins for siderophore and divalent cation uptake (TonB receptor) and cobalamin transport (OM receptor), a porin protein (H8GGQ6), a protein with a T1SS C-terminal target domain (H8GIF1), and a LPS-assembly protein (Table 1, Fig. 3).

**Table 1.** Significant peptide hits from LC-MS/MS detection of purified released S-layer protein units (n=3) organized by Genbank locus tag.

| Genbank Locus Tag | Uniprot Accession | UniProt Description | Mean PSM | Mean peptide | Mean score | COG Category |
| --- | --- | --- | --- | --- | --- | --- |
| Metal_0147 | H8GKK7 | Ca<sup>2+</sup>-binding protein, RTX toxin | 5.7 | 5.7 | 18.5 | Secondary metabolites biosynthesis, transport and catabolism |
| Metal_0479 | H8GNA1 | Uncharacterized protein (Metallo-protease PrtC*) | 30.3 | 15.7 | 106.7 | Secondary metabolites biosynthesis, transport and catabolism |
| Metal_0659 | H8GPP0 | Uncharacterized protein | 7 | 1 | 20 | Inorganic ion transport and metabolism |
| **Metal_0880** | **H8GFV3** | **S-layer protein** | **7.3** | **6.3** | **23.2** | **Secondary metabolites biosynthesis, transport and catabolism** |
| Metal_1116 | H8GHU9 | Prepilin-type N-terminal cleavage/methylation domain-containing protein | 64.3 | 4 | 231.4 | Intracellular trafficking, secretion, and vesicular transport |
| Metal_1447 | H8GKR0 | Outer membrane protein beta-barrel domain-containing protein* | 5 | 2 | 17.9 | Cell wall/membrane/envelope biogenesis |
| Metal_1728 | H8GMU5 | Flagellin | 331.7 | 22.3 | 1316.7 | Cell motility |
| Metal_1783 | H8GNF7 | Flagellar hook protein FlgE | 16 | 8.7 | 51.7 | Cell motility |
| Metal_2279 | H8GGQ6 | Uncharacterized porin | 16 | 4.7 | 52.5 | Cell wall/membrane/envelope biogenesis |
| Metal_2337 | H8GGW0 | TonB-dependent siderophore receptor | 7 | 6.3 | 20.2 | Inorganic ion transport and metabolism |
| Metal_2634 | H8GJI6 | LPS-assembly protein LptD | 16.7 | 2.7 | 43.8 | Cell wall/membrane/envelope biogenesis |
| Metal_3768 | H8GIA0 | ATP synthase epsilon chain | 9.7 | 1 | 22.4 | Lipid transport and metabolism |
| Metal_3821 | H8GIF1 | Type I secretion C-terminal target domain protein | 24.5<sup>+</sup> | 12.5<sup>+</sup> | 71.9<sup>+</sup> | Secondary metabolites biosynthesis, transport and catabolism |
| Metal_3919 | H8GJ49 | Outer membrane cobalamin receptor protein | 8 | 6.3 | 22.5 | Coenzyme transport and metabolism |

\*HHpred protein annotation (>95% probability), "+" replicates n=2, bolded protein = major S-layer unit

**Figure 3.**
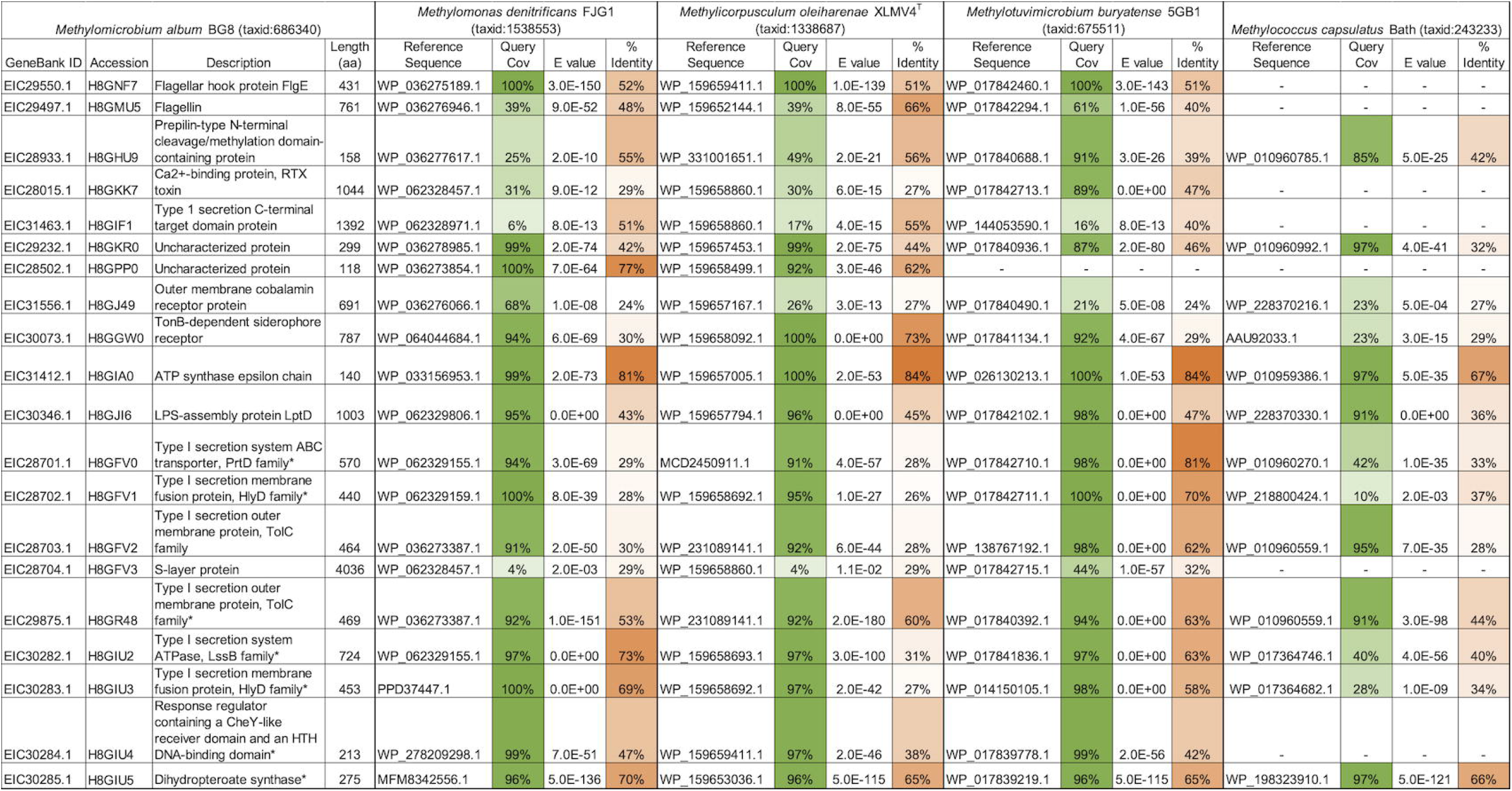
Protein sequence comparison of S-layer and T1SS-related proteins of *M. album* BG8 to gammaproteobacterial methanotrophs *M. denitrificans* FJG1, *M. oleiharenae* XLMV4^T^, *M. buryatense* 5GB1 and *M. capsulatus* Bath.

Proteins with relevance to S-layer biogenesis, but with lower statistical significance (PSM ≤5, peptides ≤5, score ≤20), included TolC, which is the outer-membrane channel component of the T1SS that interacts with the Ca^2+^ binding domains of RTX proteins for their secretion (Table 2) (20). Several membrane-related proteins with lower statistical significance included a cation diffusion facilitator protein, a general secretion pathway “protein D”, and TolB from the Tol-Pal system that may have been surreptitiously attached to the released S-layer proteins units. OmpA protein, also associated with the S-layer proteins, may play a role in S-layer biogenesis or tethering to the cell wall as it is an abundant and vital protein in the outer membrane of Gram negative bacteria (21). CorA, a protein once thought to anchor S-layer proteins to the cell wall (10), was only found in one of the triplicate proteome samples at low peptide counts. High counts of flagella-related proteins were co-purified, likely as contaminants from the culture supernatant.

**Table 2.** Additional protein hits of interest from LC-MS/MS detection of purified released S-layer protein units organized by Genbank locus tag.

| Genbank Locus Tag | UniProt Accession | UniProt Description | Mean PSM | Mean Peptide | Mean Score | COG Category |
| --- | --- | --- | --- | --- | --- | --- |
| Metal_0094 | H8GK04 | Cation diffusion facilitator family transporter | 2.5 | 1.00<sup>b</sup> | 6.19 | Inorganic ion transport and metabolism |
| Metal_0482 | H8GNA4 | Uncharacterized protein | 4.67 | 2 | 11.77 | - |
| **Metal_0879** | **H8GFV2** | **Type I secretion outer membrane protein, TolC family** | **3** | **3** | **8.09** | **Cell wall/membrane /envelope biogenesis; Intracellular trafficking, secretion, vesicular tnspt.** |
| Metal_1031 | H8GH59 | PAS domain S-box/diguanylate cyclase (GGDEF) domain-containing protein | 3.50<sup>a</sup> | 1 | 9.16 | Signal transduction mechanisms |
| Metal_1419 | H8GK80 | Outer membrane protein/peptidoglycan-associated (Lipo)protein | 3 | 2.33 | 8.32 | Cell wall/membrane /envelope biogenesis |
| Metal_1666 | H8GM80 | Autotransporter secretion inner membrane protein TamB | 3.33 | 1 | 7.05 | - |
| Metal_1730 | H8GMU7 | Flagellar hook-associated protein 2 | 5 | 5 | 13.49 | Cell motility |
| Metal_1789 | H8GNG3 | Flagellar hook-associated protein 1 | 2.67 | 2.67 | 8.48 | Cell motility |
| Metal_1790 | H8GNG4 | Flagellar hook-associated protein 3 | 2.33 | 2.33 | 6.55 | Cell motility |
| Metal_1986 | H8GPX1 | Citrate lyase beta subunit | 2.33 | 1 | 8.01<sup>c</sup> | Carbohydrate transport and metabolism |
| Metal_1998 | H8GPY3 | Outer membrane protein/peptidoglycan-associated (Lipo)protein | 2.67 | 2.33 | 7.49 | Cell wall/membrane /envelope biogenesis |
| Metal_2207 | H8GG32 | Adenosylhomocysteinase | 3.33 | 1.33 | 4.8 | Coenzyme transport and metabolism |
| Metal_2282 | H8GGQ9 | DNA-binding transcriptional regulator NtrC | 2.67 | 1 | 6.79 | Signal transduction mechanisms |
| Metal_2345 | H8GGW4 | Uncharacterized protein | 2.67 | 1 | 6.78 | Function unknown |
| Metal_2446 | H8GI24 | Uncharacterized protein | 4.00<sup>a</sup> | 1 | 10.57 | - |
| Metal_2469 | H8GI44 | PQQ-dependent dehydrogenase, methanol/ethanol family | 3 | 3 | 7.4 | Carbohydrate transport and metabolism |
| Metal_2481 | H8GI56 | Uncharacterized protein | 2.50<sup>a</sup> | 2.5 | 6.29 | - |
| Metal_2515 | H8GI88 | Probable cytosol aminopeptidase | 2.33 | 2 | 6.21 | Amino acid transport and metabolism |
| Metal_2819 | H8GL29 | Tol-Pal system protein TolB | 4.67 | 4.33 | 14.22 | Intracellular trafficking, secretion, and vesicular transport |
| Metal_282<br>1 | H8GL31 | Cell division coordinator<br>CpoB | 5 | 5 | 11.65 | Cell cycle control, cell division,<br>chromosome partitioning |
| Metal_305<br>5 | H8GN21 | Uncharacterized protein | 4 | 1 | 8.92 | Function unknown |
| Metal_310<br>6 | H8GNM<br>8 | Hemolysin-coregulated<br>protein | 3.67 | 3 | 10.3 | Function unknown |
| Metal_325<br>5 | H8GPH9 | Murein lipoprotein | 1.67 | 1.67 | 6.06 | Function unknown |
| Metal_333<br>8 | H8GQ61 | Uncharacterized protein | 3 | 2.67 | 7.27 | Function unknown |
| Metal_346<br>0 | H8GRD0 | Transaldolase | 4.33 | 4 | 10.21 | Nucleotide transport and<br>metabolism |
| Metal_367<br>9 | H8GHK8 | Gly-zipper_OmpA<br>domain-containing<br>protein | 3.5 | 2 | 12.03<sup>c</sup> | Function unknown |
| Metal_382<br>2 | H8GIF2 | VCBS repeat-containing<br>protein | 2.67 | 2 | 11.46<sup>c</sup> | Secondary metabolites<br>biosynthesis, transport and<br>catabolism |
| Metal_382<br>8 | H8GIF8 | General secretion<br>pathway protein D | 3 | 2.5 | 7.94<sup>c</sup> | Cell motility; Intracellular<br>trafficking, secretion, and<br>vesicular transport |

\* HHpred protein annotation (>95%)
<sup>a</sup> Absent in replicate A
<sup>b</sup> Absent in replicate B
<sup>c</sup> Absent in replicate C

Homologs of significantly identified proteins from the proteome were searched for in the genomes of the four gammaproteobacterial methanotrophs screened in this study. Only *M. buryatense* 5GB1 encoded the T1SS gene cluster that includes a homolog of the RsaA S-layer protein (Accession: H8GFV0-H8GFV3); this was expected as this was the only other methanotroph we included that has a S-layer similar to that of *M. album* BG8 (12) (Fig. 3). Although not expressed in the supernatant proteome, a second gene cluster – encoding T1SS ATPase, T1SS HylD membrane fusion protein, a response regulator, and a dihydropteroate synthase (Accession: H8GIU2-H8GIU5) – was identified along with a nearby gene for T1SS TolC (Accession: H8GR48). Sequences of these genes in other methanotrophs showed varying percentages of sequence identity to the genes from *M. album* BG8 (Fig. 3).

### Loss of S-layer in a strain of *M. album* BG8 adapted to grow at low pH

To test whether growth at acidic pH had a destabilizing effect on S-layer biogenesis and shedding, as has been shown in *Caulobacter crescentus* (17), we conducted adaptive laboratory evolution on *M. album* BG8 to grow in media from pH 6.8 to 4. Successive transfers in acidified NMS medium resulted in three separate lineages of low pH-adapted cultures. TEM images of the adapted cultures grown on methane or methanol showed smooth cell surfaces devoid of S-layer and the absence of shed S-layer particles into their culture medium (Fig. 4). Genome sequences between the low pH-adapted and parental (unadapted) *M. album* BG8 strains were compared. A frameshift mutation in the S-layer-encoding gene (Metal_0880) at nucleotide 2987/12111, with a Leu substitution at amino acid 996/4036, was identified (Supp. Data, Table 3). Transcriptomic analysis showed that expression of the Metal_0880 gene and its associated T1SS genes (Metal_0877-0879) (12) was ca. 50% lower in the low pH-adapted than in the parental *M. album* BG8 grown in NMS medium at pH 6.8 (Table 3) (5). A frameshift mutation in the gene for a highly expressed porin protein (H8GGQ6) in the S-layer proteome was also identified (Metal_2279), and its transcription was nearly abolished in the low pH-adapted compared to the parental strain (Table 3). These two mutations alone may have resulted in the smooth phenotype of the low pH-adapted strain due to incomplete S-layer unit expression and stabilization or anchoring of S-layer units to the cell wall by the H8GGQ6 porin protein. Thus, adapting *M. album* BG8 to grow at low pH resulted in mutations relevant to S-layer biogenesis, yielding complete loss of its S-layer (Fig. 4). No difference in growth rate was detected between the low pH-adapted and parental *M. album* BG8 strains when cultivated in NMS medium at pH 6.8 (Supp. Fig. S1). Only the low pH-adapted strains could grow in NMS medium at pH 4, albeit at a slower growth rate than at pH 6.8 (Supp. Fig. S1). The results suggest that the energetic cost of S-layer production did not affect the growth rate of *M. album* BG8 in neutral pH media and that the low pH adaptation process resulted in loss of its S-layer.

**Table 3.** Mutations and level of transcription (Transcripts per million) identified in the genome and RNAseq data of *M. album* BG8 adapted to growth at pH 4. The adapted genome was compared to the parental genome. Gene expression levels in the adapted and parental transcriptomes are reported. The two genes in bold were identified in the S-layer

| Genbank Locus Tag | UniProt Accession | Type of Mutation | UniProt Function | TPM<sup>(1)</sup> Adapted | TPM<sup>(1)</sup> Parental |
| --- | --- | --- | --- | --- | --- |
| Metal_0463 | H8GN86 | Synonymous | Serine/threonine protein kinase | 105 | 61 |
| Metal_0492 | H8GNB4 | Synonymous | Malto-oligosyltrehalose synthase | 28 | 3 |
| *\*Metal_0877* | *H8GFV0* | *none* | *Type I secretion system ABC transporter, PrtD family* | *128* | *252* |
| *\*Metal_0878* | *H8GFV1* | *none* | *Type I secretion membrane fusion protein, HlyD family* | *97* | *167* |
| *\*Metal_0879* | *H8GFV2* | *none* | *Type I secretion outer membrane protein, TolC family* | *81* | *181* |
| **Metal_0880** | **H8GFV3** | **Frameshift (delT at nt 2987; Leu at aa 996)** | **S-layer protein** | **1764** | **3948** |
| Metal_0915 | H8GFY7 | Insertion | Drug resistance transporter, EmrB/QacA subfamily | 46 | 67 |
| Metal_0961 | H8GGI4 | Frameshift | Choline dehydrogenase-like flavoprotein | 104 | 20 |
| Metal_1085 | H8GHB2 | Synonymous | Ketol-acid reductoisomerase (NADP(+)) | 449 | 703 |
| Metal_1113 | H8GHU6 | Glu – Val | Uncharacterized protein | 154 | 44 |
| Metal_1464 | H8GKS7 | Asp – Gly | DNA-binding protein H-NS | 2960 | 2618 |
| Metal_1726 | H8GMU4 | Asp – Ser | Flagellar protein FliL | 70 | 22 |
| Metal_1910 | H8GP90 | Synonymous | Rhodanese-related sulfurtransferase | 56 | 66 |
| Metal_1927 | H8GPA7 | Val – Ile | Putative PLP-dependent enzyme possibly involved in cell wall biogenesis | 115 | 96 |
| Metal_2019 | H8GQF3 | Asp – Asn | Succinate dehydrogenase hydrophobic membrane anchor subunit | 135 | 105 |
| Metal_2096 | H8GQM4 | Synonymous | Putative thymidine phosphorylase | 77 | 76 |
| Metal_2101 | H8GR29 | Val – Gly | Probable membrane transporter protein | 18 | 8 |
| Metal_2132 | H8GR59 | Synonymous | Cadherin domain-containing protein | 66 | 35 |
| Metal_2140 | H8GR66 | Val – Leu | Protein-export membrane protein SecG | 767 | 1324 |
| **Metal_2279** | **H8GGQ6** | **inframe deletion** | **Porin** | **21** | **865** |
| Metal_2323 | H8GGU6 | Glu – Gly | Cytochrome c | 45 | 58 |
| Metal_2426 | H8GHJ6 | Deletion | RHS repeat-associated core domain protein | 20 | 5 |
| Metal_2567 | H8GIU3 | Deletion | Membrane fusion protein (MFP) family protein | 132 | 91 |
| Metal_2569 | None** | Insertion | TolC family outer membrane protein | 873 | 428 |
| Metal_2678 | H8GJM5 | Glu – Ala | Tfp pilus assembly protein PilN | 95 | 48 |
| Metal_2685 | H8GJN2 | Leu – Pro | C-terminal processing peptidase | 215 | 407 |
| Metal_2779 | H8GKZ0 | Deletion | Transcriptional regulator | 35 | 40 |
| Metal_3143 | H8GNR1 | 1 missense, 2 synonymous | Transposase | 40 | 40 |
| Metal_3171 | H8GNT9 | Ser – Asn | Phosphatidylserine decarboxylase proenzyme | 72 | 271 |
| Metal_3777 | H8GIA9 | Val – Ala | Cobalt-zinc-cadmium resistance protein CzcC | 60 | 53 |
| Metal_3987 | H8GJT4 | Leu – Pro | Transglutaminase-like domain-containing protein | 36 | 93 |

<sup>(1)</sup>TPM: mRNA transcripts per million; data was collected for *M. album* BG8 growing in NMS medium at pH 6.8.
\*mRNA expression values of the three T1SS genes associated with the S-layer structural gene are reported to compare between the low pH adapted and parental strains.
\*\* UniProt accession number not available; protein function supported by InterProScan and blastp annotation.

**Figure 4.**
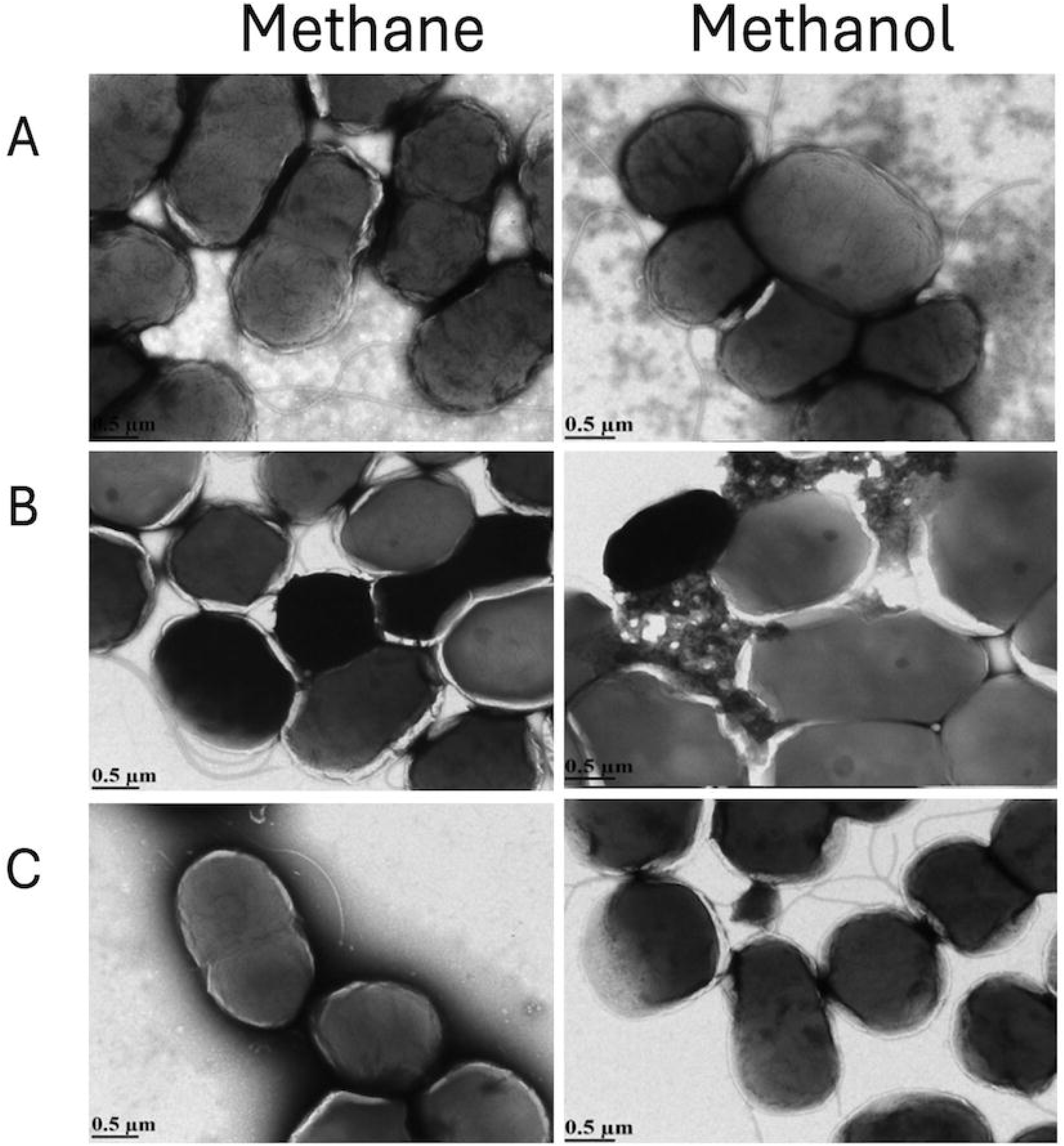
A) *M. album* BG8 (parental strain, unadapted) grown in NMS at pH 6.8 in methane or methanol showing prolific shedding of S-layer particles and the presence of flagella in the surrounding media. B) *M. album* BG8 adapted to grow at pH 4, cultivated in methane or methanol, showing absence of S-layer particles in the surrounding media. C) Low pH-adapted *M. album* BG8 grown at pH 6.8 in methane or methanol NMS showing continued absence of S-layer particles in the surrounding media.

## Discussion

*M. album* BG8 was initially isolated by Roger Whittenbury in 1971 and has been continuously maintained in culture ever since (22). It is likely that the dissociation of its S-layer observed here was acquired by random mutation over the thousands of generations that occurred in laboratory cultures and not a natural feature of the original isolate. Analysis of dissociated S-layer units offers the opportunity to understand more about the S-layer structure, its biogenesis, and its purpose in *M. album* BG8; and provides a basis for the potential utilization of the shedding phenotype to produce and collect products of industrial interest. Based on the most abundant proteins with annotated functions in the proteome of released *M. album* BG8 S-layer units, its biogenesis appears to involve calcium-mediated secretion of S-layer proteins via Ca^2+^-binding RTX domains of the associated T1SS (19, 20) and assembly with assistance of several outer membrane and other RTX-domain proteins (18, 19), including anchoring to the cell wall by highly expressed porin or other outer membrane proteins (23). The T1SS has three components: an ABC transporter located within the inner membrane, a membrane fusion protein that spans the periplasmic space, and an outer membrane porin protein to create an ATP-dependent translocation mechanism (20). TolC, a highly represented protein in the *M. album* BG8 S-layer unit proteome, forms the outer membrane porin component of the T1SS and transports Ca^2+^-binding proteins via a Ca^2+^ ratcheting mechanism (20). The protein targets of the T1SS are typically defined by several blocks of nonapeptide-binding consensus sequences located at their C-termini (24). The identification of significant RTX exoprotein and T1SS C-terminal target domain protein in the S-layer unit proteome suggests these T1SS-secreted proteins are associated with the S-layer structure. The significant presence of associated PrtC metalloprotease suggests a putative role in S-layer biogenesis or maintenance, analogous to what is observed with the Sap metalloprotease of *C. crescentus* (25).

The S-layer protein (H8GFV3) of *M. album* BG8 is 4,036 amino acids long and contains a conserved hemolysin-type (RTX) calcium-binding region (IPR018511) (19) and a CalX-like domain (IPR038081), with only 35% amino acid identity to the S-layer protein identified in *M. buryatense* 5GB1 (Fig. 3). S-layer proteins are known to display broad diversity between species (26) and the amino acid identity observed between the sequences of S-layer proteins from *M. album* BG8 and *M. buryatense* 5GB1 is above the typical threshold required to consider shared functions (27). In addition, similarly to what is found in *M. album* BG8, a Ca^+^ binding domain was identified in the RsaA S-layer protein of *C. crescentus* (15). Other Gammaproteobacteria that encode homologues to the H8GFV3 S-layer protein of *M. album* BG8 include the close relative *Methylomicrobium lacus* (49% AA ID) and several strains of *Marinobacterium georgiense* (36-37% AA ID)(Supp. Table S2) for which S-layers and membrane blebs have been reported (28). The S-layer proteins of *M. album* BG8, *M. buryatense* 5GB1 and *M. alcaliphilum* 20Z are encoded in gene clusters along with cognate T1SS proteins (H8GF0, H8GF1, H8GF2) (29). Here, we showed congruent expression of the three T1SS and S-layer protein genes in both the low pH-adapted and parental strains of *M. album* BG8 through transcriptomic analysis (Table 3), suggesting co-regulation of this gene cluster. Exposure of *C. cresentus* to low pH causes release of its S-layer, which has been used to collect S-layer particles for deeper analysis (25). For *M. album* BG8, the S-layer was already actively dissociating from the cell at neutral pH, and further adaptation to low pH resulted in complete loss of S-layer, likely due to a frameshift mutation in the S-layer-encoding gene and a deletion of an S-layer-associated porin gene (Table 3).

The copper regulated CorA and CorB proteins of *M. album* BG8 possess Ca^2+^ binding channels (30, 31), and in conjunction with the TonB-ExbB-ExbD transporter system, they can mediate copper and divalent cation uptake and ion homeostasis (30). Correspondingly, two of the proteins with high counts in the S-layer unit proteome were the TonB siderophore receptor protein (H8GGW0) and a Ca^2+^ binding protein (H8GKK7). However, CorA was only found with low counts in one proteome replicate (Supp. Table S2). Although CorA was once considered as a possible anchor point for S-layer proteins (10), a recent study on S-layer of *M. buryatense* 5GB1 showed that its deletion did not affect S-layer production or attachment but did affect copper uptake (12). As neither copper starvation nor growth in 50X trace element solution changed the pattern of S-layer production and release in *M. album* BG8, the shedding phenotype does not appear to be regulated by the abundance of trace metals. Alternatively, the highly expressed porin protein (H8GGQ6) that contained a deletion in the low pH-adapted *M. album* BG8, or the outer membrane beta-barrel domain protein (H8GKR0), could be investigated as potential cell wall anchors similarly to the porin-like proteins of *Deinococcus radiodurans* (23). It is also possible that the LPS assembly protein (H8GJI6) could be a linkage of S-layer to the cell wall in a calcium-dependent fashion similar that of *C. crescentus* (32).

Although S-layers can be produced by various methanotrophs, *M. album* BG8 was the only one out of 8 examined from both Alpha- and Gammaproteobacteria to prolifically release S-layer units into the culture. Structures in TEM images surrounding methanotrophs other than *M. album* BG8 included white patches from the phosphotungistic acid negative stain (Fig. 2), membrane protrusions around the surface of *M. oleiharenae* XLMV4^T^ cells (Fig. 2e) (33), sparse spherical particles around some *M. buryatense* 5GB1 cells (Fig. 2f), and capsule layer surrounding *M. capsulatus* Bath (Fig. 2g) (22). None of these structures resembled the released S-layer proteins of *M. album* BG8, but the patchy occurrence of S-layers and capsules among the methanotrophic bacteria continues to be an intriguing topic for future study.

### Application of the T1SS and released S-layer proteins for bioindustrial protein production

The industrialization of methanotrophs has been growing in appeal due to the potential to convert methane, a potent greenhouse gas and cheap feedstock, into value-added products such as small metabolites, biofuel precursors, biopolymers, and proteins (34). The phenotype of *M. album* BG8 wherein its S-layer units are delivered from the cytoplasm to the cell surface by the T1SS (35) and then released to the external environment, represents an advantageous protein-production platform. Using the shedding S-layer system, different strategies can be targeted: a heterologous protein can be fused to residues used for excretion through the T1SS for expression of single protein molecules, or the protein of interest can be fused to the N-terminal of the S-layer protein (36) to produce functionalized S-layer units within the culture medium. Of course, the success of such strategies is dependent on the structure and function of the protein of interest but, in each case, the recovery of a heterologous product may be facilitated by the excretion and release mechanism of S-layer units, greatly simplifying downstream processing and reducing associated costs.

### Conclusion

This study identified and characterized the S-layer shedding phenotype of *M. album* BG8 that is not shared by 7 other screened methanotrophic strains. The prolific production and release of *M. album* BG8 S-layer units into the culture medium – likely a consequence of long-term cultivation of this strain – allowed for their purification and proteomic analysis. We identified co-expression of the S-layer protein gene with its neighboring T1SS genes, and in the proteome, we observed the S-layer units, TolC proteins, proteins with T1SS signal sequences, metal-acquisition and membrane biogenesis proteins, and a prominent porin. *M. album* BG8 adapted to grow at low pH through adaptive laboratory evolution acquired mutations in genes encoding S-layer protein and the prominent porin, resulting in low or abolished expression, respectively. These two mutations may be responsible for the constitutive loss of S-layer in the low pH-adapted strain. In *C. crescentus*, S-layer proteins comprise up to 30% to total cellular protein, and considerable energy is required to generate and release S-layer proteins. Yet, *M. album* BG8 grows efficiently and rapidly on both methane and methanol, with a relatively high growth rate compared to other methanotrophs despite its constitutive production and shedding of S-layer (4). The S-layer production and shedding phenotype of *M. album* BG8 presents an opportunity to produce and secrete bioproducts of industrial interest including heterologous proteins, growth factors, and metal-chelating agents, among others. Although the cellular energy demands for constitutive S-layer production are high, an added benefit with using *M. album* BG8 is that increased methane consumption will foster optimized rates and yields of bioproduction, which can also yield a larger net climate benefit.

## Materials and Methods

### Culturing conditions and media

Eight methanotroph strains maintained in the laboratory were used: Gammaproteobacteria - *Methylomicrobium album* BG8, *Methylomonas denitrificans* FJG1, *Methylicorpusculum oleiharenae* XLMV4T, *Methylococcus capsulatus* Bath and *Methylotuvimicrobium buryatense* 5GB1; Alphaproteobacteria - *Methylocystis* sp. WRRC1, *Methylocystis* sp. Rockwell, and *Methylosinus trichosporium* OB3b. Methanotrophs were cultivated in 250-mL Wheaton bottles fitted with screw cap lids inlaid with a butyl rubber septum filled with 100 mL of nitrate (NMS) or ammonium mineral salts (AMS) medium (4), or NMS2 medium (37) for *M. buryatense* 5GB1. Gas headspace (50 mL) was removed from each bottle after which methane (60 mL) was injected via syringe fitted with a 0.22-µm filter. For *M. album* BG8 grown on methanol, 100 µL of high-performance liquid chromatography grade methanol (Sigma-Aldrich) was added. Experiments with increased trace metals included 50X standard trace element solution. Experiments with copper-free media included Chelex 100 (BioRad)-treated media (except the trace-elements solution) added to acid-washed Wheaton bottles. 1 mL (1%) inoculum from a late exponential phase culture that had been passaged at least once under the identical growth conditions for each experimental condition was used for each strain. Cultures were incubated at 30°C, with the exception of 37°C for *M. capsulatus* Bath, with shaking at 150 rpm until early stationary phase as determined by growth curves (OD540) (4).

### Adaptation of M. album BG8 to low pH and analysis of genome-wide mutations

A series of NMS medium formulations from pH 6.8 to 3.8 in 0.2 pH unit increments were generated using mixtures of KH_2_PO_4_, Na_2_HPO_4_ and citrate phosphate buffers with further acidification by dropwise addition of 0.1 M HCl to achieve the desired final pH prior to autoclaving. Cultures were grown on methane/air (30/70%) and passaged a minimum of three times before moving to the next lower pH medium. Adaptation experiments were performed from 2022-2024. Visualization of the adapted cells was performed by TEM as described below. Genomic DNA was extracted from three replicated culture lines from parental (non-adapted) and pH 4-adapted *M. album* BG8 grown on NMS medium at pH 6.8 using the DNeasy Blood and Tissue Kit (Qiagen), cleaned using the ZymoDNA clean and concentrator kit (Zymo Research, USA) and measured by the Qubit dsDNA broad range assay (Fisher Scientific, USA). Genomes were sequenced using Ilumina Novoseq (Genome Quebec, Montreal) with 1060X (adapted) and 1173X coverage. The adapted genome was aligned to the parental genome using SPAdes (v4.2.0+galaxy0). Snippy (v4.6.0+galaxy0) was used to identify mutations in the consensus adapted genome.

### Transmission electron microscopy imaging

Cells (ca. 2 mL) were removed from cultures and pelleted by centrifugation (5 min at 6000 x g; Eppendorf). Cell pellets were rinsed and resuspended in 500 µL phosphate buffer. Cell samples were placed on a 300-nm copper grid for 4 to 6 min prior to staining with 1% phosphotungistic acid. Images were captured using a Philips/FEI Transmission Electron Microscope with Gatan Camera (Morgagni). Fig. 2 was imaged with a JEOL JEM-ARM200CF S/TEM electron microscope at an accelerating voltage of 200 kV. The HAADF-STEM images were collected with the JEOL HAADF detector using the following experimental conditions: probe size 6C, condenser lens aperture 30 μm, scan speed 32 μs per pixel, and camera length 8 cm. The average diameter of S-layer proteins starting from the outer edge were measured using ImageJ (n=50).

### S-layer purification

Three technical replicates were used for purification and LC MS/MS proteomics with each replicate deriving from 1 L of *M. album* BG8 culture grown in NMS medium with methane. The purification method is illustrated in Fig. 5, and was adapted from a protocol for isolation of outer membrane vesicles (38). For each replicate, the cultures were filtered using a 0.45-μm PES stericup (Millipore) to separate biomass from filtrate containing the S-layer proteins. The filtrate was passed through a 100 kDa MW cutoff protein concentrator (GE Vivaspin) and retentate was collected by rinsing the filter with 1 mL phosphate buffer at pH 6.8. Retentate aliquots were spotted on NMS agar and incubated with methane to confirm the absence of viable *M. album* BG8 cells.

**Figure 5.**
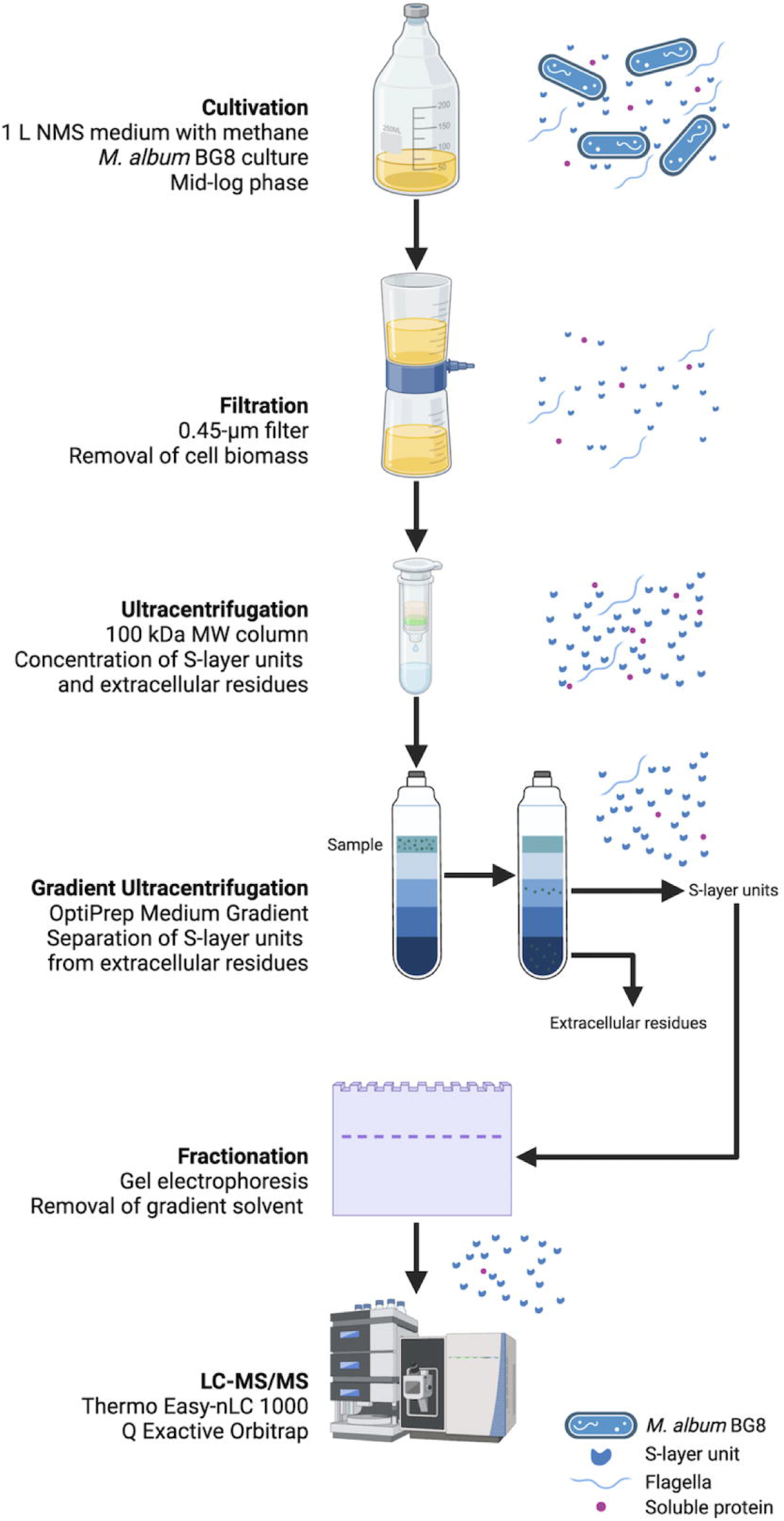
S-layer isolation and purification protocol based on density gradient centrifugation for *M. album* BG8 adapted from a protocol for OMV isolation (38).

The crude retentate was mixed with OptiPrep (iodixanol; Sigma-Aldrich) (60%) and placed at the bottom of a sterile centrifuge tube. A discontinuous density gradient was created by layering 2 mL of diluted OptiPrep (40, 35, 30, 25, 20% using OptiPrep Diluent) as per manufacturer’s instructions. Samples were centrifuged for 12 h at 53,000 rpm (~288,350 x *g*) at 4°C in a Ti70 Fixed rotor ultracentrifuge (Beckman Optima L-90K Ultracentrifuge). 1-mL fractions were removed by pipetting from the top of the density gradient, with a total of 18 fractions per replicate. Each fraction was measured for refractive index (Reichert AR200 Automatic Digital Refractometer) to ensure that the density gradient was maintained. Per fraction, 50 μL was denatured using 50 μL of 2x Laemmli Sample Buffer (BioRad) and incubation on a heat block (Baxter Canlab H2025-1 Dry Block Heater) at 95°C for 5 min. The prepared samples were separated on an 8% acrylamide gel and stained with colloidal Coomassie to determine protein presence. Protein-containing fractions were denatured and concentrated using a protein centrifuge concentrator (Amicon) to a final volume of 50 μL. Concentrated protein samples were partially resolved through a precast 4-20% acrylamide gradient gel (BioRad) to remove the Optiprep solvent until no residual sample was present in the wells of the gel. Protein samples were excised from the gel and submitted for LC-MS/MS analysis.

### In-Gel Digest of S-layer Peptides

The gel bands were cut into small pieces and destained four times with 50% Acetonitrile (ACN; Sigma) in 50⍰mM trietylammonium bicarbonate (TEAB; Sigma) at 37°C for 10 min. Destained gel pieces were washed with 100 mM TEAB at 37°C for 10 min and dehydrated by incubating them twice in 100 % ACN at room temperature for 10 min to complete drying. Cysteine residues were reduced by incubating dried gel slices in 10⍰mM dithiothreitol (DTT; BioRad) solution in 100 mM TEAB for 45 min at 37°C. Reduction solution was replaced with 55⍰mM iodoacetamide (IA; Sigma) in 100 mM TEAB buffer and incubated in the dark for 1 h at 37°C to alkylate cysteine residues. The alkylation solution was replaced by 100⍰mM TEAB and incubated for 10⍰min, repeated twice. Gel pieces were then dehydrated once more in 100% ACN and dried as stated above. Gel pieces were rehydrated by adding 6⍰ng/μL trypsin (Promega; V5113) in 100⍰mM TEAB. Peptides were digested for 16 h at 37 °C with shaking at 150 rpm. Tryptic peptides were retained, and in-gel digested peptides were further extracted by incubating in 1% (v/v) formic acid (FA; Fisher), 2% (v/v) ACN in 100 mM TEAB for 1 h at 37°C. This was followed by a second 1 h 37°C extraction using a 1:1 1% (v/v) FA in 100 mM TEAB and 100% ACN extraction buffer. Digested peptide fractions were pooled and dried. Isolated peptides were re-suspended and desalted using ZipTip C18 pipette tips (ZTC18S960; Millipore), as described (39). All peptides were dried and re-suspended in 3% (v/v) ACN / 0.1% (v/v) FA immediately prior to mass spectrometry analysis.

### Liquid chromatography mass spectrometry

Re-suspended tryptic peptides were analyzed using nano flow HPLC (Easy-nLC 1000, Thermo Scientific) with an EASY-Spray capillary HPLC column (PepMap RSLC C18, 75um x 25cm, 100Å, 2μm, Thermo Scientific) coupled to a Q Exactive Orbitrap mass spectrometer (Thermo Scientific). Dissolved samples were injected using an Easy-nLC 1000 system (Thermo Scientific). The column was equilibrated with 100% solvent A (0.1% (v/v) FA, 4% (v/v) ACN in water). Peptides were eluted using the following gradient of solvent B (0.1% (v/v) FA in 80% (v/v) ACN): 0–18% B, 0–73 min; 18–30% B, 73-101 min; 30–46% B, 101–120 min; 46–100% B, 120-123, min at a flow rate of 0.35 µl min−1 at 50°C. High-accuracy mass spectra were acquired in data-dependent acquisition mode. All precursor signals were recorded in a mass range of 300– 1,700 m/z and a resolution of 35,000 at 200 m/z. The maximum accumulation time for a target value of 1 × 10^6^ was set to 120 ms. Up to 12 data-dependent MS/MS were recorded using quadrupole isolation with a window of 2 Da and higher-energy collisional dissociation fragmentation with 26% fragmentation energy. A target value of 5 × 10^4^ was set for MS/MS using a maximum injection time of 250 ms and a resolution of 17,500 at 200 m/z. Precursor signals were selected for fragmentation with charge states from +2 to +7 and a signal intensity of at least 1 × 10^4^. All precursor signals selected for MS/MS were dynamically excluded for 30s.

All acquired mass spectrometry data was analyzed using Proteome Discoverer (ver. 2.4.1.15). Data were searched against the *M. album* BG8 proteome (UNIPROT H8GRI8_METAL; containing 3,738 protein sequences) using an automatically generated decoy database and the Proteome Discoverer processing workflow (PWF_QE_Basic_SeaquestHT) and consensus workflow (CWF_Basic) with minimal deviation. Deviations included: a precursor and fragment mass tolerance of 10 ppm and 0.6 da, respectively, trypsin digestion with 2 missed cleavages, variable modifications methionine oxidation (+15.995 Da), n-terminal acetylation (+42.011 Da), methionine loss (−131.040 Da), methionine loss + n-terminal acetylation (−89.030 Da) and fixed modifications carbamidomethylation (+57.021 Da).

### Identifying similar protein sequences in other methanotrophs

To determine if identified S-layer-associated proteins were unique to *M. album* BG8, each identified protein sequence was compared using BLASTp to the gammaproteobacterial methanotrophs: *Methylomonas denitrificans* FJG1 (taxon:416); *Methylicorpusculum oleiharenae* XLMV4T (taxon:1338687); *Methylotuvimicrobium buryatense* 5GB1 (taxon:1338687); and *Methylococcus capsulatus* Bath (taxon:414). (https://blast.ncbi.nlm.nih.gov/). The presence of T1SS proteins identified in the *M. album* BG8 genome (GenBank: CM001475.1) were also analyzed for their presence in the above gammaproteobacterial genomes. For query sequences with a high number of BLASTp hits, only matches with the highest percent identities to the query were reported.

To determine the presence of S-layer related proteins in gammaproteobacterial methanotrophs that were not analyzed in this experiment, the proteins of interest were screened against the entire *Methylococcaceae* group (taxid:135618) using BLASTp (Supp. Table S3). The S-layer protein H8GFV3 was further screened via BLASTp against all Gammaproteobacteria (taxid:1236) (Supp. Table S4).

### RNA extraction and RNAseq analysis

Total RNA was extracted from triplicate late-log phase cultures of the low-pH adapted *M. album* BG8 grown in NMS medium at pH 6.8 (Supp. Fig. S1) using the QIAWave RNA Mini Kit (Qiagen) following the manufacturer’s protocol. The extracted RNA was quantified using the Qubit RNA high sensitivity assay (Fisher Scientific, USA) and quality controlled using the Agilent 2100 Bioanalyzer. RNA libraries were prepared by Genome Quebec (Montreal, Canada) using a bacterial rRNA-depleted RNA-seq library preparation workflow, followed by sequencing on an Illumina NovaSeq (PE100, ~25 M reads per sample, ±5 M). Raw paired-end RNAseq reads were analyzed using the Galaxy web platform (usegalaxy.eu; version 25.0). First, reads were quality filtered using Trimmomatic (v0.39+galaxy2) with default parameters. High-quality reads were mapped to the low-pH adapted *M. album* BG8_1aa genome using HISAT2 (v2.2.1+galaxy0). The number of reads mapped to each gene was calculated by HTseq-count (v2.0.9+galaxy0) and the output was used to calculate normalized transcript abundances in transcripts per million (TPM). Statistically validated TPM values for the same genes from the parental *M. album* BG8 strain grown in NMS media with methane were used from a prior RNAseq experiment with triplicate samples (5).

## Supporting information

supp_FigS1-tablesS1-S4

## Acknowledgments

The authors would like to thank Arlene Oatway and Dr. Kacie Norton from the Biological Sciences Microscopy Unit and Haoyang Yu from NanoFAB with assisting in transmission electron microscopy imaging. All peptide digestion and LC-MS/MS preparation were performed at the Alberta Proteomics and Mass Spectrometry Facility at the Univ. Alberta.

## Data availability

All raw proteomics files were submitted to the PRoteomics IDEntification Database (PRIDE; https://www.ebi.ac.uk/pride/). Reviewer access details are as follows: Dataset: PXD050176, Username, Password: pGJMeScS.. The genome and RNAseq data for low pH-adapted *M. album* BG8 grown in NMS at pH 6.8 can be found under Genbank accession number PRJNA698057. Transcriptome data for parental *M. album* BG8 grown in NMS at pH 6.8 can be found under accession Genbank number PRJNA698057. Further information on the genome mutations and calculations of RNAseq TPM values for the low pH-adapted strain are in the Supp. Data Sheet.

## Author Contributions: Author Contributions

M.K.H., D.S. and L.Y.S. conceptualized the research. M.K.H. carried out the experiments and created the figures and tables. R.G.U. and M.C.R.G. provided peptide digestion, LC-MS/MS preparation and corresponding proteome analysis. A.M.G. provided bioinformatics analysis. R.R. performed the pH adaptation experiments, genome sequencing of the parental and adapted strains, and RNAseq analysis. M.K.H., D.S. L.Y.S. wrote the initial version of the manuscript. All authors edited the manuscript. D.S. and L.Y.S. co-supervised the work. All authors have given consent to the final version of the manuscript.

## Permission to print information from another source

Figure 5 was an adapted protocol with permission from Microbiological Research.

## Originality statement

None of the work in this manuscript has been published or submitted for publication elsewhere.

