## Supplementary material for "Prolific S-layer shedding and associated proteins from the methanotroph *Methylomicrobium album* BG8": supp_FigS1-tablesS1-S4

**Supplemental Figure S1.** Growth of low pH adapted and un-adapted *M. album* BG8 strains in NMS media at pH 6.8 (▲ = adapted; ◼ = un-adapted) and pH 4 (🞬 = adapted; ⚫ = un-adapted)

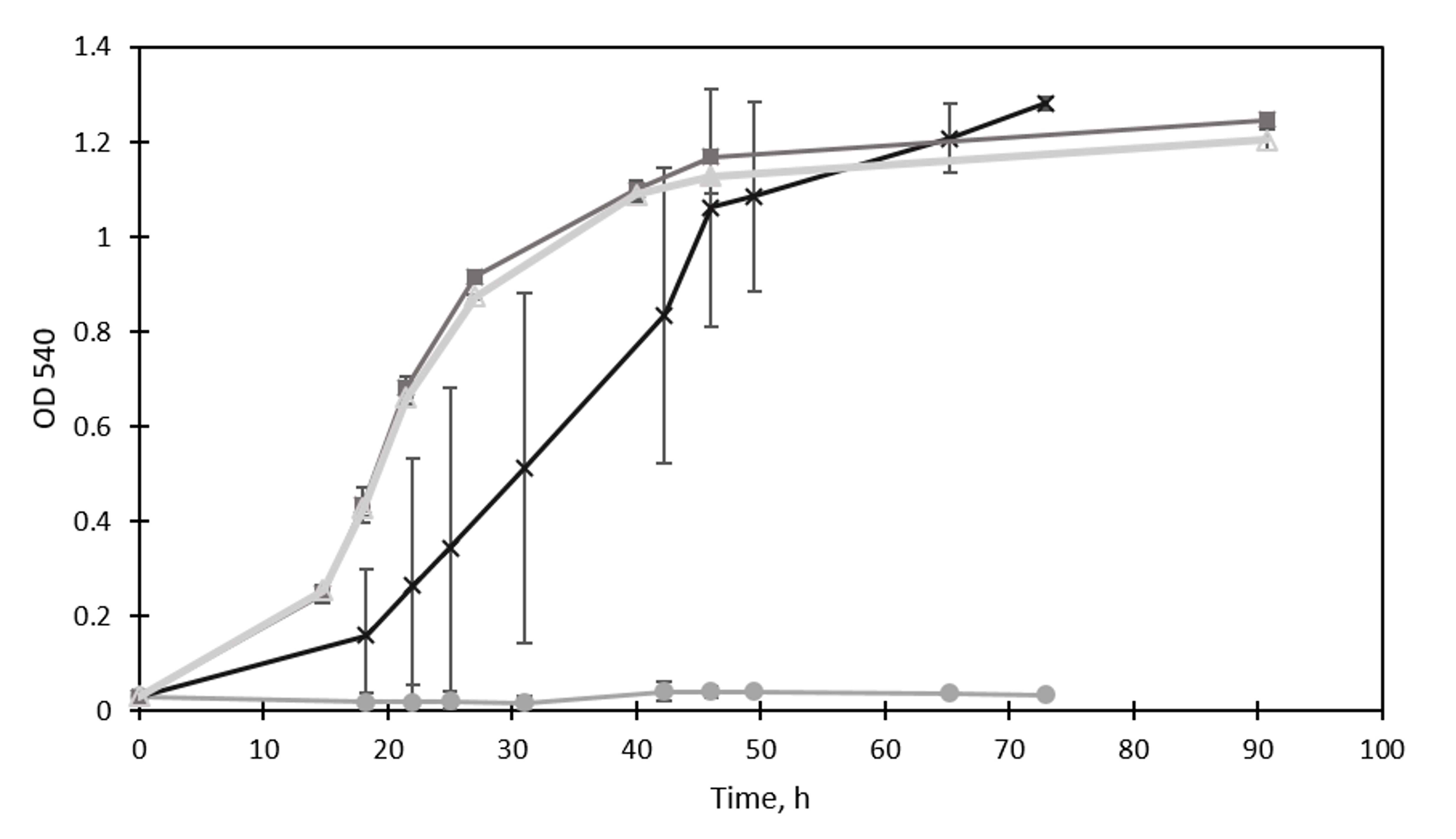

**Supplemental Table S1**. Average diameter of *M. album* BG8 S-layer protein units (n=50) per tested media condition.

| **Media Condition** | **Avg. Size (nm)** | | **Range (nm)** |
| --- | --- | --- | --- |
| CH_4_/NMS | | 60.9 ± 5.7 | 49.5 – 75.2 |
| CH_4_/AMS | | 64.5 ± 6.5 | 49.2 – 78.7 |
| CH_3_OH/NMS | | 64.8 ± 8.1 | 49.7 – 84.0 |
| CH_3_OH/AMS | | 66.9 ± 4.8 | 54.4 – 77.5 |
| CH_4_/x50 trace elements | | 61.0 ± 6.8 | 53.1 – 78.0 |
| CH_4_/vitamin stock | | 67.0 ± 6.7 | 49.4 – 82.0 |

**Supplemental Table S2.** All LC-MS/MS proteome hits from purified S-layer units of *M. album* BG8.

|  |  | Replicate A | | | Replicate B | | | Replicate C | | |
| --- | --- | --- | --- | --- | --- | --- | --- | --- | --- | --- |
| Accession | Description | PSM | peptide | score | PSM | peptide | score | PSM | peptide | score |
| H8GMU5 | Flagellin OS=Methylomicrobium album BG8 OX=686340 GN=Metal_1728 PE=3 SV=1 | 399 | 23 | 1512.12 | 428 | 23 | 1726.29 | 168 | 21 | 711.56 |
| H8GHU9 | Prepilin-type N-terminal cleavage/methylation domain-containing protein OS=Methylomicrobium album BG8 OX=686340 GN=Metal_1116 PE=3 SV=1 | 80 | 3 | 290.36 | 78 | 4 | 275.33 | 35 | 5 | 128.6 |
| H8GNA1 | Uncharacterized protein OS=Methylomicrobium album BG8 OX=686340 GN=Metal_0479 PE=4 SV=1 | 35 | 17 | 115.46 | 53 | 27 | 196.88 | 3 | 3 | 7.69 |
| H8GIF1 | Type 1 secretion C-terminal target domain protein OS=Methylomicrobium album BG8 OX=686340 GN=Metal_3821 PE=4 SV=1 | 30 | 12 | 89.27 | 19 | 13 | 54.51 | #N/A | #N/A | #N/A |
| H8GNF7 | Flagellar hook protein FlgE OS=Methylomicrobium album BG8 OX=686340 GN=Metal_1783 PE=3 SV=1 | 19 | 8 | 56.21 | 16 | 9 | 53.75 | 13 | 9 | 45.08 |
| H8GGQ6 | Uncharacterized protein OS=Methylomicrobium album BG8 OX=686340 GN=Metal_2279 PE=4 SV=1 | 14 | 4 | 45.02 | 26 | 6 | 87.31 | 8 | 4 | 25.19 |
| H8GIA0 | ATP synthase epsilon chain OS=Methylomicrobium album BG8 OX=686340 GN=atpC PE=3 SV=1 | 13 | 1 | 30.07 | 10 | 1 | 22.98 | 6 | 1 | 14.16 |
| H8GJI6 | LPS-assembly protein LptD OS=Methylomicrobium album BG8 OX=686340 GN=lptD PE=3 SV=1 | 12 | 1 | 31.37 | 28 | 6 | 74.49 | 10 | 1 | 25.47 |
| H8GJ49 | Outer membrane cobalamin receptor protein OS=Methylomicrobium album BG8 OX=686340 GN=Metal_3919 PE=3 SV=1 | 9 | 7 | 25.16 | 10 | 7 | 28.65 | 5 | 5 | 13.78 |
| H8GGW0 | TonB-dependent siderophore receptor OS=Methylomicrobium album BG8 OX=686340 GN=Metal_2337 PE=3 SV=1 | 7 | 7 | 21.15 | 12 | 10 | 33.06 | 2 | 2 | 6.32 |
| *H8GN21* | *Uncharacterized protein OS=Methylomicrobium album BG8 OX=686340 GN=Metal_3055 PE=3 SV=1* | 6 | 1 | 13.3 | 5 | 1 | 11.19 | 1 | 1 | 2.28 |
| H8GKK7 | Ca2+-binding protein, RTX toxin OS=Methylomicrobium album BG8 OX=686340 GN=Metal_0147 PE=4 SV=1 | 6 | 6 | 21.79 | 5 | 5 | 13.87 | 6 | 6 | 19.79 |
| *H8GNA4* | *Uncharacterized protein OS=Methylomicrobium album BG8 OX=686340 GN=Metal_0482 PE=4 SV=1* | 6 | 2 | 15.55 | 4 | 2 | 9.73 | 4 | 2 | 10.02 |
| *H8GNM8* | *Hemolysin-coregulated protein (Uncharacterized) OS=Methylomicrobium album BG8 OX=686340 GN=Metal_3106 PE=4 SV=1* | 5 | 4 | 13.52 | 4 | 3 | 11 | 2 | 2 | 6.38 |
| *H8GL31* | *Cell division coordinator CpoB OS=Methylomicrobium album BG8 OX=686340 GN=cpoB PE=3 SV=1* | 5 | 5 | 10.9 | 5 | 5 | 12.56 | 5 | 5 | 11.49 |
| H8GKR0 | Uncharacterized protein OS=Methylomicrobium album BG8 OX=686340 GN=Metal_1447 PE=4 SV=1 | 4 | 1 | 12.03 | 7 | 2 | 28.1 | 4 | 3 | 13.69 |
| *H8GIF2* | *VCBS repeat-containing protein OS=Methylomicrobium album BG8 OX=686340 GN=Metal_3822 PE=4 SV=1* | 4 | 2 | 11.5 | 4 | 2 | 11.41 | #N/A | #N/A | #N/A |
| H8GFV3 | Uncharacterized protein OS=Methylomicrobium album BG8 OX=686340 GN=Metal_0880 PE=4 SV=1 | 4 | 4 | 11.06 | 16 | 13 | 52.42 | 2 | 2 | 6.04 |
| *H8GI88* | *Probable cytosol aminopeptidase OS=Methylomicrobium album BG8 OX=686340 GN=pepA PE=3 SV=1* | 4 | 3 | 10.23 | 1 | 1 | 2.89 | 2 | 2 | 5.5 |
| *H8GL29* | *Tol-Pal system protein TolB OS=Methylomicrobium album BG8 OX=686340 GN=tolB PE=3 SV=1* | 4 | 3 | 11.74 | 5 | 5 | 17.2 | 5 | 5 | 13.72 |
| *H8GM80* | *Uncharacterized protein OS=Methylomicrobium album BG8 OX=686340 GN=Metal_1666 PE=4 SV=1* | 4 | 1 | 8.24 | 4 | 1 | 8.25 | 2 | 1 | 4.67 |
| *H8GFV2* | *Type I secretion outer membrane protein, TolC family OS=Methylomicrobium album BG8 OX=686340 GN=Metal_0879 PE=3 SV=1* | 4 | 4 | 9.69 | 4 | 4 | 11.76 | 1 | 1 | 2.82 |
| *H8GI44* | *PQQ-dependent dehydrogenase, methanol/ethanol family OS=Methylomicrobium album BG8 OX=686340 GN=Metal_2469 PE=3 SV=1* | 4 | 4 | 9.42 | 3 | 3 | 8.65 | 2 | 2 | 4.14 |
| *H8GHK8* | *Gly-zipper_OmpA domain-containing protein OS=Methylomicrobium album BG8 OX=686340 GN=Metal_3679 PE=4 SV=1* | 3 | 2 | 10.1 | 4 | 2 | 13.96 | #N/A | #N/A | #N/A |
| *H8GMU7* | *Flagellar hook-associated protein 2 OS=Methylomicrobium album BG8 OX=686340 GN=Metal_1730 PE=3 SV=1* | 3 | 3 | 7.33 | 5 | 5 | 13.8 | 7 | 7 | 19.33 |
| H8GNF2 | Histidine kinase OS=Methylomicrobium album BG8 OX=686340 GN=Metal_1778 PE=4 SV=1 | 3 | 1 | 6.33 | 2 | 1 | 4.7 | #N/A | #N/A | #N/A |
| *H8GK80* | *Outer membrane protein/peptidoglycan-associated (Lipo)protein OS=Methylomicrobium album BG8 OX=686340 GN=Metal_1419 PE=4 SV=1* | 3 | 2 | 6.94 | 4 | 3 | 12.53 | 2 | 2 | 5.5 |
| *H8GNG4* | *Flagellar hook-associated protein 3 OS=Methylomicrobium album BG8 OX=686340 GN=Metal_1790 PE=4 SV=1* | 2 | 2 | 5.57 | 2 | 2 | 4.76 | 3 | 3 | 9.31 |
| H8GKM9 | Thiol:disulfide interchange protein DsbA OS=Methylomicrobium album BG8 OX=686340 GN=Metal_0170 PE=3 SV=1 | 2 | 1 | 4.21 | #N/A | #N/A | #N/A | #N/A | #N/A | #N/A |
| H8GH31 | ABC-type antimicrobial peptide transport system, permease component OS=Methylomicrobium album BG8 OX=686340 GN=Metal_3659 PE=3 SV=1 | 2 | 1 | 4.01 | #N/A | #N/A | #N/A | #N/A | #N/A | #N/A |
| H8GK49 | Putative secreted protein OS=Methylomicrobium album BG8 OX=686340 GN=Metal_1386 PE=4 SV=1 | 2 | 1 | 3.95 | 2 | 1 | 4.2 | 4 | 1 | 7.83 |
| H8GJN2 | C-terminal processing peptidase OS=Methylomicrobium album BG8 OX=686340 GN=Metal_2685 PE=3 SV=1 | 2 | 2 | 4.51 | 2 | 2 | 5.22 | #N/A | #N/A | #N/A |
| H8GPH9 | Uncharacterized protein OS=Methylomicrobium album BG8 OX=686340 GN=Metal_3255 PE=4 SV=1 | 2 | 2 | 7.69 | 2 | 2 | 6.74 | 1 | 1 | 3.74 |
| H8GIB8 | Aspartate--tRNA(Asp/Asn) ligase OS=Methylomicrobium album BG8 OX=686340 GN=aspS PE=3 SV=1 | 2 | 1 | 3.86 | #N/A | #N/A | #N/A | #N/A | #N/A | #N/A |
| *H8GPX1* | *Citrate lyase beta subunit OS=Methylomicrobium album BG8 OX=686340 GN=Metal_1986 PE=4 SV=1* | 2 | 1 | 4.67 | 5 | 1 | 11.34 | #N/A | #N/A | #N/A |
| *H8GIF8* | *General secretion pathway protein D OS=Methylomicrobium album BG8 OX=686340 GN=Metal_3828 PE=3 SV=1* | 2 | 2 | 6.09 | 4 | 3 | 9.79 | #N/A | #N/A | #N/A |
| *H8GGQ9* | *DNA-binding transcriptional regulator NtrC OS=Methylomicrobium album BG8 OX=686340 GN=ntrC PE=4 SV=1* | 2 | 1 | 4.85 | 2 | 1 | 4.99 | 4 | 1 | 10.54 |
| *H8GNG3* | *Flagellar hook-associated protein 1 OS=Methylomicrobium album BG8 OX=686340 GN=flgK PE=3 SV=1* | 2 | 2 | 6.58 | 3 | 3 | 9.63 | 3 | 3 | 9.22 |
| H8GNG1 | Flagellar P-ring protein OS=Methylomicrobium album BG8 OX=686340 GN=flgI PE=3 SV=1 | 2 | 2 | 5.92 | 1 | 1 | 2.93 | #N/A | #N/A | #N/A |
| H8GK31 | DUF4114 domain-containing protein OS=Methylomicrobium album BG8 OX=686340 GN=Metal_1368 PE=4 SV=1 | 2 | 2 | 5.73 | 2 | 2 | 6.12 | #N/A | #N/A | #N/A |
| H8GKT0 | Dihydrolipoyllysine-residue succinyltransferase component of 2-oxoglutarate dehydrogenase complex OS=Methylomicrobium album BG8 OX=686340 GN=Metal_1467 PE=3 SV=1 | 2 | 1 | 4.46 | #N/A | #N/A | #N/A | 2 | 1 | 4.4 |
| H8GJN0 | Sulfite reductase, beta subunit (Hemoprotein) OS=Methylomicrobium album BG8 OX=686340 GN=Metal_2683 PE=4 SV=1 | 2 | 1 | 4.05 | 1 | 1 | 2.03 | #N/A | #N/A | #N/A |
| H8GPQ2 | Bacterioferritin (Cytochrome b1) OS=Methylomicrobium album BG8 OX=686340 GN=Metal_0675 PE=4 SV=1 | 2 | 1 | 4.32 | #N/A | #N/A | #N/A | #N/A | #N/A | #N/A |
| H8GPP0 | Uncharacterized protein OS=Methylomicrobium album BG8 OX=686340 GN=Metal_0659 PE=4 SV=1 | 2 | 1 | 5 | 15 | 1 | 42.99 | 4 | 1 | 12.02 |
| *H8GGW4* | *Uncharacterized protein OS=Methylomicrobium album BG8 OX=686340 GN=Metal_2345 PE=4 SV=1* | 2 | 1 | 5 | 2 | 1 | 5.68 | 4 | 1 | 9.66 |
| H8GQ90 | Sigma54-dependent transcription regulator containing an AAA-type ATPase domain and a DNA-binding domain OS=Methylomicrobium album BG8 OX=686340 GN=Metal_0714 PE=4 SV=1 | 2 | 2 | 4.31 | #N/A | #N/A | #N/A | 1 | 1 | 1.91 |
| *H8GRD0* | *Transaldolase OS=Methylomicrobium album BG8 OX=686340 GN=tal PE=3 SV=1* | 2 | 2 | 4.63 | 6 | 5 | 13.77 | 5 | 5 | 12.22 |
| *H8GG32* | *Adenosylhomocysteinase OS=Methylomicrobium album BG8 OX=686340 GN=ahcY PE=3 SV=1* | 2 | 1 | 4.13 | 6 | 2 | 8.27 | 2 | 1 | 1.99 |
| H8GQ43 | Elongation factor Tu OS=Methylomicrobium album BG8 OX=686340 GN=tuf PE=3 SV=1 | 1 | 1 | 2.65 | 1 | 1 | 2.49 | #N/A | #N/A | #N/A |
| H8GRI4 | UPF0102 protein Metal_3515 OS=Methylomicrobium album BG8 OX=686340 GN=Metal_3515 PE=3 SV=1 | 1 | 1 | 2 | #N/A | #N/A | #N/A | #N/A | #N/A | #N/A |
| H8GN32 | DNA/RNA helicase, superfamily II OS=Methylomicrobium album BG8 OX=686340 GN=Metal_3066 PE=3 SV=1 | 1 | 1 | 2.34 | #N/A | #N/A | #N/A | 1 | 1 | 2.37 |
| H8GGQ7 | Glutamine synthetase OS=Methylomicrobium album BG8 OX=686340 GN=Metal_2280 PE=3 SV=1 | 1 | 1 | 2.91 | #N/A | #N/A | #N/A | 2 | 2 | 4.95 |
| H8GM78 | Glycine dehydrogenase (decarboxylating) OS=Methylomicrobium album BG8 OX=686340 GN=gcvP PE=3 SV=1 | 1 | 1 | 2.29 | #N/A | #N/A | #N/A | #N/A | #N/A | #N/A |
| H8GHU8 | Uncharacterized protein OS=Methylomicrobium album BG8 OX=686340 GN=Metal_1115 PE=4 SV=1 | 1 | 1 | 2.42 | #N/A | #N/A | #N/A | #N/A | #N/A | #N/A |
| H8GQQ0 | ResIII domain-containing protein OS=Methylomicrobium album BG8 OX=686340 GN=Metal_3375 PE=4 SV=1 | 1 | 1 | 1.98 | #N/A | #N/A | #N/A | 1 | 1 | 2.06 |
| H8GQY4 | Uncharacterized protein OS=Methylomicrobium album BG8 OX=686340 GN=Metal_0813 PE=4 SV=1 | 1 | 1 | 2.96 | #N/A | #N/A | #N/A | 2 | 1 | 4.47 |
| H8GLB5 | Signal peptide peptidase SppA, 36K type OS=Methylomicrobium album BG8 OX=686340 GN=Metal_0251 PE=3 SV=1 | 1 | 1 | 2.38 | #N/A | #N/A | #N/A | 3 | 1 | 7.1 |
| H8GIE5 | Transposase_31 domain-containing protein OS=Methylomicrobium album BG8 OX=686340 GN=Metal_3815 PE=4 SV=1 | 1 | 1 | 2.15 | #N/A | #N/A | #N/A | #N/A | #N/A | #N/A |
| H8GIH5 | Periplasmic serine endoprotease DegP-like OS=Methylomicrobium album BG8 OX=686340 GN=Metal_1191 PE=3 SV=1 | 1 | 1 | 2.26 | #N/A | #N/A | #N/A | #N/A | #N/A | #N/A |
| H8GRD5 | 3-hexulose-6-phosphate synthase OS=Methylomicrobium album BG8 OX=686340 GN=Metal_3465 PE=3 SV=1 | 1 | 1 | 2.4 | 1 | 1 | 2.18 | #N/A | #N/A | #N/A |
| H8GGE8 | EAL domain-containing protein OS=Methylomicrobium album BG8 OX=686340 GN=Metal_3580 PE=4 SV=1 | 1 | 1 | 2.03 | 3 | 1 | 6.15 | 1 | 1 | 1.92 |
| H8GPJ5 | Thiol:disulfide interchange protein OS=Methylomicrobium album BG8 OX=686340 GN=Metal_0612 PE=3 SV=1 | 1 | 1 | 1.96 | #N/A | #N/A | #N/A | #N/A | #N/A | #N/A |
| H8GKS9 | Oxoglutarate dehydrogenase (succinyl-transferring) OS=Methylomicrobium album BG8 OX=686340 GN=Metal_1466 PE=3 SV=1 | 1 | 1 | 1.93 | 1 | 1 | 2.16 | 2 | 1 | 4.02 |
| H8GKK3 | Outer membrane protein assembly factor BamA OS=Methylomicrobium album BG8 OX=686340 GN=bamA PE=3 SV=1 | 1 | 1 | 2.74 | 2 | 2 | 4.94 | #N/A | #N/A | #N/A |
| H8GPQ4 | Uncharacterized protein OS=Methylomicrobium album BG8 OX=686340 GN=Metal_0677 PE=4 SV=1 | 1 | 1 | 1.96 | #N/A | #N/A | #N/A | #N/A | #N/A | #N/A |
| H8GI22 | Uncharacterized protein OS=Methylomicrobium album BG8 OX=686340 GN=Metal_2444 PE=4 SV=1 | 1 | 1 | 2.45 | 1 | 1 | 2.18 | #N/A | #N/A | #N/A |
| H8GNS0 | 3-hexulose-6-phosphate synthase OS=Methylomicrobium album BG8 OX=686340 GN=Metal_3152 PE=3 SV=1 | 1 | 1 | 1.96 | 1 | 1 | 3.52 | #N/A | #N/A | #N/A |
| H8GH58 | Uncharacterized protein OS=Methylomicrobium album BG8 OX=686340 GN=Metal_1030 PE=4 SV=1 | 1 | 1 | 2.12 | #N/A | #N/A | #N/A | #N/A | #N/A | #N/A |
| H8GQI9 | Uncharacterized protein OS=Methylomicrobium album BG8 OX=686340 GN=Metal_2057 PE=4 SV=1 | 1 | 1 | 2.09 | #N/A | #N/A | #N/A | #N/A | #N/A | #N/A |
| H8GN99 | Large extracellular alpha-helical protein OS=Methylomicrobium album BG8 OX=686340 GN=Metal_0477 PE=3 SV=1 | 1 | 1 | 1.9 | #N/A | #N/A | #N/A | #N/A | #N/A | #N/A |
| H8GJT9 | Outer membrane protein/peptidoglycan-associated (Lipo)protein OS=Methylomicrobium album BG8 OX=686340 GN=Metal_3992 PE=4 SV=1 | 1 | 1 | 2.05 | #N/A | #N/A | #N/A | #N/A | #N/A | #N/A |
| H8GPL2 | Adenine-specific DNA methylase containing a Zn-ribbon OS=Methylomicrobium album BG8 OX=686340 GN=Metal_0629 PE=4 SV=1 | 1 | 1 | 1.92 | #N/A | #N/A | #N/A | 1 | 1 | 1.97 |
| H8GL99 | UDP-N-acetylglucosamine 1-carboxyvinyltransferase OS=Methylomicrobium album BG8 OX=686340 GN=murA PE=3 SV=1 | 1 | 1 | 1.9 | #N/A | #N/A | #N/A | 1 | 1 | 2.58 |
| H8GJT1 | Uncharacterized protein OS=Methylomicrobium album BG8 OX=686340 GN=Metal_3984 PE=4 SV=1 | 1 | 1 | 2.5 | 1 | 1 | 2.74 | 1 | 1 | 2.29 |
| H8GID1 | Periplasmic serine endoprotease DegP-like OS=Methylomicrobium album BG8 OX=686340 GN=Metal_3800 PE=3 SV=1 | 1 | 1 | 2.94 | 2 | 2 | 5.25 | 2 | 2 | 5.11 |
| H8GKD1 | Fe2+-dicitrate sensor, membrane component OS=Methylomicrobium album BG8 OX=686340 GN=Metal_2720 PE=4 SV=1 | 1 | 1 | 2.95 | #N/A | #N/A | #N/A | #N/A | #N/A | #N/A |
| H8GGF2 | 60 kDa chaperonin OS=Methylomicrobium album BG8 OX=686340 GN=groL PE=3 SV=1 | 1 | 1 | 3.63 | #N/A | #N/A | #N/A | #N/A | #N/A | #N/A |
| H8GFX0 | Uncharacterized protein OS=Methylomicrobium album BG8 OX=686340 GN=Metal_0897 PE=4 SV=1 | 1 | 1 | 2.14 | #N/A | #N/A | #N/A | #N/A | #N/A | #N/A |
| H8GPU4 | Outer membrane protein/peptidoglycan-associated (Lipo)protein OS=Methylomicrobium album BG8 OX=686340 GN=Metal_1958 PE=4 SV=1 | 1 | 1 | 4.15 | 1 | 1 | 4.41 | 1 | 1 | 4.07 |
| H8GQH8 | Chaperone protein DnaK OS=Methylomicrobium album BG8 OX=686340 GN=dnaK PE=2 SV=1 | 1 | 1 | 3.27 | 2 | 2 | 5.25 | #N/A | #N/A | #N/A |
| H8GH44 | Uncharacterized protein OS=Methylomicrobium album BG8 OX=686340 GN=Metal_3672 PE=4 SV=1 | 1 | 1 | 2.15 | #N/A | #N/A | #N/A | #N/A | #N/A | #N/A |
| H8GMZ9 | Uncharacterized protein OS=Methylomicrobium album BG8 OX=686340 GN=Metal_3033 PE=4 SV=1 | 1 | 1 | 1.96 | 1 | 1 | 1.91 | 1 | 1 | 1.99 |
| H8GGM0 | Uncharacterized protein OS=Methylomicrobium album BG8 OX=686340 GN=Metal_0997 PE=4 SV=1 | 1 | 1 | 2.55 | 1 | 1 | 2.01 | #N/A | #N/A | #N/A |
| H8GQ63 | Peptide chain release factor 1 OS=Methylomicrobium album BG8 OX=686340 GN=prfA PE=3 SV=1 | 1 | 1 | 1.93 | #N/A | #N/A | #N/A | #N/A | #N/A | #N/A |
| H8GRI9 | Uncharacterized protein OS=Methylomicrobium album BG8 OX=686340 GN=Metal_4000 PE=4 SV=1 | 1 | 1 | 2.44 | #N/A | #N/A | #N/A | #N/A | #N/A | #N/A |
| H8GLG1 | Pyridoxine 5'-phosphate synthase OS=Methylomicrobium album BG8 OX=686340 GN=pdxJ PE=3 SV=1 | 1 | 1 | 1.92 | #N/A | #N/A | #N/A | #N/A | #N/A | #N/A |
| H8GPS4 | Type IV pilus biogenesis/stability protein PilW OS=Methylomicrobium album BG8 OX=686340 GN=Metal_1937 PE=4 SV=1 | 1 | 1 | 2.11 | 2 | 2 | 4.72 | #N/A | #N/A | #N/A |
| *H8GK04* | *Cation diffusion facilitator family transporter OS=Methylomicrobium album BG8 OX=686340 GN=Metal_0094 PE=4 SV=1* | 1 | 1 | 2.32 | #N/A | #N/A | #N/A | 4 | 1 | 10.05 |
| *H8GPY3* | *Outer membrane protein/peptidoglycan-associated (Lipo)protein OS=Methylomicrobium album BG8 OX=686340 GN=Metal_1998 PE=4 SV=1* | 1 | 1 | 1.96 | 4 | 3 | 12.67 | 3 | 3 | 7.83 |
| H8GN43 | Uncharacterized protein OS=Methylomicrobium album BG8 OX=686340 GN=Metal_3078 PE=4 SV=1 | 1 | 1 | 2.09 | #N/A | #N/A | #N/A | #N/A | #N/A | #N/A |
| H8GHZ8 | Outer membrane protein OS=Methylomicrobium album BG8 OX=686340 GN=Metal_1171 PE=3 SV=1 | 1 | 1 | 2.02 | #N/A | #N/A | #N/A | 1 | 1 | 2.26 |
| H8GQU1 | Uncharacterized protein OS=Methylomicrobium album BG8 OX=686340 GN=Metal_3423 PE=4 SV=1 | 1 | 1 | 2.28 | 1 | 1 | 2.32 | #N/A | #N/A | #N/A |
| H8GP62 | Uncharacterized protein OS=Methylomicrobium album BG8 OX=686340 GN=Metal_1882 PE=4 SV=1 | 1 | 1 | 2.6 | #N/A | #N/A | #N/A | #N/A | #N/A | #N/A |
| H8GMC0 | Uncharacterized protein OS=Methylomicrobium album BG8 OX=686340 GN=Metal_2963 PE=4 SV=1 | 1 | 1 | 1.98 | #N/A | #N/A | #N/A | #N/A | #N/A | #N/A |
| H8GKL3 | Uncharacterized protein OS=Methylomicrobium album BG8 OX=686340 GN=Metal_0154 PE=4 SV=1 | 1 | 1 | 1.9 | #N/A | #N/A | #N/A | #N/A | #N/A | #N/A |
| H8GP21 | Uncharacterized protein OS=Methylomicrobium album BG8 OX=686340 GN=Metal_0598 PE=4 SV=1 | 1 | 1 | 2.45 | #N/A | #N/A | #N/A | #N/A | #N/A | #N/A |
| H8GIS9 | Rnf electron transport complex subunit B OS=Methylomicrobium album BG8 OX=686340 GN=Metal_2552 PE=4 SV=1 | 1 | 1 | 2.08 | #N/A | #N/A | #N/A | #N/A | #N/A | #N/A |
| H8GMG9 | ABC-type transport system involved in resistance to organic solvents, ATPase component OS=Methylomicrobium album BG8 OX=686340 GN=Metal_3013 PE=4 SV=1 | 1 | 1 | 1.93 | 1 | 1 | 1.92 | #N/A | #N/A | #N/A |
| H8GG77 | Putative stress response protein, TerZ-and CABP1 OS=Methylomicrobium album BG8 OX=686340 GN=Metal_2261 PE=4 SV=1 | 1 | 1 | 1.91 | 2 | 1 | 4.55 | #N/A | #N/A | #N/A |
| *H8GQ61* | *Uncharacterized protein OS=Methylomicrobium album BG8 OX=686340 GN=Metal_3338 PE=4 SV=1* | 1 | 1 | 2.05 | 7 | 6 | 17.76 | 1 | 1 | 1.99 |
| H8GFX2 | Bifunctional ligase/repressor BirA OS=Methylomicrobium album BG8 OX=686340 GN=birA PE=3 SV=1 | 1 | 1 | 2.1 | #N/A | #N/A | #N/A | #N/A | #N/A | #N/A |
| H8GGS6 | Cation/multidrug efflux pump OS=Methylomicrobium album BG8 OX=686340 GN=Metal_2301 PE=4 SV=1 | 1 | 1 | 1.93 | #N/A | #N/A | #N/A | #N/A | #N/A | #N/A |
| H8GFZ0 | Efflux transporter, outer membrane factor lipoprotein, NodT family OS=Methylomicrobium album BG8 OX=686340 GN=Metal_0918 PE=3 SV=1 | 1 | 1 | 2.17 | #N/A | #N/A | #N/A | 1 | 1 | 2.3 |
| H8GNC0 | Uncharacterized protein OS=Methylomicrobium album BG8 OX=686340 GN=Metal_0498 PE=4 SV=1 | 1 | 1 | 2.27 | #N/A | #N/A | #N/A | #N/A | #N/A | #N/A |
| H8GMW7 | Response regulator with CheY-like receiver, AAA-type ATPase, and DNA-binding domains OS=Methylomicrobium album BG8 OX=686340 GN=Metal_1750 PE=4 SV=1 | #N/A | #N/A | #N/A | 4 | 1 | 8.36 | 3 | 1 | 6.63 |
| *H8GI56* | *Uncharacterized protein OS=Methylomicrobium album BG8 OX=686340 GN=Metal_2481 PE=4 SV=1* | #N/A | #N/A | #N/A | 4 | 4 | 9.84 | 1 | 1 | 2.73 |
| H8GK39 | Phosphomannomutase OS=Methylomicrobium album BG8 OX=686340 GN=Metal_1376 PE=3 SV=1 | #N/A | #N/A | #N/A | 3 | 2 | 6.35 | #N/A | #N/A | #N/A |
| H8GNN9 | Site-specific DNA-methyltransferase (adenine-specific) OS=Methylomicrobium album BG8 OX=686340 GN=Metal_3119 PE=3 SV=1 | #N/A | #N/A | #N/A | 3 | 2 | 6.16 | 1 | 1 | 2.08 |
| H8GNH6 | Sulfate adenylyltransferase subunit 2 OS=Methylomicrobium album BG8 OX=686340 GN=cysD PE=3 SV=1 | #N/A | #N/A | #N/A | 3 | 1 | 6.57 | #N/A | #N/A | #N/A |
| H8GJK1 | Outer membrane receptor protein OS=Methylomicrobium album BG8 OX=686340 GN=Metal_2650 PE=3 SV=1 | #N/A | #N/A | #N/A | 3 | 2 | 9.75 | #N/A | #N/A | #N/A |
| H8GJK6 | Bifunctional NAD(P)H-hydrate repair enzyme OS=Methylomicrobium album BG8 OX=686340 GN=nnrD PE=3 SV=1 | #N/A | #N/A | #N/A | 3 | 1 | 8.13 | #N/A | #N/A | #N/A |
| H8GQF7 | Putative metal-dependent hydrolase OS=Methylomicrobium album BG8 OX=686340 GN=Metal_2023 PE=4 SV=1 | #N/A | #N/A | #N/A | 3 | 3 | 6.54 | 2 | 2 | 4.99 |
| H8GJ52 | Cobyric acid synthase OS=Methylomicrobium album BG8 OX=686340 GN=cobQ PE=3 SV=1 | #N/A | #N/A | #N/A | 3 | 2 | 6.8 | #N/A | #N/A | #N/A |
| H8GQ31 | 50S ribosomal protein L14 OS=Methylomicrobium album BG8 OX=686340 GN=rplN PE=3 SV=1 | #N/A | #N/A | #N/A | 2 | 1 | 5.69 | #N/A | #N/A | #N/A |
| H8GHQ5 | RecBCD enzyme subunit RecD OS=Methylomicrobium album BG8 OX=686340 GN=recD PE=3 SV=1 | #N/A | #N/A | #N/A | 2 | 1 | 3.91 | #N/A | #N/A | #N/A |
| H8GIS0 | ATP synthase, F1 beta subunit OS=Methylomicrobium album BG8 OX=686340 GN=Metal_2543 PE=3 SV=1 | #N/A | #N/A | #N/A | 2 | 1 | 5.66 | #N/A | #N/A | #N/A |
| H8GPM4 | Putative phage-type endonuclease OS=Methylomicrobium album BG8 OX=686340 GN=Metal_0643 PE=4 SV=1 | #N/A | #N/A | #N/A | 2 | 1 | 4.18 | #N/A | #N/A | #N/A |
| H8GI94 | Death-on-curing family protein OS=Methylomicrobium album BG8 OX=686340 GN=Metal_3762 PE=4 SV=1 | #N/A | #N/A | #N/A | 2 | 1 | 4.69 | 3 | 1 | 6.02 |
| *H8GH59* | *PAS domain S-box/diguanylate cyclase (GGDEF) domain-containing protein OS=Methylomicrobium album BG8 OX=686340 GN=Metal_1031 PE=4 SV=1* | #N/A | #N/A | #N/A | 2 | 1 | 5.01 | 5 | 1 | 13.31 |
| H8GGU1 | Uncharacterized protein OS=Methylomicrobium album BG8 OX=686340 GN=Metal_2318 PE=4 SV=1 | #N/A | #N/A | #N/A | 2 | 2 | 7.39 | #N/A | #N/A | #N/A |
| H8GL30 | Peptidoglycan-associated protein OS=Methylomicrobium album BG8 OX=686340 GN=pal PE=3 SV=1 | #N/A | #N/A | #N/A | 2 | 2 | 4.84 | #N/A | #N/A | #N/A |
| H8GR09 | Tetratricopeptide repeat protein OS=Methylomicrobium album BG8 OX=686340 GN=Metal_0839 PE=4 SV=1 | #N/A | #N/A | #N/A | 2 | 1 | 6.38 | 1 | 1 | 2.49 |
| H8GGQ1 | Uncharacterized protein involved in outer membrane biogenesis OS=Methylomicrobium album BG8 OX=686340 GN=Metal_2274 PE=4 SV=1 | #N/A | #N/A | #N/A | 2 | 2 | 3.85 | 1 | 1 | 2.38 |
| H8GJ24 | REJ domain protein OS=Methylomicrobium album BG8 OX=686340 GN=Metal_3894 PE=4 SV=1 | #N/A | #N/A | #N/A | 2 | 2 | 5.51 | #N/A | #N/A | #N/A |
| H8GN53 | Rhs element Vgr protein OS=Methylomicrobium album BG8 OX=686340 GN=Metal_3088 PE=3 SV=1 | #N/A | #N/A | #N/A | 2 | 1 | 4.2 | 1 | 1 | 1.93 |
| H8GNY3 | Putative TIM-barrel fold metal-dependent hydrolase OS=Methylomicrobium album BG8 OX=686340 GN=Metal_0555 PE=4 SV=1 | #N/A | #N/A | #N/A | 2 | 1 | 4.7 | 1 | 1 | 2.62 |
| H8GNN3 | Uncharacterized protein OS=Methylomicrobium album BG8 OX=686340 GN=Metal_3113 PE=4 SV=1 | #N/A | #N/A | #N/A | 2 | 2 | 8.27 | #N/A | #N/A | #N/A |
| H8GH93 | Periplasmic serine endoprotease DegP-like OS=Methylomicrobium album BG8 OX=686340 GN=Metal_1066 PE=3 SV=1 | #N/A | #N/A | #N/A | 2 | 1 | 6.1 | #N/A | #N/A | #N/A |
| H8GR73 | Carbamoyl-phosphate synthase large chain OS=Methylomicrobium album BG8 OX=686340 GN=carB PE=3 SV=1 | #N/A | #N/A | #N/A | 2 | 1 | 3.96 | #N/A | #N/A | #N/A |
| H8GKF8 | HAMP domain-containing protein,cache domain-containing protein OS=Methylomicrobium album BG8 OX=686340 GN=Metal_2751 PE=4 SV=1 | #N/A | #N/A | #N/A | 2 | 1 | 4.81 | #N/A | #N/A | #N/A |
| H8GID7 | Uncharacterized protein OS=Methylomicrobium album BG8 OX=686340 GN=Metal_3807 PE=4 SV=1 | #N/A | #N/A | #N/A | 2 | 1 | 4.08 | 1 | 1 | 2.06 |
| H8GMR8 | Probable chorismate pyruvate-lyase OS=Methylomicrobium album BG8 OX=686340 GN=ubiC PE=3 SV=1 | #N/A | #N/A | #N/A | 2 | 1 | 5.51 | #N/A | #N/A | #N/A |
| H8GIA3 | ATP synthase subunit alpha OS=Methylomicrobium album BG8 OX=686340 GN=atpA PE=3 SV=1 | #N/A | #N/A | #N/A | 2 | 1 | 6.59 | 1 | 1 | 2.08 |
| *H8GI24* | *Uncharacterized protein OS=Methylomicrobium album BG8 OX=686340 GN=Metal_2446 PE=4 SV=1* | #N/A | #N/A | #N/A | 2 | 1 | 5.39 | 6 | 1 | 15.75 |
| H8GMP6 | Uncharacterized protein OS=Methylomicrobium album BG8 OX=686340 GN=Metal_0432 PE=4 SV=1 | #N/A | #N/A | #N/A | 1 | 1 | 1.92 | #N/A | #N/A | #N/A |
| H8GLR3 | Uncharacterized protein OS=Methylomicrobium album BG8 OX=686340 GN=Metal_2908 PE=4 SV=1 | #N/A | #N/A | #N/A | 1 | 1 | 1.95 | #N/A | #N/A | #N/A |
| H8GQN6 | Multifunctional CCA protein OS=Methylomicrobium album BG8 OX=686340 GN=cca PE=3 SV=1 | #N/A | #N/A | #N/A | 1 | 1 | 1.93 | #N/A | #N/A | #N/A |
| H8GM23 | ABC transporter, substrate binding protein, PQQ-dependent alcohol dehydrogenase system OS=Methylomicrobium album BG8 OX=686340 GN=Metal_1605 PE=4 SV=1 | #N/A | #N/A | #N/A | 1 | 1 | 1.94 | #N/A | #N/A | #N/A |
| H8GMH4 | Uncharacterized protein OS=Methylomicrobium album BG8 OX=686340 GN=Metal_3018 PE=4 SV=1 | #N/A | #N/A | #N/A | 1 | 1 | 2.82 | #N/A | #N/A | #N/A |
| H8GQN1 | Cell division protein OS=Methylomicrobium album BG8 OX=686340 GN=Metal_3353 PE=4 SV=1 | #N/A | #N/A | #N/A | 1 | 1 | 1.94 | #N/A | #N/A | #N/A |
| H8GIT2 | EAL domain-containing protein OS=Methylomicrobium album BG8 OX=686340 GN=Metal_2555 PE=4 SV=1 | #N/A | #N/A | #N/A | 1 | 1 | 2.31 | #N/A | #N/A | #N/A |
| H8GK84 | Uncharacterized protein OS=Methylomicrobium album BG8 OX=686340 GN=Metal_1423 PE=4 SV=1 | #N/A | #N/A | #N/A | 1 | 1 | 2.32 | #N/A | #N/A | #N/A |
| H8GJJ4 | Transposase OS=Methylomicrobium album BG8 OX=686340 GN=Metal_0719 PE=4 SV=1 | #N/A | #N/A | #N/A | 1 | 1 | 1.95 | #N/A | #N/A | #N/A |
| H8GM09 | Uncharacterized protein OS=Methylomicrobium album BG8 OX=686340 GN=Metal_0347 PE=4 SV=1 | #N/A | #N/A | #N/A | 1 | 1 | 2.53 | #N/A | #N/A | #N/A |
| H8GII0 | TonB family protein OS=Methylomicrobium album BG8 OX=686340 GN=Metal_1196 PE=3 SV=1 | #N/A | #N/A | #N/A | 1 | 1 | 2.06 | #N/A | #N/A | #N/A |
| H8GPS3 | Dual-specificity RNA methyltransferase RlmN OS=Methylomicrobium album BG8 OX=686340 GN=rlmN PE=3 SV=1 | #N/A | #N/A | #N/A | 1 | 1 | 2.21 | 1 | 1 | 2.41 |
| H8GLK6 | CRISPR-associated protein Cas7/Cse4/CasC, subtype I-E OS=Methylomicrobium album BG8 OX=686340 GN=Metal_1591 PE=4 SV=1 | #N/A | #N/A | #N/A | 1 | 1 | 2.43 | #N/A | #N/A | #N/A |
| H8GMH5 | Uncharacterized protein OS=Methylomicrobium album BG8 OX=686340 GN=Metal_3019 PE=4 SV=1 | #N/A | #N/A | #N/A | 1 | 1 | 2.37 | #N/A | #N/A | #N/A |
| H8GMF5 | Parvulin-like peptidyl-prolyl isomerase OS=Methylomicrobium album BG8 OX=686340 GN=Metal_2999 PE=4 SV=1 | #N/A | #N/A | #N/A | 1 | 1 | 4.06 | #N/A | #N/A | #N/A |
| H8GQE7 | Cytochrome c domain-containing protein OS=Methylomicrobium album BG8 OX=686340 GN=Metal_0773 PE=4 SV=1 | #N/A | #N/A | #N/A | 1 | 1 | 2.33 | #N/A | #N/A | #N/A |
| H8GNP5 | Type I restriction enzyme R Protein OS=Methylomicrobium album BG8 OX=686340 GN=Metal_3125 PE=3 SV=1 | #N/A | #N/A | #N/A | 1 | 1 | 1.97 | 2 | 1 | 3.94 |
| H8GGA9 | Uncharacterized protein OS=Methylomicrobium album BG8 OX=686340 GN=Metal_3541 PE=4 SV=1 | #N/A | #N/A | #N/A | 1 | 1 | 2.4 | #N/A | #N/A | #N/A |
| H8GN39 | MuF_C domain-containing protein OS=Methylomicrobium album BG8 OX=686340 GN=Metal_3073 PE=4 SV=1 | #N/A | #N/A | #N/A | 1 | 1 | 2.02 | #N/A | #N/A | #N/A |
| H8GIJ6 | Peptide methionine sulfoxide reductase MsrA OS=Methylomicrobium album BG8 OX=686340 GN=msrA PE=3 SV=1 | #N/A | #N/A | #N/A | 1 | 1 | 1.92 | #N/A | #N/A | #N/A |
| H8GLH8 | Uncharacterized protein OS=Methylomicrobium album BG8 OX=686340 GN=Metal_1560 PE=4 SV=1 | #N/A | #N/A | #N/A | 1 | 1 | 1.97 | #N/A | #N/A | #N/A |
| H8GPP4 | Stress-induced acidophilic repeat motif-containing protein OS=Methylomicrobium album BG8 OX=686340 GN=Metal_0664 PE=4 SV=1 | #N/A | #N/A | #N/A | 1 | 1 | 2.47 | 1 | 1 | 2.61 |
| H8GM37 | 3-oxoacyl-[acyl-carrier-protein] reductase OS=Methylomicrobium album BG8 OX=686340 GN=Metal_1619 PE=3 SV=1 | #N/A | #N/A | #N/A | 1 | 1 | 2.07 | #N/A | #N/A | #N/A |
| H8GIR1 | GBBH-like_N domain-containing protein OS=Methylomicrobium album BG8 OX=686340 GN=Metal_2534 PE=4 SV=1 | #N/A | #N/A | #N/A | 1 | 1 | 2.55 | #N/A | #N/A | #N/A |
| H8GHC9 | Translation initiation factor IF-2 OS=Methylomicrobium album BG8 OX=686340 GN=infB PE=3 SV=1 | #N/A | #N/A | #N/A | 1 | 1 | 2.33 | #N/A | #N/A | #N/A |
| H8GQG9 | Dihydroxy-acid dehydratase OS=Methylomicrobium album BG8 OX=686340 GN=ilvD PE=3 SV=1 | #N/A | #N/A | #N/A | 1 | 1 | 2.01 | #N/A | #N/A | #N/A |
| H8GPN4 | Uncharacterized protein OS=Methylomicrobium album BG8 OX=686340 GN=Metal_0653 PE=4 SV=1 | #N/A | #N/A | #N/A | 1 | 1 | 2.31 | #N/A | #N/A | #N/A |
| H8GNL3 | RHS repeat-associated core domain protein OS=Methylomicrobium album BG8 OX=686340 GN=Metal_1840 PE=4 SV=1 | #N/A | #N/A | #N/A | 1 | 1 | 2.6 | #N/A | #N/A | #N/A |
| H8GK23 | Uncharacterized protein OS=Methylomicrobium album BG8 OX=686340 GN=Metal_0113 PE=3 SV=1 | #N/A | #N/A | #N/A | 1 | 1 | 2.29 | #N/A | #N/A | #N/A |
| H8GGF3 | EAL domain-containing protein OS=Methylomicrobium album BG8 OX=686340 GN=Metal_3585 PE=4 SV=1 | #N/A | #N/A | #N/A | 1 | 1 | 2.46 | #N/A | #N/A | #N/A |
| H8GQJ9 | Uncharacterized protein OS=Methylomicrobium album BG8 OX=686340 GN=Metal_2068 PE=4 SV=1 | #N/A | #N/A | #N/A | 1 | 1 | 2.67 | 1 | 1 | 2.69 |
| H8GIG2 | Type II secretory pathway, component PulK OS=Methylomicrobium album BG8 OX=686340 GN=Metal_3832 PE=3 SV=1 | #N/A | #N/A | #N/A | 1 | 1 | 2.39 | #N/A | #N/A | #N/A |
| H8GNE5 | 3-isopropylmalate dehydrogenase OS=Methylomicrobium album BG8 OX=686340 GN=Metal_0524 PE=3 SV=1 | #N/A | #N/A | #N/A | 1 | 1 | 2.41 | #N/A | #N/A | #N/A |
| H8GI84 | Valine--tRNA ligase OS=Methylomicrobium album BG8 OX=686340 GN=valS PE=3 SV=1 | #N/A | #N/A | #N/A | 1 | 1 | 1.98 | 1 | 1 | 1.96 |
| H8GNJ7 | RND family efflux transporter, MFP subunit OS=Methylomicrobium album BG8 OX=686340 GN=Metal_1823 PE=3 SV=1 | #N/A | #N/A | #N/A | 1 | 1 | 2.6 | #N/A | #N/A | #N/A |
| H8GH97 | Glutamyl-tRNA reductase OS=Methylomicrobium album BG8 OX=686340 GN=Metal_1070 PE=4 SV=1 | #N/A | #N/A | #N/A | 1 | 1 | 2 | #N/A | #N/A | #N/A |
| H8GHI0 | Elongation factor Ts OS=Methylomicrobium album BG8 OX=686340 GN=tsf PE=3 SV=1 | #N/A | #N/A | #N/A | 1 | 1 | 1.93 | #N/A | #N/A | #N/A |
| H8GQT5 | ABC-type nitrate/sulfonate/bicarbonate transport system, ATPase component OS=Methylomicrobium album BG8 OX=686340 GN=Metal_3417 PE=4 SV=1 | #N/A | #N/A | #N/A | 1 | 1 | 2.43 | #N/A | #N/A | #N/A |
| H8GNH8 | FAD-dependent dehydrogenase OS=Methylomicrobium album BG8 OX=686340 GN=Metal_1804 PE=4 SV=1 | #N/A | #N/A | #N/A | 1 | 1 | 3.15 | #N/A | #N/A | #N/A |
| H8GJ39 | Acyl-CoA dehydrogenase OS=Methylomicrobium album BG8 OX=686340 GN=Metal_3909 PE=3 SV=1 | #N/A | #N/A | #N/A | 1 | 1 | 2.09 | 1 | 1 | 2.59 |
| H8GL38 | Aminodeoxychorismate synthase OS=Methylomicrobium album BG8 OX=686340 GN=Metal_2828 PE=3 SV=1 | #N/A | #N/A | #N/A | 1 | 1 | 1.95 | #N/A | #N/A | #N/A |
| H8GNM3 | TPR_REGION domain-containing protein OS=Methylomicrobium album BG8 OX=686340 GN=Metal_1850 PE=4 SV=1 | #N/A | #N/A | #N/A | 1 | 1 | 2.04 | #N/A | #N/A | #N/A |
| H8GHM2 | Uncharacterized protein OS=Methylomicrobium album BG8 OX=686340 GN=Metal_3693 PE=4 SV=1 | #N/A | #N/A | #N/A | 1 | 1 | 2.73 | 1 | 1 | 2.4 |
| H8GLU9 | Glyceraldehyde-3-phosphate dehydrogenase OS=Methylomicrobium album BG8 OX=686340 GN=Metal_0282 PE=3 SV=1 | #N/A | #N/A | #N/A | 1 | 1 | 2.68 | #N/A | #N/A | #N/A |
| H8GIU9 | 2-isopropylmalate synthase OS=Methylomicrobium album BG8 OX=686340 GN=leuA PE=3 SV=1 | #N/A | #N/A | #N/A | 1 | 1 | 2.06 | #N/A | #N/A | #N/A |
| H8GNA2 | OMP_b-brl domain-containing protein OS=Methylomicrobium album BG8 OX=686340 GN=Metal_0480 PE=4 SV=1 | #N/A | #N/A | #N/A | 1 | 1 | 3.12 | #N/A | #N/A | #N/A |
| H8GI30 | Histidine kinase OS=Methylomicrobium album BG8 OX=686340 GN=Metal_2453 PE=4 SV=1 | #N/A | #N/A | #N/A | 1 | 1 | 2 | #N/A | #N/A | #N/A |
| H8GKK2 | Outer membrane protein OS=Methylomicrobium album BG8 OX=686340 GN=Metal_0142 PE=3 SV=1 | #N/A | #N/A | #N/A | 1 | 1 | 2.24 | #N/A | #N/A | #N/A |
| H8GIW8 | DNA (cytosine-5-)-methyltransferase OS=Methylomicrobium album BG8 OX=686340 GN=Metal_2593 PE=4 SV=1 | #N/A | #N/A | #N/A | 1 | 1 | 2.39 | 1 | 1 | 2.04 |
| H8GN52 | Type VI secretion lipoprotein, VC_A0113 family OS=Methylomicrobium album BG8 OX=686340 GN=Metal_3087 PE=4 SV=1 | #N/A | #N/A | #N/A | 1 | 1 | 2.81 | #N/A | #N/A | #N/A |
| H8GM59 | Uncharacterized protein OS=Methylomicrobium album BG8 OX=686340 GN=Metal_1641 PE=4 SV=1 | #N/A | #N/A | #N/A | 1 | 1 | 2.3 | 1 | 1 | 1.96 |
| H8GR31 | Cation/multidrug efflux pump OS=Methylomicrobium album BG8 OX=686340 GN=Metal_2103 PE=4 SV=1 | #N/A | #N/A | #N/A | 1 | 1 | 2.17 | #N/A | #N/A | #N/A |
| H8GQ38 | 50S ribosomal protein L2 OS=Methylomicrobium album BG8 OX=686340 GN=rplB PE=3 SV=1 | #N/A | #N/A | #N/A | 1 | 1 | 2.85 | #N/A | #N/A | #N/A |
| H8GLD8 | LysM domain-containing protein OS=Methylomicrobium album BG8 OX=686340 GN=Metal_0274 PE=4 SV=1 | #N/A | #N/A | #N/A | 1 | 1 | 2.61 | #N/A | #N/A | #N/A |
| H8GIL3 | 5-bromo-4-chloroindolyl phosphate hydrolysis protein OS=Methylomicrobium album BG8 OX=686340 GN=Metal_1234 PE=4 SV=1 | #N/A | #N/A | #N/A | 1 | 1 | 1.92 | 1 | 1 | 2 |
| H8GLL2 | DUF2384 domain-containing protein OS=Methylomicrobium album BG8 OX=686340 GN=Metal_1597 PE=4 SV=1 | #N/A | #N/A | #N/A | 1 | 1 | 2.28 | 1 | 1 | 2.03 |
| H8GGZ2 | Probable septum site-determining protein MinC OS=Methylomicrobium album BG8 OX=686340 GN=minC PE=3 SV=1 | #N/A | #N/A | #N/A | 1 | 1 | 1.91 | #N/A | #N/A | #N/A |
| H8GQD0 | Uncharacterized protein OS=Methylomicrobium album BG8 OX=686340 GN=Metal_0756 PE=4 SV=1 | #N/A | #N/A | #N/A | 1 | 1 | 1.98 | #N/A | #N/A | #N/A |
| H8GR60 | Uncharacterized protein OS=Methylomicrobium album BG8 OX=686340 GN=Metal_2133 PE=4 SV=1 | #N/A | #N/A | #N/A | 1 | 1 | 3.01 | #N/A | #N/A | #N/A |
| H8GLS4 | Protein HflK OS=Methylomicrobium album BG8 OX=686340 GN=Metal_2920 PE=3 SV=1 | #N/A | #N/A | #N/A | 1 | 1 | 3.03 | #N/A | #N/A | #N/A |
| H8GNE6 | Uncharacterized protein OS=Methylomicrobium album BG8 OX=686340 GN=Metal_0525 PE=4 SV=1 | #N/A | #N/A | #N/A | 1 | 1 | 2.34 | #N/A | #N/A | #N/A |
| H8GG30 | Response regulator with CheY-like receiver, AAA-type ATPase, and DNA-binding domains OS=Methylomicrobium album BG8 OX=686340 GN=Metal_2204 PE=4 SV=1 | #N/A | #N/A | #N/A | 1 | 1 | 2.01 | #N/A | #N/A | #N/A |
| H8GKU5 | Heavy metal efflux pump, cobalt-zinc-cadmium OS=Methylomicrobium album BG8 OX=686340 GN=Metal_1482 PE=3 SV=1 | #N/A | #N/A | #N/A | 1 | 1 | 2.07 | #N/A | #N/A | #N/A |
| H8GLC0 | Diguanylate cyclase (GGDEF) domain-containing protein OS=Methylomicrobium album BG8 OX=686340 GN=Metal_0256 PE=4 SV=1 | #N/A | #N/A | #N/A | 1 | 1 | 2.15 | 1 | 1 | 2.56 |
| H8GGM3 | CDP-diacylglycerol--glycerol-3-phosphate 3-phosphatidyltransferase OS=Methylomicrobium album BG8 OX=686340 GN=Metal_1000 PE=3 SV=1 | #N/A | #N/A | #N/A | 1 | 1 | 1.95 | #N/A | #N/A | #N/A |
| H8GKL5 | Uncharacterized protein OS=Methylomicrobium album BG8 OX=686340 GN=Metal_0156 PE=4 SV=1 | #N/A | #N/A | #N/A | 1 | 1 | 2.19 | #N/A | #N/A | #N/A |
| H8GLG0 | DNA repair protein RecO OS=Methylomicrobium album BG8 OX=686340 GN=recO PE=3 SV=1 | #N/A | #N/A | #N/A | 1 | 1 | 2.3 | #N/A | #N/A | #N/A |
| H8GJW4 | tRNA-dihydrouridine synthase B OS=Methylomicrobium album BG8 OX=686340 GN=dusB PE=3 SV=1 | #N/A | #N/A | #N/A | 1 | 1 | 2.45 | #N/A | #N/A | #N/A |
| H8GGS7 | RND family efflux transporter, MFP subunit OS=Methylomicrobium album BG8 OX=686340 GN=Metal_2302 PE=3 SV=1 | #N/A | #N/A | #N/A | 1 | 1 | 1.97 | #N/A | #N/A | #N/A |
| H8GMS3 | Phosphoglycerate kinase OS=Methylomicrobium album BG8 OX=686340 GN=pgk PE=3 SV=1 | #N/A | #N/A | #N/A | 1 | 1 | 2.32 | #N/A | #N/A | #N/A |
| H8GMX5 | Methyl-accepting chemotaxis protein OS=Methylomicrobium album BG8 OX=686340 GN=Metal_1758 PE=4 SV=1 | #N/A | #N/A | #N/A | 1 | 1 | 2.15 | #N/A | #N/A | #N/A |
| H8GPS7 | Outer membrane protein assembly factor BamB OS=Methylomicrobium album BG8 OX=686340 GN=bamB PE=3 SV=1 | #N/A | #N/A | #N/A | 1 | 1 | 2.4 | #N/A | #N/A | #N/A |
| H8GJ35 | Sigma-54 interacting regulator OS=Methylomicrobium album BG8 OX=686340 GN=Metal_3905 PE=4 SV=1 | #N/A | #N/A | #N/A | 1 | 1 | 2.61 | #N/A | #N/A | #N/A |
| H8GI74 | Arginine biosynthesis bifunctional protein ArgJ OS=Methylomicrobium album BG8 OX=686340 GN=argJ PE=3 SV=1 | #N/A | #N/A | #N/A | 1 | 1 | 2.94 | #N/A | #N/A | #N/A |
| H8GKT3 | Methylase involved in ubiquinone/menaquinone biosynthesis OS=Methylomicrobium album BG8 OX=686340 GN=Metal_1470 PE=4 SV=1 | #N/A | #N/A | #N/A | 1 | 1 | 2.1 | #N/A | #N/A | #N/A |
| H8GQH4 | Acyl-CoA synthetase (NDP forming) OS=Methylomicrobium album BG8 OX=686340 GN=Metal_2042 PE=4 SV=1 | #N/A | #N/A | #N/A | 1 | 1 | 2.81 | 1 | 1 | 2.47 |
| H8GQA6 | 56kDa selenium binding protein (SBP56) OS=Methylomicrobium album BG8 OX=686340 GN=Metal_0731 PE=3 SV=1 | #N/A | #N/A | #N/A | 1 | 1 | 2.41 | #N/A | #N/A | #N/A |
| H8GMI8 | DNA polymerase I OS=Methylomicrobium album BG8 OX=686340 GN=polA PE=3 SV=1 | #N/A | #N/A | #N/A | 1 | 1 | 2.32 | 2 | 1 | 4.95 |
| H8GLR8 | Major facilitator superfamily permease OS=Methylomicrobium album BG8 OX=686340 GN=Metal_2914 PE=4 SV=1 | #N/A | #N/A | #N/A | 1 | 1 | 2.66 | #N/A | #N/A | #N/A |
| H8GGS9 | Non-ribosomal peptide synthase/amino acid adenylation enzyme OS=Methylomicrobium album BG8 OX=686340 GN=Metal_2304 PE=4 SV=1 | #N/A | #N/A | #N/A | 1 | 1 | 2.08 | 1 | 1 | 2.22 |
| H8GP63 | Single-stranded-DNA-specific exonuclease RecJ OS=Methylomicrobium album BG8 OX=686340 GN=Metal_1883 PE=3 SV=1 | #N/A | #N/A | #N/A | 1 | 1 | 2 | #N/A | #N/A | #N/A |
| H8GNR8 | Metalloendopeptidase-like membrane protein OS=Methylomicrobium album BG8 OX=686340 GN=Metal_3150 PE=4 SV=1 | #N/A | #N/A | #N/A | 1 | 1 | 2.02 | #N/A | #N/A | #N/A |
| H8GPC6 | Site-specific DNA-methyltransferase (adenine-specific) OS=Methylomicrobium album BG8 OX=686340 GN=Metal_3199 PE=3 SV=1 | #N/A | #N/A | #N/A | 1 | 1 | 1.99 | #N/A | #N/A | #N/A |
| H8GJ90 | Amino acid adenylation enzyme/thioester reductase family protein OS=Methylomicrobium album BG8 OX=686340 GN=Metal_1280 PE=4 SV=1 | #N/A | #N/A | #N/A | 1 | 1 | 2.44 | #N/A | #N/A | #N/A |
| H8GK75 | Sulfite reductase, alpha subunit (Flavoprotein) OS=Methylomicrobium album BG8 OX=686340 GN=Metal_1414 PE=4 SV=1 | #N/A | #N/A | #N/A | 1 | 1 | 2.39 | #N/A | #N/A | #N/A |
| H8GPU9 | UPF0753 protein Metal_1963 OS=Methylomicrobium album BG8 OX=686340 GN=Metal_1963 PE=3 SV=1 | #N/A | #N/A | #N/A | 1 | 1 | 2.4 | #N/A | #N/A | #N/A |
| H8GQA8 | GMP synthase [glutamine-hydrolyzing] OS=Methylomicrobium album BG8 OX=686340 GN=guaA PE=3 SV=1 | #N/A | #N/A | #N/A | 1 | 1 | 1.96 | #N/A | #N/A | #N/A |
| H8GIM0 | Transcriptional regulator OS=Methylomicrobium album BG8 OX=686340 GN=Metal_1242 PE=3 SV=1 | #N/A | #N/A | #N/A | 1 | 1 | 2.22 | #N/A | #N/A | #N/A |
| H8GRL6 | Nucleotidyltransferase/DNA polymerase involved in DNA repair OS=Methylomicrobium album BG8 OX=686340 GN=Metal_4028 PE=3 SV=1 | #N/A | #N/A | #N/A | 1 | 1 | 1.93 | #N/A | #N/A | #N/A |
| H8GLG6 | Uncharacterized protein OS=Methylomicrobium album BG8 OX=686340 GN=Metal_1548 PE=4 SV=1 | #N/A | #N/A | #N/A | 1 | 1 | 2.73 | #N/A | #N/A | #N/A |
| H8GN57 | Aspzincin_M35 domain-containing protein OS=Methylomicrobium album BG8 OX=686340 GN=Metal_3092 PE=3 SV=1 | #N/A | #N/A | #N/A | 1 | 1 | 1.93 | 2 | 2 | 4.31 |
| H8GPX4 | Cytochrome c peroxidase OS=Methylomicrobium album BG8 OX=686340 GN=Metal_1989 PE=4 SV=1 | #N/A | #N/A | #N/A | 1 | 1 | 2.54 | 1 | 1 | 2.51 |
| H8GPG7 | P-type Ca(2+) transporter OS=Methylomicrobium album BG8 OX=686340 GN=Metal_3243 PE=4 SV=1 | #N/A | #N/A | #N/A | 1 | 1 | 2.19 | #N/A | #N/A | #N/A |
| H8GPJ4 | Bacterial conjugation TrbI-like protein OS=Methylomicrobium album BG8 OX=686340 GN=Metal_0611 PE=4 SV=1 | #N/A | #N/A | #N/A | 1 | 1 | 2.14 | #N/A | #N/A | #N/A |
| H8GL42 | Site-specific DNA-methyltransferase (adenine-specific) OS=Methylomicrobium album BG8 OX=686340 GN=Metal_2832 PE=3 SV=1 | #N/A | #N/A | #N/A | 1 | 1 | 2.44 | #N/A | #N/A | #N/A |
| H8GRC9 | Fructose-bisphosphate aldolase OS=Methylomicrobium album BG8 OX=686340 GN=Metal_3459 PE=3 SV=1 | #N/A | #N/A | #N/A | 1 | 1 | 4.19 | 1 | 1 | 3.58 |
| H8GIB0 | Heavy metal efflux pump, cobalt-zinc-cadmium OS=Methylomicrobium album BG8 OX=686340 GN=Metal_3778 PE=3 SV=1 | #N/A | #N/A | #N/A | 1 | 1 | 1.95 | #N/A | #N/A | #N/A |
| H8GJJ3 | Adenylate kinase OS=Methylomicrobium album BG8 OX=686340 GN=adk PE=3 SV=1 | #N/A | #N/A | #N/A | 1 | 1 | 2.26 | #N/A | #N/A | #N/A |
| H8GL62 | Surface lipoprotein OS=Methylomicrobium album BG8 OX=686340 GN=Metal_2856 PE=3 SV=1 | #N/A | #N/A | #N/A | 1 | 1 | 2.57 | #N/A | #N/A | #N/A |
| H8GR02 | Homoserine O-succinyltransferase OS=Methylomicrobium album BG8 OX=686340 GN=metAS PE=3 SV=1 | #N/A | #N/A | #N/A | 1 | 1 | 2.09 | #N/A | #N/A | #N/A |
| H8GMH3 | Putative divalent heavy-metal cations transporter OS=Methylomicrobium album BG8 OX=686340 GN=Metal_3017 PE=4 SV=1 | #N/A | #N/A | #N/A | 1 | 1 | 2.36 | #N/A | #N/A | #N/A |
| H8GMV1 | Regulator of stationary/sporulation gene expression OS=Methylomicrobium album BG8 OX=686340 GN=Metal_1734 PE=4 SV=1 | #N/A | #N/A | #N/A | 1 | 1 | 2.04 | #N/A | #N/A | #N/A |
| H8GRB9 | Response regulator with CheY-like receiver domain and winged-helix DNA-binding domain OS=Methylomicrobium album BG8 OX=686340 GN=Metal_3448 PE=4 SV=1 | #N/A | #N/A | #N/A | 1 | 1 | 2.45 | #N/A | #N/A | #N/A |
| H8GQX9 | Phosphate regulon transcriptional regulatory protein PhoB OS=Methylomicrobium album BG8 OX=686340 GN=Metal_0808 PE=4 SV=1 | #N/A | #N/A | #N/A | 1 | 1 | 2.51 | #N/A | #N/A | #N/A |
| H8GH68 | Putative ATPase OS=Methylomicrobium album BG8 OX=686340 GN=Metal_1040 PE=4 SV=1 | #N/A | #N/A | #N/A | 1 | 1 | 2.02 | #N/A | #N/A | #N/A |
| H8GI40 | Uncharacterized protein OS=Methylomicrobium album BG8 OX=686340 GN=Metal_2464 PE=4 SV=1 | #N/A | #N/A | #N/A | 1 | 1 | 1.99 | #N/A | #N/A | #N/A |
| H8GHU1 | Putative K(+)-stimulated pyrophosphate-energized sodium pump OS=Methylomicrobium album BG8 OX=686340 GN=hppA PE=3 SV=1 | #N/A | #N/A | #N/A | 1 | 1 | 2.33 | #N/A | #N/A | #N/A |
| H8GKA7 | YkuD domain-containing protein OS=Methylomicrobium album BG8 OX=686340 GN=Metal_2692 PE=3 SV=1 | #N/A | #N/A | #N/A | 1 | 1 | 2.45 | #N/A | #N/A | #N/A |
| H8GQN2 | Arginine--tRNA ligase OS=Methylomicrobium album BG8 OX=686340 GN=argS PE=3 SV=1 | #N/A | #N/A | #N/A | 1 | 1 | 2.62 | #N/A | #N/A | #N/A |
| H8GL01 | Phosphate-starvation-inducible E OS=Methylomicrobium album BG8 OX=686340 GN=Metal_2790 PE=4 SV=1 | #N/A | #N/A | #N/A | 1 | 1 | 2.01 | #N/A | #N/A | #N/A |
| H8GH27 | Putative periplasmic or secreted lipoprotein OS=Methylomicrobium album BG8 OX=686340 GN=Metal_3655 PE=4 SV=1 | #N/A | #N/A | #N/A | 1 | 1 | 1.91 | #N/A | #N/A | #N/A |
| H8GJI7 | Chaperone SurA OS=Methylomicrobium album BG8 OX=686340 GN=surA PE=3 SV=1 | #N/A | #N/A | #N/A | 1 | 1 | 2.24 | 1 | 1 | 2.61 |
| H8GMA4 | CBS domain-containing protein OS=Methylomicrobium album BG8 OX=686340 GN=Metal_2947 PE=3 SV=1 | #N/A | #N/A | #N/A | #N/A | #N/A | #N/A | 3 | 1 | 7.4 |
| H8GIK8 | Chemotaxis signal transduction protein OS=Methylomicrobium album BG8 OX=686340 GN=Metal_1229 PE=4 SV=1 | #N/A | #N/A | #N/A | #N/A | #N/A | #N/A | 2 | 2 | 5.02 |
| H8GFU5 | Isoleucine--tRNA ligase OS=Methylomicrobium album BG8 OX=686340 GN=ileS PE=3 SV=1 | #N/A | #N/A | #N/A | #N/A | #N/A | #N/A | 2 | 1 | 4.16 |
| H8GNN0 | Type VI secretion protein, VC_A0107 family OS=Methylomicrobium album BG8 OX=686340 GN=Metal_3108 PE=4 SV=1 | #N/A | #N/A | #N/A | #N/A | #N/A | #N/A | 2 | 1 | 4.07 |
| H8GHV3 | Amidophosphoribosyltransferase OS=Methylomicrobium album BG8 OX=686340 GN=purF PE=3 SV=1 | #N/A | #N/A | #N/A | #N/A | #N/A | #N/A | 2 | 1 | 4.34 |
| H8GHC3 | Putative ATPase OS=Methylomicrobium album BG8 OX=686340 GN=Metal_1096 PE=4 SV=1 | #N/A | #N/A | #N/A | #N/A | #N/A | #N/A | 2 | 2 | 4.07 |
| H8GM13 | Putative transcriptional regulator OS=Methylomicrobium album BG8 OX=686340 GN=Metal_0352 PE=4 SV=1 | #N/A | #N/A | #N/A | #N/A | #N/A | #N/A | 2 | 1 | 4.66 |
| H8GJR6 | Uncharacterized protein OS=Methylomicrobium album BG8 OX=686340 GN=Metal_3966 PE=4 SV=1 | #N/A | #N/A | #N/A | #N/A | #N/A | #N/A | 1 | 1 | 2.57 |
| H8GQZ7 | PAS domain S-box OS=Methylomicrobium album BG8 OX=686340 GN=Metal_0827 PE=4 SV=1 | #N/A | #N/A | #N/A | #N/A | #N/A | #N/A | 1 | 1 | 2 |
| H8GGN3 | DNA repair protein radc OS=Methylomicrobium album BG8 OX=686340 GN=Metal_1010 PE=3 SV=1 | #N/A | #N/A | #N/A | #N/A | #N/A | #N/A | 1 | 1 | 3.11 |
| H8GKR5 | NADH:ubiquinone oxidoreductase, NADH-binding (51 kD) subunit OS=Methylomicrobium album BG8 OX=686340 GN=Metal_1452 PE=3 SV=1 | #N/A | #N/A | #N/A | #N/A | #N/A | #N/A | 1 | 1 | 2.49 |
| H8GG05 | RNA polymerase-associated protein RapA OS=Methylomicrobium album BG8 OX=686340 GN=rapA PE=3 SV=1 | #N/A | #N/A | #N/A | #N/A | #N/A | #N/A | 1 | 1 | 1.93 |
| H8GQ29 | 50S ribosomal protein L5 OS=Methylomicrobium album BG8 OX=686340 GN=rplE PE=3 SV=1 | #N/A | #N/A | #N/A | #N/A | #N/A | #N/A | 1 | 1 | 2.09 |
| H8GPI5 | Outer membrane protein assembly factor BamD OS=Methylomicrobium album BG8 OX=686340 GN=bamD PE=3 SV=1 | #N/A | #N/A | #N/A | #N/A | #N/A | #N/A | 1 | 1 | 2.3 |
| H8GKF5 | Anti-sigma factor antagonist OS=Methylomicrobium album BG8 OX=686340 GN=Metal_2748 PE=3 SV=1 | #N/A | #N/A | #N/A | #N/A | #N/A | #N/A | 1 | 1 | 1.9 |
| H8GJN4 | Putative transcriptional regulator with HTH domain OS=Methylomicrobium album BG8 OX=686340 GN=Metal_3927 PE=4 SV=1 | #N/A | #N/A | #N/A | #N/A | #N/A | #N/A | 1 | 1 | 2.05 |
| H8GI45 | Methanol oxidation system protein MoxJ OS=Methylomicrobium album BG8 OX=686340 GN=Metal_2470 PE=4 SV=1 | #N/A | #N/A | #N/A | #N/A | #N/A | #N/A | 1 | 1 | 3.95 |
| H8GLY3 | Putative PLP-dependent enzyme possibly involved in cell wall biogenesis OS=Methylomicrobium album BG8 OX=686340 GN=Metal_0320 PE=3 SV=1 | #N/A | #N/A | #N/A | #N/A | #N/A | #N/A | 1 | 1 | 2.81 |
| H8GG42 | Putative metal-binding integral membrane protein OS=Methylomicrobium album BG8 OX=686340 GN=Metal_2220 PE=4 SV=1 | #N/A | #N/A | #N/A | #N/A | #N/A | #N/A | 1 | 1 | 2.63 |
| H8GIL5 | Fibronectin type III domain-containing protein OS=Methylomicrobium album BG8 OX=686340 GN=Metal_1236 PE=4 SV=1 | #N/A | #N/A | #N/A | #N/A | #N/A | #N/A | 1 | 1 | 2.11 |
| H8GKU9 | RNA pyrophosphohydrolase OS=Methylomicrobium album BG8 OX=686340 GN=rppH PE=3 SV=1 | #N/A | #N/A | #N/A | #N/A | #N/A | #N/A | 1 | 1 | 2.3 |
| H8GL47 | Response regulator containing a CheY-like receiver domain and an HTH DNA-binding domain OS=Methylomicrobium album BG8 OX=686340 GN=Metal_2839 PE=4 SV=1 | #N/A | #N/A | #N/A | #N/A | #N/A | #N/A | 1 | 1 | 2.3 |
| H8GMC6 | RNA polymerase sigma factor RpoD OS=Methylomicrobium album BG8 OX=686340 GN=rpoD PE=3 SV=1 | #N/A | #N/A | #N/A | #N/A | #N/A | #N/A | 1 | 1 | 2.29 |
| H8GKK0 | Acyl-[acyl-carrier-protein]--UDP-N-acetylglucosamine O-acyltransferase OS=Methylomicrobium album BG8 OX=686340 GN=lpxA PE=3 SV=1 | #N/A | #N/A | #N/A | #N/A | #N/A | #N/A | 1 | 1 | 2.12 |
| H8GJB7 | Lipoprotein releasing system, transmembrane protein, LolC/E family OS=Methylomicrobium album BG8 OX=686340 GN=Metal_1309 PE=3 SV=1 | #N/A | #N/A | #N/A | #N/A | #N/A | #N/A | 1 | 1 | 2.45 |
| H8GKJ9 | Ribonuclease HII OS=Methylomicrobium album BG8 OX=686340 GN=rnhB PE=3 SV=1 | #N/A | #N/A | #N/A | #N/A | #N/A | #N/A | 1 | 1 | 3.27 |
| H8GKB0 | ATPase component of various ABC-type transport systems with duplicated ATPase domain OS=Methylomicrobium album BG8 OX=686340 GN=Metal_2695 PE=4 SV=1 | #N/A | #N/A | #N/A | #N/A | #N/A | #N/A | 1 | 1 | 1.91 |
| H8GKC8 | Cation/multidrug efflux pump OS=Methylomicrobium album BG8 OX=686340 GN=Metal_2717 PE=4 SV=1 | #N/A | #N/A | #N/A | #N/A | #N/A | #N/A | 1 | 1 | 2.43 |
| H8GIR3 | Ubiquinone biosynthesis accessory factor UbiJ OS=Methylomicrobium album BG8 OX=686340 GN=ubiJ PE=3 SV=1 | #N/A | #N/A | #N/A | #N/A | #N/A | #N/A | 1 | 1 | 1.9 |
| H8GH32 | ABC-type antimicrobial peptide transport system, permease component OS=Methylomicrobium album BG8 OX=686340 GN=Metal_3660 PE=3 SV=1 | #N/A | #N/A | #N/A | #N/A | #N/A | #N/A | 1 | 1 | 2.3 |
| H8GNX4 | Urease subunit gamma OS=Methylomicrobium album BG8 OX=686340 GN=ureA PE=3 SV=1 | #N/A | #N/A | #N/A | #N/A | #N/A | #N/A | 1 | 1 | 1.9 |
| H8GJT2 | Phosphate-selective porin OS=Methylomicrobium album BG8 OX=686340 GN=Metal_3985 PE=4 SV=1 | #N/A | #N/A | #N/A | #N/A | #N/A | #N/A | 1 | 1 | 2.98 |
| H8GN71 | DNA gyrase subunit A OS=Methylomicrobium album BG8 OX=686340 GN=gyrA PE=3 SV=1 | #N/A | #N/A | #N/A | #N/A | #N/A | #N/A | 1 | 1 | 2.37 |
| H8GLD5 | DNA topoisomerase 1 OS=Methylomicrobium album BG8 OX=686340 GN=topA PE=3 SV=1 | #N/A | #N/A | #N/A | #N/A | #N/A | #N/A | 1 | 1 | 2.12 |
| H8GMH2 | GDT1 family protein OS=Methylomicrobium album BG8 OX=686340 GN=Metal_3016 PE=3 SV=1 | #N/A | #N/A | #N/A | #N/A | #N/A | #N/A | 1 | 1 | 2.34 |
| H8GQP1 | Lon protease OS=Methylomicrobium album BG8 OX=686340 GN=lon PE=2 SV=1 | #N/A | #N/A | #N/A | #N/A | #N/A | #N/A | 1 | 1 | 2.6 |
| H8GH42 | 2-polyprenyl-6-methoxyphenol 4-hydroxylase OS=Methylomicrobium album BG8 OX=686340 GN=Metal_3670 PE=3 SV=1 | #N/A | #N/A | #N/A | #N/A | #N/A | #N/A | 1 | 1 | 2.08 |
| H8GGL0 | Uncharacterized protein OS=Methylomicrobium album BG8 OX=686340 GN=Metal_0987 PE=3 SV=1 | #N/A | #N/A | #N/A | #N/A | #N/A | #N/A | 1 | 1 | 2.17 |
| H8GJK4 | Diguanylate cyclase (GGDEF) domain-containing protein OS=Methylomicrobium album BG8 OX=686340 GN=Metal_2654 PE=4 SV=1 | #N/A | #N/A | #N/A | #N/A | #N/A | #N/A | 1 | 1 | 2.3 |
| H8GPL4 | DNA/RNA helicase, superfamily II OS=Methylomicrobium album BG8 OX=686340 GN=Metal_0631 PE=4 SV=1 | #N/A | #N/A | #N/A | #N/A | #N/A | #N/A | 1 | 1 | 1.99 |
| H8GPZ4 | Serine--tRNA ligase OS=Methylomicrobium album BG8 OX=686340 GN=serS PE=3 SV=1 | #N/A | #N/A | #N/A | #N/A | #N/A | #N/A | 1 | 1 | 2.42 |
| H8GNY7 | Transcriptional regulator OS=Methylomicrobium album BG8 OX=686340 GN=Metal_0559 PE=3 SV=1 | #N/A | #N/A | #N/A | #N/A | #N/A | #N/A | 1 | 1 | 2 |
| H8GJ71 | Mutator family transposase OS=Methylomicrobium album BG8 OX=686340 GN=Metal_0017 PE=3 SV=1 | #N/A | #N/A | #N/A | #N/A | #N/A | #N/A | 1 | 1 | 2.61 |
| H8GJA1 | Putative addiction module killer protein OS=Methylomicrobium album BG8 OX=686340 GN=Metal_1291 PE=4 SV=1 | #N/A | #N/A | #N/A | #N/A | #N/A | #N/A | 1 | 1 | 2.06 |
| H8GMA8 | Magnesium transport protein CorA OS=Methylomicrobium album BG8 OX=686340 GN=corA PE=3 SV=1 | #N/A | #N/A | #N/A | #N/A | #N/A | #N/A | 1 | 1 | 2.54 |
| H8GP30 | Putative helicase OS=Methylomicrobium album BG8 OX=686340 GN=Metal_0607 PE=4 SV=1 | #N/A | #N/A | #N/A | #N/A | #N/A | #N/A | 1 | 1 | 1.95 |
| H8GPF9 | Adenine-specific DNA methylase containing a Zn-ribbon OS=Methylomicrobium album BG8 OX=686340 GN=Metal_3235 PE=4 SV=1 | #N/A | #N/A | #N/A | #N/A | #N/A | #N/A | 1 | 1 | 2.3 |
| H8GQH0 | 3-hydroxyacyl-CoA dehydrogenase OS=Methylomicrobium album BG8 OX=686340 GN=Metal_2038 PE=4 SV=1 | #N/A | #N/A | #N/A | #N/A | #N/A | #N/A | 1 | 1 | 2.66 |
| H8GIC1 | Helix-turn-helix protein OS=Methylomicrobium album BG8 OX=686340 GN=Metal_3789 PE=4 SV=1 | #N/A | #N/A | #N/A | #N/A | #N/A | #N/A | 1 | 1 | 2.5 |
| H8GQC6 | DNA translocase FtsK OS=Methylomicrobium album BG8 OX=686340 GN=Metal_0751 PE=3 SV=1 | #N/A | #N/A | #N/A | #N/A | #N/A | #N/A | 1 | 1 | 2.42 |
| H8GG08 | Uncharacterized protein OS=Methylomicrobium album BG8 OX=686340 GN=Metal_0939 PE=4 SV=1 | #N/A | #N/A | #N/A | #N/A | #N/A | #N/A | 1 | 1 | 1.99 |
| H8GR39 | Uncharacterized protein OS=Methylomicrobium album BG8 OX=686340 GN=Metal_2111 PE=4 SV=1 | #N/A | #N/A | #N/A | #N/A | #N/A | #N/A | 1 | 1 | 1.93 |
| H8GLS3 | Protein HflC OS=Methylomicrobium album BG8 OX=686340 GN=Metal_2919 PE=3 SV=1 | #N/A | #N/A | #N/A | #N/A | #N/A | #N/A | 1 | 1 | 2.04 |
| H8GGF8 | Methane monooxygenase/ammonia monooxygenase, subunit B OS=Methylomicrobium album BG8 OX=686340 GN=Metal_3591 PE=4 SV=1 | #N/A | #N/A | #N/A | #N/A | #N/A | #N/A | 1 | 1 | 2.36 |
| H8GPD5 | 6-phosphogluconate dehydrogenase, decarboxylating OS=Methylomicrobium album BG8 OX=686340 GN=Metal_3210 PE=3 SV=1 | #N/A | #N/A | #N/A | #N/A | #N/A | #N/A | 1 | 1 | 1.94 |
| H8GN04 | Diguanylate cyclase (GGDEF) domain-containing protein OS=Methylomicrobium album BG8 OX=686340 GN=Metal_3038 PE=4 SV=1 | #N/A | #N/A | #N/A | #N/A | #N/A | #N/A | 1 | 1 | 2.04 |
| H8GI72 | Putative tRNA(5-methylaminomethyl-2-thiouridylate) methyltransferase with PP-loop ATPase domain OS=Methylomicrobium album BG8 OX=686340 GN=Metal_2497 PE=4 SV=1 | #N/A | #N/A | #N/A | #N/A | #N/A | #N/A | 1 | 1 | 2.1 |
| H8GR81 | Aspartokinase OS=Methylomicrobium album BG8 OX=686340 GN=Metal_2155 PE=3 SV=1 | #N/A | #N/A | #N/A | #N/A | #N/A | #N/A | 1 | 1 | 2.39 |
| H8GR90 | Lipid-A-disaccharide synthase OS=Methylomicrobium album BG8 OX=686340 GN=lpxB PE=3 SV=1 | #N/A | #N/A | #N/A | #N/A | #N/A | #N/A | 1 | 1 | 2.45 |
| H8GNN2 | Outer membrane autotransporter barrel domain-containing protein OS=Methylomicrobium album BG8 OX=686340 GN=Metal_3112 PE=4 SV=1 | #N/A | #N/A | #N/A | #N/A | #N/A | #N/A | 1 | 1 | 2.87 |
| H8GHD8 | Putative sugar kinase OS=Methylomicrobium album BG8 OX=686340 GN=Metal_2363 PE=4 SV=1 | #N/A | #N/A | #N/A | #N/A | #N/A | #N/A | 1 | 1 | 1.98 |
| H8GIT3 | ATP-dependent helicase HrpA OS=Methylomicrobium album BG8 OX=686340 GN=Metal_2557 PE=4 SV=1 | #N/A | #N/A | #N/A | #N/A | #N/A | #N/A | 1 | 1 | 1.98 |
| H8GNT1 | Non-specific serine/threonine protein kinase OS=Methylomicrobium album BG8 OX=686340 GN=Metal_3163 PE=4 SV=1 | #N/A | #N/A | #N/A | #N/A | #N/A | #N/A | 1 | 1 | 2.51 |
| H8GIP6 | EH_Signature domain-containing protein OS=Methylomicrobium album BG8 OX=686340 GN=Metal_1273 PE=4 SV=1 | #N/A | #N/A | #N/A | #N/A | #N/A | #N/A | 1 | 1 | 2.59 |
| Bolded proteins are referenced in Table 2  Italicized proteins are referenced in Table 3 | | | | | | | | | | |

**Supplemental Table S3.** Abundant proteome hits (using BLASTp) from purified *M. album* BG8 S-layer units identified in other members of the Methylococcaceae family (taxid:135618)

| **Metal_1783** | **EIC29550.1** | **H8GNF7** | **Flagellar hook protein FlgE** | | | | | |
| --- | --- | --- | --- | --- | --- | --- | --- | --- |
| *Description* | *Scientific Name* | *Max Score* | *Total Score* | *Query Cover* | *E value* | *Per. ident* | *Acc. Len* | *Accession* |
| flagellar hook protein FlgE [Methylomicrobium agile] | Methylomicrobium agile | 797 | 797 | 100% | 0 | 99.07 | 431 | [WP_031431566.1](https://www.ncbi.nlm.nih.gov/protein/WP_031431566.1?report=genbank&log$=prottop&blast_rank=1&RID=864PNV0C013) |
| flagellar hook protein FlgE [Methylomicrobium sp. RS1] | Methylomicrobium sp. RS1 | 747 | 747 | 100% | 0 | 91.88 | 431 | [WP_202053483.1](https://www.ncbi.nlm.nih.gov/protein/WP_202053483.1?report=genbank&log$=prottop&blast_rank=2&RID=864PNV0C013) |
| flagellar hook protein FlgE [Methylobacter] | Methylobacter | 622 | 622 | 100% | 0 | 73.27 | 432 | [WP_020157946.1](https://www.ncbi.nlm.nih.gov/protein/WP_020157946.1?report=genbank&log$=prottop&blast_rank=3&RID=864PNV0C013) |
| flagellar hook protein FlgE [Methylosarcina fibrata] | Methylosarcina fibrata | 592 | 592 | 100% | 0 | 69.21 | 432 | [WP_020565742.1](https://www.ncbi.nlm.nih.gov/protein/WP_020565742.1?report=genbank&log$=prottop&blast_rank=4&RID=864PNV0C013) |
| flagellar hook-basal body protein [Methylobacter sp.] | Methylobacter sp. | 510 | 510 | 99% | 2.00E-179 | 65.2 | 428 | [PPD05312.1](https://www.ncbi.nlm.nih.gov/protein/PPD05312.1?report=genbank&log$=prottop&blast_rank=5&RID=864PNV0C013) |
| **Metal_1728** | **EIC29497.1** | **H8GMU5** | **Flagellin** | | | | | |
| *Description* | *Scientific Name* | *Max Score* | *Total Score* | *Query Cover* | *E value* | *Per. ident* | *Acc. Len* | *Accession* |
| flagellin/flagellar hook associated protein [Methylomicrobium] | Methylomicrobium | 1009 | 1009 | 100% | 0 | 100 | 761 | [WP_005371394.1](https://www.ncbi.nlm.nih.gov/protein/WP_005371394.1?report=genbank&log$=prottop&blast_rank=1&RID=864ZV0DX013) |
| hypothetical protein [Methylomicrobium sp. RS1] | Methylomicrobium sp. RS1 | 402 | 626 | 65% | 2.00E-129 | 88.03 | 632 | [WP_236994296.1](https://www.ncbi.nlm.nih.gov/protein/WP_236994296.1?report=genbank&log$=prottop&blast_rank=2&RID=864ZV0DX013) |
| flagellin [Methylococcales bacterium] | Methylococcales bacterium | 224 | 355 | 56% | 7.00E-63 | 58.1 | 581 | [RIZ72076.1](https://www.ncbi.nlm.nih.gov/protein/RIZ72076.1?report=genbank&log$=prottop&blast_rank=3&RID=864ZV0DX013) |
| flagellin [Methylobacter sp.] | Methylobacter sp. | 226 | 343 | 56% | 1.00E-62 | 48.99 | 737 | [TAN70616.1](https://www.ncbi.nlm.nih.gov/protein/TAN70616.1?report=genbank&log$=prottop&blast_rank=4&RID=864ZV0DX013) |
| flagellin [Methylovulum sp.] | Methylovulum sp. | 217 | 341 | 55% | 2.00E-59 | 50.45 | 709 | [TAL46828.1](https://www.ncbi.nlm.nih.gov/protein/TAL46828.1?report=genbank&log$=prottop&blast_rank=5&RID=864ZV0DX013) |
| flagellin [Methylococcaceae bacterium] | Methylococcaceae bacterium | 262 | 410 | 55% | 1.00E-76 | 64.22 | 609 | [NOT11437.1](https://www.ncbi.nlm.nih.gov/protein/NOT11437.1?report=genbank&log$=prottop&blast_rank=6&RID=864ZV0DX013) |
| flagellin [Methylomarinum vadi] | Methylomarinum vadi | 185 | 291 | 47% | 1.00E-49 | 49.25 | 487 | [WP_031435694.1](https://www.ncbi.nlm.nih.gov/protein/WP_031435694.1?report=genbank&log$=prottop&blast_rank=7&RID=864ZV0DX013) |
| B-type flagellin [Candidatus Methylobacter favarea] | Candidatus Methylobacter favarea | 193 | 326 | 46% | 1.00E-52 | 55.38 | 504 | [CAA9892324.1](https://www.ncbi.nlm.nih.gov/protein/CAA9892324.1?report=genbank&log$=prottop&blast_rank=8&RID=864ZV0DX013) |
| flagellin [Methylothermaceae bacteria B42] | Methylothermaceae bacteria B42 | 187 | 286 | 46% | 3.00E-50 | 49.62 | 495 | [KXJ40563.1](https://www.ncbi.nlm.nih.gov/protein/KXJ40563.1?report=genbank&log$=prottop&blast_rank=9&RID=864ZV0DX013) |
| hypothetical protein VZ94_14540 [Methylocucumis oryzae] | Methylocucumis oryzae | 216 | 340 | 46% | 3.00E-59 | 53.96 | 709 | [KJV05952.1](https://www.ncbi.nlm.nih.gov/protein/KJV05952.1?report=genbank&log$=prottop&blast_rank=10&RID=864ZV0DX013) |
| **Metal_1116** | **EIC28933.1** | **H8GHU9** | **Prepilin-type N-terminal cleavage/methylation domain-containing protein** | | | | | |
| *Description* | *Scientific Name* | *Max Score* | *Total Score* | *Query Cover* | *E value* | *Per. ident* | *Acc. Len* | *Accession* |
| TPA: prepilin-type N-terminal cleavage/methylation domain-containing protein [Methyloprofundus sp.] | Methyloprofundus sp. | 129 | 129 | 90% | 4.00E-38 | 63.19 | 141 | [HIG65939.1](https://www.ncbi.nlm.nih.gov/protein/HIG65939.1?report=genbank&log$=prottop&blast_rank=1&RID=86587YNK016) |
| prepilin-type N-terminal cleavage/methylation domain-containing protein [Methylococcaceae bacterium FWC3] | Methylococcaceae bacterium FWC3 | 70.5 | 70.5 | 90% | 7.00E-15 | 46.58 | 152 | [RYU58359.1](https://www.ncbi.nlm.nih.gov/protein/RYU58359.1?report=genbank&log$=prottop&blast_rank=2&RID=86587YNK016) |
| pilin [Methylovulum sp.] | Methylovulum sp. | 58.5 | 58.5 | 90% | 3.00E-10 | 38.89 | 177 | [MCF7986727.1](https://www.ncbi.nlm.nih.gov/protein/MCF7986727.1?report=genbank&log$=prottop&blast_rank=3&RID=86587YNK016) |
| pilin [Methylomicrobium sp.] | Methylomicrobium sp. | 56.6 | 56.6 | 90% | 2.00E-09 | 40.54 | 180 | [MBS3953249.1](https://www.ncbi.nlm.nih.gov/protein/MBS3953249.1?report=genbank&log$=prottop&blast_rank=4&RID=86587YNK016) |
| pilin [Methyloglobulus sp.] | Methyloglobulus sp. | 46.2 | 46.2 | 90% | 1.00E-05 | 35.14 | 182 | [NOU43879.1](https://www.ncbi.nlm.nih.gov/protein/NOU43879.1?report=genbank&log$=prottop&blast_rank=5&RID=86587YNK016) |
| prepilin-type N-terminal cleavage/methylation domain-containing protein [Methylomagnum ishizawai] | Methylomagnum ishizawai | 63.2 | 63.2 | 89% | 4.00E-12 | 46.15 | 150 | [BBL73386.1](https://www.ncbi.nlm.nih.gov/protein/BBL73386.1?report=genbank&log$=prottop&blast_rank=6&RID=86587YNK016) |
| prepilin-type cleavage/methylation domain-containing protein [Methylomonas sp.] | Methylomonas sp. | 47.8 | 47.8 | 89% | 4.00E-06 | 38.24 | 169 | [PPD29451.1](https://www.ncbi.nlm.nih.gov/protein/PPD29451.1?report=genbank&log$=prottop&blast_rank=7&RID=86587YNK016) |
| **Metal_0147** | **EIC28015.1** | **H8GKK7** | **Ca^2+^-binding protein, RTX toxin** | | | | | |
| *Description* | *Scientific Name* | *Max Score* | *Total Score* | *Query Cover* | *E value* | *Per. ident* | *Acc. Len* | *Accession* |
| calcium-binding protein [Methylomicrobium agile] | Methylomicrobium agile | 1868 | 1868 | 100% | 0 | 98.48 | 1056 | [WP_031430404.1](https://www.ncbi.nlm.nih.gov/protein/WP_031430404.1?report=genbank&log$=prottop&blast_rank=1&RID=865F6HVT013) |
| calcium-binding protein [Methylomicrobium sp. RS1] | Methylomicrobium sp. RS1 | 1697 | 1697 | 100% | 0 | 88.43 | 1037 | [WP_202050871.1](https://www.ncbi.nlm.nih.gov/protein/WP_202050871.1?report=genbank&log$=prottop&blast_rank=2&RID=865F6HVT013) |
| calcium-binding protein [Methylomicrobium lacus] | Methylomicrobium lacus | 1535 | 1535 | 100% | 0 | 80.99 | 1036 | [WP_152539446.1](https://www.ncbi.nlm.nih.gov/protein/WP_152539446.1?report=genbank&log$=prottop&blast_rank=3&RID=865F6HVT013) |
| calcium-binding protein [Methylomonas rhizoryzae] | Methylomonas rhizoryzae | 662 | 714 | 92% | 0 | 47.84 | 910 | [WP_150050555.1](https://www.ncbi.nlm.nih.gov/protein/WP_150050555.1?report=genbank&log$=prottop&blast_rank=4&RID=865F6HVT013) |
| calcium-binding protein [Methylotuvimicrobium buryatense] | Methylotuvimicrobium buryatense | 620 | 620 | 89% | 0 | 47.22 | 893 | [WP_017842713.1](https://www.ncbi.nlm.nih.gov/protein/WP_017842713.1?report=genbank&log$=prottop&blast_rank=5&RID=865F6HVT013) |
| **Metal_3821** | **EIC31463.1** | **H8GIF1** | **Type 1 secretion C-terminal target domain protein** | | | | | |
| *Description* | *Scientific Name* | *Max Score* | *Total Score* | *Query Cover* | *E value* | *Per. ident* | *Acc. Len* | *Accession* |
| retention module-containing protein [Methylomicrobium agile] | Methylomicrobium agile | 103 | 550 | 47% | 1.00E-20 | 52.05 | 3280 | [WP_084675305.1](https://www.ncbi.nlm.nih.gov/protein/WP_084675305.1?report=genbank&log$=prottop&blast_rank=1&RID=865MGRY1016) |
| type I secretion C-terminal target domain-containing protein [Methylococcaceae bacterium] | Methylococcaceae bacterium | 85.5 | 163 | 41% | 2.00E-15 | 28.11 | 939 | [NJD05510.1](https://www.ncbi.nlm.nih.gov/protein/NJD05510.1?report=genbank&log$=prottop&blast_rank=2&RID=865MGRY1016) |
| type I secretion C-terminal target domain-containing protein [Methylovulum sp.] | Methylovulum sp. | 88.2 | 146 | 19% | 2.00E-16 | 40.84 | 691 | [TAL50852.1](https://www.ncbi.nlm.nih.gov/protein/TAL50852.1?report=genbank&log$=prottop&blast_rank=3&RID=865MGRY1016) |
| putative Ig domain-containing protein [Methylicorpusculum oleiharenae] | Methylicorpusculum oleiharenae | 78.2 | 669 | 18% | 4.00E-13 | 54.76 | 2587 | [WP_159658860.1](https://www.ncbi.nlm.nih.gov/protein/WP_159658860.1?report=genbank&log$=prottop&blast_rank=4&RID=865MGRY1016) |
| hypothetical protein EPN17_15535 [Methylobacter sp.] | Methylobacter sp. | 82.4 | 420 | 16% | 2.00E-14 | 35.47 | 1009 | [TAN65811.1](https://www.ncbi.nlm.nih.gov/protein/TAN65811.1?report=genbank&log$=prottop&blast_rank=5&RID=865MGRY1016) |
| hypothetical protein CTY16_02775 [Methylobacter sp.] | Methylobacter sp. | 80.1 | 504 | 16% | 9.00E-14 | 45.45 | 1035 | [PPD49895.1](https://www.ncbi.nlm.nih.gov/protein/PPD49895.1?report=genbank&log$=prottop&blast_rank=6&RID=865MGRY1016) |
| calcium-binding protein [Methylococcaceae bacterium] | Methylococcaceae bacterium | 76.3 | 650 | 15% | 2.00E-12 | 50.49 | 1923 | [MBK8816867.1](https://www.ncbi.nlm.nih.gov/protein/MBK8816867.1?report=genbank&log$=prottop&blast_rank=7&RID=865MGRY1016) |
| hypothetical protein CTY16_05180 [Methylobacter sp.] | Methylobacter sp. | 75.9 | 352 | 15% | 2.00E-12 | 35.94 | 1556 | [PPD48861.1](https://www.ncbi.nlm.nih.gov/protein/PPD48861.1?report=genbank&log$=prottop&blast_rank=8&RID=865MGRY1016) |
| hypothetical protein EPN89_17425 [Methylovulum sp.] | Methylovulum sp. | 84.7 | 512 | 14% | 4.00E-15 | 40.5 | 1167 | [TAL42544.1](https://www.ncbi.nlm.nih.gov/protein/TAL42544.1?report=genbank&log$=prottop&blast_rank=9&RID=865MGRY1016) |
| calcium-binding protein [Methylovulum psychrotolerans] | Methylovulum psychrotolerans | 87.8 | 661 | 14% | 5.00E-16 | 45.33 | 1341 | [WP_215477966.1](https://www.ncbi.nlm.nih.gov/protein/WP_215477966.1?report=genbank&log$=prottop&blast_rank=10&RID=865MGRY1016) |
| Ig-like domain-containing protein [Methylomagnum ishizawai] | Methylomagnum ishizawai | 82.4 | 82.4 | 14% | 3.00E-14 | 38.58 | 2799 | [WP_125468845.1](https://www.ncbi.nlm.nih.gov/protein/WP_125468845.1?report=genbank&log$=prottop&blast_rank=11&RID=865MGRY1016) |
| **Metal_1447** | **EIC29232.1** | **H8GKR0** | **Uncharacterized protein** | | | | | |
| *Description* | *Scientific Name* | *Max Score* | *Total Score* | *Query Cover* | *E value* | *Per. ident* | *Acc. Len* | *Accession* |
| hypothetical protein [Methylomicrobium agile] | Methylomicrobium agile | 576 | 576 | 100% | 0 | 99.67 | 299 | [WP_031431375.1](https://www.ncbi.nlm.nih.gov/protein/WP_031431375.1?report=genbank&log$=prottop&blast_rank=1&RID=865UVCK6016) |
| hypothetical protein [Methylomicrobium lacus] | Methylomicrobium lacus | 429 | 429 | 100% | 9.00E-152 | 78.62 | 304 | [WP_024297952.1](https://www.ncbi.nlm.nih.gov/protein/WP_024297952.1?report=genbank&log$=prottop&blast_rank=2&RID=865UVCK6016) |
| porin family protein [Methylomicrobium sp. RS1] | Methylomicrobium sp. RS1 | 386 | 386 | 100% | 9.00E-135 | 70 | 295 | [WP_202051219.1](https://www.ncbi.nlm.nih.gov/protein/WP_202051219.1?report=genbank&log$=prottop&blast_rank=3&RID=865UVCK6016) |
| hypothetical protein [Methylosarcina fibrata] | Methylosarcina fibrata | 319 | 319 | 99% | 2.00E-108 | 58.19 | 292 | [WP_026223532.1](https://www.ncbi.nlm.nih.gov/protein/WP_026223532.1?report=genbank&log$=prottop&blast_rank=4&RID=865UVCK6016) |
| porin family protein [Methylobacter sp.] | Methylobacter sp. | 366 | 366 | 91% | 5.00E-127 | 65.16 | 289 | [TAN65355.1](https://www.ncbi.nlm.nih.gov/protein/TAN65355.1?report=genbank&log$=prottop&blast_rank=5&RID=865UVCK6016) |
| porin family protein [Methylococcales bacterium] | Methylococcales bacterium | 361 | 361 | 91% | 1.00E-124 | 60.88 | 317 | [MSP28215.1](https://www.ncbi.nlm.nih.gov/protein/MSP28215.1?report=genbank&log$=prottop&blast_rank=6&RID=865UVCK6016) |
| **Metal_0659** | **EIC28502.1** | **H8GPP0** | **Uncharacterized protein** | | | | | |
| *Description* | *Scientific Name* | *Max Score* | *Total Score* | *Query Cover* | *E value* | *Per. ident* | *Acc. Len* | *Accession* |
| hypothetical protein [Methylomicrobium] | Methylomicrobium | 228 | 228 | 92% | 7.00E-78 | 100 | 118 | [WP_005369594.1](https://www.ncbi.nlm.nih.gov/protein/WP_005369594.1?report=genbank&log$=prottop&blast_rank=1&RID=8660WNVN013) |
| hypothetical protein [Methylomicrobium sp. RS1] | Methylomicrobium sp. RS1 | 205 | 205 | 92% | 6.00E-69 | 89.91 | 118 | [WP_202051802.1](https://www.ncbi.nlm.nih.gov/protein/WP_202051802.1?report=genbank&log$=prottop&blast_rank=2&RID=8660WNVN013) |
| hypothetical protein [Methylomonas sp. EFPC1] | Methylomonas sp. EFPC1 | 177 | 177 | 92% | 4.00E-58 | 75.23 | 118 | [WP_205450661.1](https://www.ncbi.nlm.nih.gov/protein/WP_205450661.1?report=genbank&log$=prottop&blast_rank=3&RID=8660WNVN013) |
| hypothetical protein [Methylomonas sp. ZR1] | Methylomonas sp. ZR1 | 177 | 177 | 92% | 7.00E-58 | 75.23 | 118 | [WP_171695834.1](https://www.ncbi.nlm.nih.gov/protein/WP_171695834.1?report=genbank&log$=prottop&blast_rank=4&RID=8660WNVN013) |
| hypothetical protein [Methylomonas sp. 11b] | Methylomonas sp. 11b | 177 | 177 | 92% | 9.00E-58 | 75.23 | 118 | [WP_026604202.1](https://www.ncbi.nlm.nih.gov/protein/WP_026604202.1?report=genbank&log$=prottop&blast_rank=5&RID=8660WNVN013) |
| hypothetical protein [Methylomonas sp. MK1] | Methylomonas sp. MK1 | 175 | 175 | 92% | 3.00E-57 | 74.31 | 118 | [WP_020485651.1](https://www.ncbi.nlm.nih.gov/protein/WP_020485651.1?report=genbank&log$=prottop&blast_rank=6&RID=8660WNVN013) |
| hypothetical protein [Methylomonas sp. LWB] | Methylomonas sp. LWB | 174 | 174 | 92% | 1.00E-56 | 73.39 | 120 | [WP_071160780.1](https://www.ncbi.nlm.nih.gov/protein/WP_071160780.1?report=genbank&log$=prottop&blast_rank=7&RID=8660WNVN013) |
| hypothetical protein [Methylomonas koyamae] | Methylomonas koyamae | 174 | 174 | 92% | 2.00E-56 | 71.56 | 118 | [WP_064028347.1](https://www.ncbi.nlm.nih.gov/protein/WP_064028347.1?report=genbank&log$=prottop&blast_rank=8&RID=8660WNVN013) |
| **Metal_3919** | **EIC31556.1** | **H8GJ49** | **Outer membrane cobalamin receptor protein** | | | | | |
| *Description* | *Scientific Name* | *Max Score* | *Total Score* | *Query Cover* | *E value* | *Per. ident* | *Acc. Len* | *Accession* |
| TonB-dependent receptor [Methylomicrobium agile] | Methylomicrobium agile | 1412 | 1412 | 100% | 0 | 99.71 | 691 | [WP_031430284.1](https://www.ncbi.nlm.nih.gov/protein/WP_031430284.1?report=genbank&log$=prottop&blast_rank=1&RID=8665CEF5016) |
| TonB-dependent receptor [Methylomicrobium sp. RS1] | Methylomicrobium sp. RS1 | 1380 | 1380 | 100% | 0 | 97.25 | 691 | [WP_202053908.1](https://www.ncbi.nlm.nih.gov/protein/WP_202053908.1?report=genbank&log$=prottop&blast_rank=2&RID=8665CEF5016) |
| TonB-dependent receptor [Methylococcales bacterium] | Methylococcales bacterium | 749 | 749 | 98% | 0 | 54.1 | 685 | [MBT7575564.1](https://www.ncbi.nlm.nih.gov/protein/MBT7575564.1?report=genbank&log$=prottop&blast_rank=3&RID=8665CEF5016) |
| TonB-dependent receptor [Methylococcales bacterium] | Methylococcales bacterium | 746 | 746 | 98% | 0 | 53.96 | 685 | [MBT3698098.1](https://www.ncbi.nlm.nih.gov/protein/MBT3698098.1?report=genbank&log$=prottop&blast_rank=4&RID=8665CEF5016) |
| TonB-dependent receptor [Methylococcales bacterium] | Methylococcales bacterium | 746 | 746 | 98% | 0 | 53.96 | 685 | [MBT7108356.1](https://www.ncbi.nlm.nih.gov/protein/MBT7108356.1?report=genbank&log$=prottop&blast_rank=5&RID=8665CEF5016) |
| TonB-dependent receptor [Methyloglobulus sp.] | Methyloglobulus sp. | 942 | 942 | 98% | 0 | 66.03 | 682 | [NOS75890.1](https://www.ncbi.nlm.nih.gov/protein/NOS75890.1?report=genbank&log$=prottop&blast_rank=6&RID=8665CEF5016) |
| TonB-dependent receptor [Methylococcales bacterium] | Methylococcales bacterium | 749 | 749 | 98% | 0 | 54.56 | 685 | [MBT3506297.1](https://www.ncbi.nlm.nih.gov/protein/MBT3506297.1?report=genbank&log$=prottop&blast_rank=7&RID=8665CEF5016) |
| TonB-dependent receptor [Methylococcales bacterium] | Methylococcales bacterium | 749 | 749 | 98% | 0 | 54.56 | 685 | [MBT4766168.1](https://www.ncbi.nlm.nih.gov/protein/MBT4766168.1?report=genbank&log$=prottop&blast_rank=8&RID=8665CEF5016) |
| TonB-dependent receptor [Methylococcales bacterium] | Methylococcales bacterium | 747 | 747 | 98% | 0 | 54.56 | 685 | [MBT4664543.1](https://www.ncbi.nlm.nih.gov/protein/MBT4664543.1?report=genbank&log$=prottop&blast_rank=9&RID=8665CEF5016) |
| TonB-dependent receptor [Methylococcales bacterium] | Methylococcales bacterium | 746 | 746 | 98% | 0 | 54.41 | 685 | [MBT3814959.1](https://www.ncbi.nlm.nih.gov/protein/MBT3814959.1?report=genbank&log$=prottop&blast_rank=10&RID=8665CEF5016) |
| TonB-dependent receptor [Methylococcales bacterium] | Methylococcales bacterium | 745 | 745 | 98% | 0 | 54.41 | 685 | [MBT6523188.1](https://www.ncbi.nlm.nih.gov/protein/MBT6523188.1?report=genbank&log$=prottop&blast_rank=11&RID=8665CEF5016) |
| TPA: hypothetical protein [Methylococcales bacterium] | Methylococcales bacterium | 672 | 672 | 95% | 0 | 50.95 | 715 | [HIN68511.1](https://www.ncbi.nlm.nih.gov/protein/HIN68511.1?report=genbank&log$=prottop&blast_rank=12&RID=8665CEF5016) |
| hypothetical protein DSY87_06945 [Methylococcus sp.] | Methylococcus sp. | 682 | 682 | 91% | 0 | 54.32 | 639 | [RUM52252.1](https://www.ncbi.nlm.nih.gov/protein/RUM52252.1?report=genbank&log$=prottop&blast_rank=13&RID=8665CEF5016) |
| TonB-dependent receptor [Methylomonas sp.] | Methylomonas sp. | 63.2 | 63.2 | 88% | 5.00E-09 | 20.72 | 614 | [MBS3963638.1](https://www.ncbi.nlm.nih.gov/protein/MBS3963638.1?report=genbank&log$=prottop&blast_rank=14&RID=8665CEF5016) |
| TonB-dependent receptor [Methylococcaceae bacterium] | Methylococcaceae bacterium | 55.5 | 55.5 | 78% | 1.00E-06 | 21.78 | 663 | [NOT86334.1](https://www.ncbi.nlm.nih.gov/protein/NOT86334.1?report=genbank&log$=prottop&blast_rank=15&RID=8665CEF5016) |
| TonB-dependent receptor [Methylococcales bacterium] | Methylococcales bacterium | 587 | 587 | 78% | 0 | 53.61 | 545 | [MBT4347282.1](https://www.ncbi.nlm.nih.gov/protein/MBT4347282.1?report=genbank&log$=prottop&blast_rank=16&RID=8665CEF5016) |
| TonB-dependent receptor [Methylomonas sp. DH-1] | Methylomonas sp. DH-1 | 64.3 | 64.3 | 76% | 2.00E-09 | 21.49 | 697 | [WP_064020155.1](https://www.ncbi.nlm.nih.gov/protein/WP_064020155.1?report=genbank&log$=prottop&blast_rank=17&RID=8665CEF5016) |
| TonB-dependent receptor [Methylovulum sp.] | Methylovulum sp. | 60.5 | 60.5 | 75% | 3.00E-08 | 23.22 | 619 | [TAL48853.1](https://www.ncbi.nlm.nih.gov/protein/TAL48853.1?report=genbank&log$=prottop&blast_rank=18&RID=8665CEF5016) |
| TPA: TonB-dependent receptor [Methyloprofundus sp.] | Methyloprofundus sp. | 59.7 | 59.7 | 74% | 6.00E-08 | 23.42 | 609 | [HIG65757.1](https://www.ncbi.nlm.nih.gov/protein/HIG65757.1?report=genbank&log$=prottop&blast_rank=19&RID=8665CEF5016) |
| **Metal_2337** | **EIC30073.1** | **H8GGW0** | **TonB-dependent siderophore receptor** | | | | | |
| *Description* | *Scientific Name* | *Max Score* | *Total Score* | *Query Cover* | *E value* | *Per. ident* | *Acc. Len* | *Accession* |
| TonB-dependent siderophore receptor [Methylicorpusculum oleiharenae] | Methylicorpusculum oleiharenae | 1163 | 1163 | 100% | 0 | 72.93 | 782 | [WP_159658092.1](https://www.ncbi.nlm.nih.gov/protein/WP_159658092.1?report=genbank&log$=prottop&blast_rank=1&RID=866ATZBX016) |
| TonB-dependent siderophore receptor [Methylomicrobium sp. RS1] | Methylomicrobium sp. RS1 | 1148 | 1148 | 100% | 0 | 71.96 | 771 | [WP_202053524.1](https://www.ncbi.nlm.nih.gov/protein/WP_202053524.1?report=genbank&log$=prottop&blast_rank=2&RID=866ATZBX016) |
| TonB-dependent siderophore receptor [Methylosarcina fibrata] | Methylosarcina fibrata | 1097 | 1097 | 100% | 0 | 69.55 | 771 | [WP_083918196.1](https://www.ncbi.nlm.nih.gov/protein/WP_083918196.1?report=genbank&log$=prottop&blast_rank=3&RID=866ATZBX016) |
| TonB-dependent siderophore receptor [Methylomonas koyamae] | Methylomonas koyamae | 1046 | 1046 | 99% | 0 | 67.64 | 766 | [WP_157197793.1](https://www.ncbi.nlm.nih.gov/protein/WP_157197793.1?report=genbank&log$=prottop&blast_rank=4&RID=866ATZBX016) |
| TonB-dependent siderophore receptor [Methylobacter sp.] | Methylobacter sp. | 263 | 263 | 97% | 3.00E-75 | 29.22 | 778 | [PPD05063.1](https://www.ncbi.nlm.nih.gov/protein/PPD05063.1?report=genbank&log$=prottop&blast_rank=5&RID=866ATZBX016) |
| TonB-dependent siderophore receptor [Methylococcaceae bacterium] | Methylococcaceae bacterium | 243 | 243 | 97% | 5.00E-68 | 27.9 | 772 | [NOT12940.1](https://www.ncbi.nlm.nih.gov/protein/NOT12940.1?report=genbank&log$=prottop&blast_rank=6&RID=866ATZBX016) |
| TonB-dependent siderophore receptor [Methylomicrobium] | Methylomicrobium | 1550 | 1550 | 96% | 0 | 99.87 | 763 | [WP_245549351.1](https://www.ncbi.nlm.nih.gov/protein/WP_245549351.1?report=genbank&log$=prottop&blast_rank=7&RID=866ATZBX016) |
| **Metal_3768** | **EIC31412.1** | **H8GIA0** | **ATP synthase epsilon chain** | | | | | |
| *Description* | *Scientific Name* | *Max Score* | *Total Score* | *Query Cover* | *E value* | *Per. ident* | *Acc. Len* | *Accession* |
| F0F1 ATP synthase subunit epsilon [Methylomicrobium agile] | Methylomicrobium agile | 182 | 182 | 100% | 2.00E-59 | 99.29 | 140 | [WP_031430124.1](https://www.ncbi.nlm.nih.gov/protein/WP_031430124.1?report=genbank&log$=prottop&blast_rank=1&RID=866G6M9M016) |
| F0F1 ATP synthase subunit epsilon [Methylomicrobium lacus] | Methylomicrobium lacus | 173 | 173 | 100% | 1.00E-55 | 92.86 | 140 | [WP_024298897.1](https://www.ncbi.nlm.nih.gov/protein/WP_024298897.1?report=genbank&log$=prottop&blast_rank=2&RID=866G6M9M016) |
| F0F1 ATP synthase subunit epsilon [Methylomicrobium sp. RS1] | Methylomicrobium sp. RS1 | 169 | 169 | 100% | 4.00E-54 | 95 | 140 | [WP_202051583.1](https://www.ncbi.nlm.nih.gov/protein/WP_202051583.1?report=genbank&log$=prottop&blast_rank=3&RID=866G6M9M016) |
| F0F1 ATP synthase subunit epsilon [Methylomicrobium sp.] | Methylomicrobium sp. | 164 | 164 | 100% | 3.00E-52 | 83.57 | 140 | [MBS3952104.1](https://www.ncbi.nlm.nih.gov/protein/MBS3952104.1?report=genbank&log$=prottop&blast_rank=4&RID=866G6M9M016) |
| F0F1 ATP synthase subunit epsilon [Methylotuvimicrobium buryatense] | Methylotuvimicrobium buryatense | 164 | 164 | 100% | 3.00E-52 | 84.29 | 140 | [WP_026130213.1](https://www.ncbi.nlm.nih.gov/protein/WP_026130213.1?report=genbank&log$=prottop&blast_rank=5&RID=866G6M9M016) |
| F0F1 ATP synthase subunit epsilon [Methyloglobulus sp.] | Methyloglobulus sp. | 164 | 164 | 100% | 5.00E-52 | 82.86 | 140 | [NOU45254.1](https://www.ncbi.nlm.nih.gov/protein/NOU45254.1?report=genbank&log$=prottop&blast_rank=6&RID=866G6M9M016) |
| F0F1 ATP synthase subunit epsilon [Methylicorpusculum oleiharenae] | Methylicorpusculum oleiharenae | 164 | 164 | 100% | 5.00E-52 | 83.57 | 140 | [WP_159657005.1](https://www.ncbi.nlm.nih.gov/protein/WP_159657005.1?report=genbank&log$=prottop&blast_rank=7&RID=866G6M9M016) |
| F0F1 ATP synthase subunit epsilon [Methyloglobulus sp.] | Methyloglobulus sp. | 164 | 164 | 100% | 7.00E-52 | 83.57 | 140 | [NOS74980.1](https://www.ncbi.nlm.nih.gov/protein/NOS74980.1?report=genbank&log$=prottop&blast_rank=8&RID=866G6M9M016) |
| **Metal_2634** | **EIC30346.1** | **H8GJI6** | **LPS-assembly protein LptD** | | | | | |
| *Description* | *Scientific Name* | *Max Score* | *Total Score* | *Query Cover* | *E value* | *Per. ident* | *Acc. Len* | *Accession* |
| LPS assembly protein LptD [Methylomicrobium sp. RS1] | Methylomicrobium sp. RS1 | 1914 | 1914 | 100% | 0 | 94.53 | 1004 | [WP_202052202.1](https://www.ncbi.nlm.nih.gov/protein/WP_202052202.1?report=genbank&log$=prottop&blast_rank=1&RID=866V0GYK016) |
| LPS assembly protein LptD [Methylomicrobium] | Methylomicrobium | 1963 | 1963 | 98% | 0 | 100 | 983 | [WP_031431959.1](https://www.ncbi.nlm.nih.gov/protein/WP_031431959.1?report=genbank&log$=prottop&blast_rank=2&RID=866V0GYK016) |
| LPS assembly protein LptD [Methylococcaceae bacterium WWC4] | Methylococcaceae bacterium WWC4 | 799 | 799 | 98% | 0 | 42.69 | 953 | [NJA07554.1](https://www.ncbi.nlm.nih.gov/protein/NJA07554.1?report=genbank&log$=prottop&blast_rank=3&RID=866V0GYK016) |
| LPS assembly protein LptD [Methylomonas koyamae] | Methylomonas koyamae | 799 | 799 | 97% | 0 | 42.45 | 962 | [WP_064031546.1](https://www.ncbi.nlm.nih.gov/protein/WP_064031546.1?report=genbank&log$=prottop&blast_rank=4&RID=866V0GYK016) |
| LPS assembly protein LptD [Methylomonas sp. LWB] | Methylomonas sp. LWB | 799 | 799 | 97% | 0 | 43.17 | 953 | [WP_071155450.1](https://www.ncbi.nlm.nih.gov/protein/WP_071155450.1?report=genbank&log$=prottop&blast_rank=5&RID=866V0GYK016) |
| LPS-assembly protein LptD [Methylomicrobium lacus] | Methylomicrobium lacus | 1520 | 1520 | 97% | 0 | 76.04 | 986 | [WP_024296629.1](https://www.ncbi.nlm.nih.gov/protein/WP_024296629.1?report=genbank&log$=prottop&blast_rank=6&RID=866V0GYK016) |
| LPS assembly protein LptD [Candidatus Methylobacter favarea] | Candidatus Methylobacter favarea | 1119 | 1119 | 96% | 0 | 56.61 | 949 | [WP_174627629.1](https://www.ncbi.nlm.nih.gov/protein/WP_174627629.1?report=genbank&log$=prottop&blast_rank=7&RID=866V0GYK016) |
| **Metal_0880** | **EIC28704.1** | **H8GFV3** | **Uncharacterized protein** | | | | |  |
| *Description* | *Scientific Name* | *Max Score* | *Total Score* | *Query Cover* | *E value* | *Per. ident* | *Acc. Len* | *Accession* |
| hypothetical protein [Methylomicrobium album] | Methylomicrobium album | 1837 | 1837 | 100% | 0 | 100 | 4036 | [WP_005369970.1](https://www.ncbi.nlm.nih.gov/protein/WP_005369970.1?report=genbank&log$=prottop&blast_rank=1&RID=8671YNNA016) |
| hypothetical protein [Methylomicrobium lacus] | Methylomicrobium lacus | 437 | 655 | 95% | 8.00E-128 | 49.05 | 3720 | [WP_024298611.1](https://www.ncbi.nlm.nih.gov/protein/WP_024298611.1?report=genbank&log$=prottop&blast_rank=2&RID=8671YNNA016) |
| **Metal_0879** | **EIC28703.1** | **H8GFV2** | **type I secretion outer membrane protein, TolC family** | | | | | |
| *Description* | *Scientific Name* | *Max Score* | *Total Score* | *Query Cover* | *E value* | *Per. ident* | *Acc. Len* | *Accession* |
| TolC family outer membrane protein [Methylococcaceae bacterium] | Methylococcaceae bacterium | 483 | 483 | 95% | 4.00E-168 | 56.4 | 441 | [NOT13939.1](https://www.ncbi.nlm.nih.gov/protein/NOT13939.1?report=genbank&log$=prottop&blast_rank=1&RID=867EPX4F016) |
| TolC family outer membrane protein [Methylotuvimicrobium buryatense] | Methylotuvimicrobium buryatense | 562 | 562 | 94% | 0 | 62.87 | 513 | [WP_138767192.1](https://www.ncbi.nlm.nih.gov/protein/WP_138767192.1?report=genbank&log$=prottop&blast_rank=2&RID=867EPX4F016) |
| TolC family outer membrane protein [Crenothrix polyspora] | Crenothrix polyspora | 197 | 197 | 94% | 1.00E-56 | 30.62 | 467 | [WP_176371095.1](https://www.ncbi.nlm.nih.gov/protein/WP_176371095.1?report=genbank&log$=prottop&blast_rank=3&RID=867EPX4F016) |
| putative Type I secretion outer membrane protein [Methylotuvimicrobium alcaliphilum 20Z] | Methylotuvimicrobium alcaliphilum 20Z | 559 | 559 | 93% | 0 | 61.83 | 520 | [CCE22660.1](https://www.ncbi.nlm.nih.gov/protein/CCE22660.1?report=genbank&log$=prottop&blast_rank=4&RID=867EPX4F016) |
| TolC family outer membrane protein [Methylotuvimicrobium alcaliphilum] | Methylotuvimicrobium alcaliphilum | 558 | 558 | 93% | 0 | 61.83 | 508 | [WP_223842350.1](https://www.ncbi.nlm.nih.gov/protein/WP_223842350.1?report=genbank&log$=prottop&blast_rank=5&RID=867EPX4F016) |
| TolC family outer membrane protein [Methylomicrobium lacus] | Methylomicrobium lacus | 686 | 686 | 92% | 0 | 81.12 | 456 | [WP_024298613.1](https://www.ncbi.nlm.nih.gov/protein/WP_024298613.1?report=genbank&log$=prottop&blast_rank=6&RID=867EPX4F016) |
| hypothetical protein [Methylococcaceae bacterium] | Methylococcaceae bacterium | 170 | 170 | 91% | 2.00E-46 | 28.22 | 464 | [NJD08156.1](https://www.ncbi.nlm.nih.gov/protein/NJD08156.1?report=genbank&log$=prottop&blast_rank=7&RID=867EPX4F016) |
| **Metal_0878** | **EIC28702.1** | **H8GFV1** | **type I secretion membrane fusion protein, HlyD family** | | | | | |
| *Description* | *Scientific Name* | *Max Score* | *Total Score* | *Query Cover* | *E value* | *Per. ident* | *Acc. Len* | *Accession* |
| HlyD family type I secretion periplasmic adaptor subunit [Methylomicrobium lacus] | Methylomicrobium lacus | 764 | 764 | 100% | 0 | 84.77 | 440 | [WP_024298614.1](https://www.ncbi.nlm.nih.gov/protein/WP_024298614.1?report=genbank&log$=prottop&blast_rank=1&RID=867KY5KA013) |
| HlyD family type I secretion periplasmic adaptor subunit [Methylomicrobium sp. RS1] | Methylomicrobium sp. RS1 | 706 | 706 | 100% | 0 | 78.18 | 440 | [WP_202052890.1](https://www.ncbi.nlm.nih.gov/protein/WP_202052890.1?report=genbank&log$=prottop&blast_rank=2&RID=867KY5KA013) |
| HlyD family type I secretion periplasmic adaptor subunit [Methylotuvimicrobium buryatense] | Methylotuvimicrobium buryatense | 632 | 632 | 100% | 0 | 69.61 | 441 | [WP_017842711.1](https://www.ncbi.nlm.nih.gov/protein/WP_017842711.1?report=genbank&log$=prottop&blast_rank=3&RID=867KY5KA013) |
| HlyD family type I secretion periplasmic adaptor subunit [Methylotuvimicrobium alcaliphilum] | Methylotuvimicrobium alcaliphilum | 612 | 612 | 100% | 0 | 67.8 | 441 | [WP_014147460.1](https://www.ncbi.nlm.nih.gov/protein/WP_014147460.1?report=genbank&log$=prottop&blast_rank=4&RID=867KY5KA013) |
| HlyD family type I secretion periplasmic adaptor subunit [Methylomonas paludis] | Methylomonas paludis | 532 | 532 | 100% | 0 | 57.69 | 442 | [WP_215583218.1](https://www.ncbi.nlm.nih.gov/protein/WP_215583218.1?report=genbank&log$=prottop&blast_rank=5&RID=867KY5KA013) |
| **Metal_0877** | **EIC28701.1** | **H8GFV0** | **type I secretion system ABC transporter, PrtD family** | | | | | |
| *Description* | *Scientific Name* | *Max Score* | *Total Score* | *Query Cover* | *E value* | *Per. ident* | *Acc. Len* | *Accession* |
| type I secretion system permease/ATPase [Methylomicrobium lacus] | Methylomicrobium lacus | 1005 | 1005 | 100% | 0 | 90.18 | 570 | [WP_024298615.1](https://www.ncbi.nlm.nih.gov/protein/WP_024298615.1?report=genbank&log$=prottop&blast_rank=1&RID=867WUAC0016) |
| type I secretion system permease/ATPase [Methylobacter sp. S3L5C] | Methylobacter sp. S3L5C | 798 | 798 | 99% | 0 | 69.43 | 579 | [WP_243218477.1](https://www.ncbi.nlm.nih.gov/protein/WP_243218477.1?report=genbank&log$=prottop&blast_rank=2&RID=867WUAC0016) |
| type I secretion system permease/ATPase [Methylococcaceae bacterium] | Methylococcaceae bacterium | 373 | 373 | 98% | 1.00E-121 | 38.33 | 570 | [NJD06278.1](https://www.ncbi.nlm.nih.gov/protein/NJD06278.1?report=genbank&log$=prottop&blast_rank=3&RID=867WUAC0016) |
| type I secretion system permease/ATPase [Candidatus Methylospira mobilis] | Candidatus Methylospira mobilis | 573 | 573 | 98% | 0 | 51.87 | 594 | [WP_153247369.1](https://www.ncbi.nlm.nih.gov/protein/WP_153247369.1?report=genbank&log$=prottop&blast_rank=4&RID=867WUAC0016) |
| type I secretion system permease/ATPase [Methylomicrobium] | Methylomicrobium | 937 | 937 | 97% | 0 | 84.32 | 569 | [WP_031430937.1](https://www.ncbi.nlm.nih.gov/protein/WP_031430937.1?report=genbank&log$=prottop&blast_rank=5&RID=867WUAC0016) |
| peptidase [Methylococcaceae bacterium] | Methylococcaceae bacterium | 805 | 805 | 97% | 0 | 71.04 | 573 | [GDX85526.1](https://www.ncbi.nlm.nih.gov/protein/GDX85526.1?report=genbank&log$=prottop&blast_rank=6&RID=867WUAC0016) |
| type I secretion system permease/ATPase [Methylomonas sp.] | Methylomonas sp. | 798 | 798 | 97% | 0 | 69.06 | 572 | [PPD32759.1](https://www.ncbi.nlm.nih.gov/protein/PPD32759.1?report=genbank&log$=prottop&blast_rank=7&RID=867WUAC0016) |
| type I secretion system permease/ATPase [Methylomonas koyamae] | Methylomonas koyamae | 785 | 785 | 97% | 0 | 68.83 | 575 | [WP_221053505.1](https://www.ncbi.nlm.nih.gov/protein/WP_221053505.1?report=genbank&log$=prottop&blast_rank=8&RID=867WUAC0016) |
| type I secretion system permease/ATPase [Methylomonas koyamae] | Methylomonas koyamae | 784 | 784 | 97% | 0 | 68.83 | 575 | [WP_096876456.1](https://www.ncbi.nlm.nih.gov/protein/WP_096876456.1?report=genbank&log$=prottop&blast_rank=9&RID=867WUAC0016) |
| **Metal_2566** | **EIC30282.1** | **H8GIU2** | **type I secretion system ATPase, LssB family** | | | | | |
| *Description* | *Scientific Name* | *Max Score* | *Total Score* | *Query Cover* | *E value* | *Per. ident* | *Acc. Len* | *Accession* |
| type I secretion system permease/ATPase [Methylomicrobium agile] | Methylomicrobium agile | 1324 | 1324 | 100% | 0 | 99.17 | 724 | [WP_031431945.1](https://www.ncbi.nlm.nih.gov/protein/WP_031431945.1?report=genbank&log$=prottop&blast_rank=1&RID=8680WNJJ013) |
| type I secretion system permease/ATPase [Methylomonas koyamae] | Methylomonas koyamae | 937 | 937 | 96% | 0 | 70.23 | 716 | [WP_140913031.1](https://www.ncbi.nlm.nih.gov/protein/WP_140913031.1?report=genbank&log$=prottop&blast_rank=2&RID=8680WNJJ013) |
| type I secretion system permease/ATPase [Methylococcaceae bacterium WWC4] | Methylococcaceae bacterium WWC4 | 919 | 919 | 96% | 0 | 67.38 | 715 | [NJA07647.1](https://www.ncbi.nlm.nih.gov/protein/NJA07647.1?report=genbank&log$=prottop&blast_rank=3&RID=8680WNJJ013) |
| type I secretion system permease/ATPase [Methylomonas koyamae] | Methylomonas koyamae | 917 | 917 | 96% | 0 | 67.24 | 715 | [WP_064029659.1](https://www.ncbi.nlm.nih.gov/protein/WP_064029659.1?report=genbank&log$=prottop&blast_rank=4&RID=8680WNJJ013) |
| type I secretion system permease/ATPase [Methylococcaceae bacterium] | Methylococcaceae bacterium | 917 | 917 | 96% | 0 | 65.38 | 713 | [NOT83223.1](https://www.ncbi.nlm.nih.gov/protein/NOT83223.1?report=genbank&log$=prottop&blast_rank=5&RID=8680WNJJ013) |
| type I secretion system permease/ATPase [Methylomonas sp. LWB] | Methylomonas sp. LWB | 916 | 916 | 96% | 0 | 67.38 | 715 | [WP_071160497.1](https://www.ncbi.nlm.nih.gov/protein/WP_071160497.1?report=genbank&log$=prottop&blast_rank=6&RID=8680WNJJ013) |
| **Metal_2121** | **EIC29875.1** | **H8GR48** | **type I secretion outer membrane protein, TolC family** | | | | | |
| *Description* | *Scientific Name* | *Max Score* | *Total Score* | *Query Cover* | *E value* | *Per. ident* | *Acc. Len* | *Accession* |
| TolC family outer membrane protein [Methylomicrobium sp. RS1] | Methylomicrobium sp. RS1 | 887 | 887 | 100% | 0 | 95.52 | 469 | [WP_202052365.1](https://www.ncbi.nlm.nih.gov/protein/WP_202052365.1?report=genbank&log$=prottop&blast_rank=1&RID=8684HEFM013) |
| TolC family outer membrane protein [Candidatus Methylobacter oryzae] | Candidatus Methylobacter oryzae | 495 | 495 | 97% | 1.00E-172 | 55.04 | 456 | [WP_127030250.1](https://www.ncbi.nlm.nih.gov/protein/WP_127030250.1?report=genbank&log$=prottop&blast_rank=2&RID=8684HEFM013) |
| outer membrane channel protein [Methylomonas koyamae] | Methylomonas koyamae | 440 | 440 | 96% | 6.00E-151 | 50.22 | 457 | [BBL58608.1](https://www.ncbi.nlm.nih.gov/protein/BBL58608.1?report=genbank&log$=prottop&blast_rank=3&RID=8684HEFM013) |
| channel protein TolC [Methylomonas koyamae] | Methylomonas koyamae | 439 | 439 | 96% | 1.00E-150 | 50.22 | 457 | [ATG90413.1](https://www.ncbi.nlm.nih.gov/protein/ATG90413.1?report=genbank&log$=prottop&blast_rank=4&RID=8684HEFM013) |
| channel protein TolC [Methylobacter sp.] | Methylobacter sp. | 528 | 528 | 96% | 0 | 58.37 | 447 | [PPD23459.1](https://www.ncbi.nlm.nih.gov/protein/PPD23459.1?report=genbank&log$=prottop&blast_rank=5&RID=8684HEFM013) |
| TolC family outer membrane protein [Methylomarinum sp.] | Methylomarinum sp. | 466 | 466 | 96% | 3.00E-161 | 52.32 | 437 | [NOR69952.1](https://www.ncbi.nlm.nih.gov/protein/NOR69952.1?report=genbank&log$=prottop&blast_rank=6&RID=8684HEFM013) |
| TolC family outer membrane protein [Methylomicrobium lacus] | Methylomicrobium lacus | 649 | 649 | 95% | 0 | 71.99 | 467 | [WP_024297425.1](https://www.ncbi.nlm.nih.gov/protein/WP_024297425.1?report=genbank&log$=prottop&blast_rank=7&RID=8684HEFM013) |
| TolC family outer membrane protein [Methylobacter luteus] | Methylobacter luteus | 582 | 582 | 95% | 0 | 65.11 | 455 | [WP_027157674.1](https://www.ncbi.nlm.nih.gov/protein/WP_027157674.1?report=genbank&log$=prottop&blast_rank=8&RID=8684HEFM013) |
| TolC family outer membrane protein [Candidatus Methylobacter favarea] | Candidatus Methylobacter favarea | 526 | 526 | 95% | 0 | 61.69 | 450 | [WP_174624822.1](https://www.ncbi.nlm.nih.gov/protein/WP_174624822.1?report=genbank&log$=prottop&blast_rank=9&RID=8684HEFM013) |
| TolC family outer membrane protein [Methylomicrobium sp.] | Methylomicrobium sp. | 510 | 510 | 95% | 1.00E-178 | 58.44 | 445 | [MBS3953231.1](https://www.ncbi.nlm.nih.gov/protein/MBS3953231.1?report=genbank&log$=prottop&blast_rank=10&RID=8684HEFM013) |
| channel protein TolC [Methyloglobulus sp.] | Methyloglobulus sp. | 530 | 530 | 95% | 0 | 59.6 | 447 | [MSS76617.1](https://www.ncbi.nlm.nih.gov/protein/MSS76617.1?report=genbank&log$=prottop&blast_rank=11&RID=8684HEFM013) |
| channel protein TolC [Methylobacter sp.] | Methylobacter sp. | 494 | 494 | 95% | 2.00E-172 | 56.82 | 455 | [TAN67820.1](https://www.ncbi.nlm.nih.gov/protein/TAN67820.1?report=genbank&log$=prottop&blast_rank=12&RID=8684HEFM013) |
| outer membrane protein TolC [Methyloglobulus morosus KoM1] | Methyloglobulus morosus KoM1 | 518 | 518 | 95% | 0 | 62.11 | 475 | [ESS73184.1](https://www.ncbi.nlm.nih.gov/protein/ESS73184.1?report=genbank&log$=prottop&blast_rank=13&RID=8684HEFM013) |

**Supplemental Table S4.** Major S-layer protein (H8GFV3) of M. album BG8 screened against Gammaproteobacteria (taxid:1236) using BLASTp. Other methanotrophic bacteria are denoted in bold text.

| **Metal_0880** | **EIC28704.1** | **H8GFV3** | **Major S-layer protein** | | | | | |
| --- | --- | --- | --- | --- | --- | --- | --- | --- |
| Description | Scientific Name | Max Score | Total Score | Query Cover | E value | Per. ident | Acc. Len | Accession |
| **hypothetical protein [Methylomicrobium lacus]** | **Methylomicrobium lacus** | **435** | **435** | **96%** | **2.00E-128** | **49.05** | **3720** | [**WP_024298611.1**](https://www.ncbi.nlm.nih.gov/protein/WP_024298611.1?report=genbank&log$=prottop&blast_rank=1&RID=891PGCJ3013) |
| hypothetical protein [Allochromatium warmingii] | Allochromatium warmingii | 221 | 221 | 90% | 5.00E-56 | 38.52 | 4262 | [WP_091332676.1](https://www.ncbi.nlm.nih.gov/protein/WP_091332676.1?report=genbank&log$=prottop&blast_rank=2&RID=891PGCJ3013) |
| hypothetical protein CFI10_17000 [Marinobacterium georgiense] | Marinobacterium georgiense | 224 | 224 | 90% | 4.00E-57 | 37.3 | 3651 | [QSR36649.1](https://www.ncbi.nlm.nih.gov/protein/QSR36649.1?report=genbank&log$=prottop&blast_rank=3&RID=891PGCJ3013) |
| VWA domain-containing protein [Marinobacterium georgiense] | Marinobacterium georgiense | 224 | 224 | 90% | 4.00E-57 | 37.3 | 3663 | [WP_206836830.1](https://www.ncbi.nlm.nih.gov/protein/WP_206836830.1?report=genbank&log$=prottop&blast_rank=4&RID=891PGCJ3013) |
| hypothetical protein CFI10_16905 [Marinobacterium georgiense] | Marinobacterium georgiense | 212 | 212 | 90% | 3.00E-53 | 36.48 | 3420 | [QSR36631.1](https://www.ncbi.nlm.nih.gov/protein/QSR36631.1?report=genbank&log$=prottop&blast_rank=5&RID=891PGCJ3013) |
| hypothetical protein [Marinobacterium georgiense] | Marinobacterium georgiense | 212 | 212 | 90% | 3.00E-53 | 36.48 | 3409 | [WP_206836786.1](https://www.ncbi.nlm.nih.gov/protein/WP_206836786.1?report=genbank&log$=prottop&blast_rank=6&RID=891PGCJ3013) |
| VWA domain-containing protein [Marinobacterium georgiense] | Marinobacterium georgiense | 159 | 222 | 80% | 2.00E-36 | 35.75 | 3873 | [WP_206836888.1](https://www.ncbi.nlm.nih.gov/protein/WP_206836888.1?report=genbank&log$=prottop&blast_rank=7&RID=891PGCJ3013) |
| hypothetical protein [Spongiibacter nanhainus] | Spongiibacter nanhainus | 123 | 195 | 80% | 6.00E-25 | 40.35 | 4067 | [WP_198569587.1](https://www.ncbi.nlm.nih.gov/protein/WP_198569587.1?report=genbank&log$=prottop&blast_rank=8&RID=891PGCJ3013) |
| DUF4214 domain-containing protein [Halomonas lysinitropha] | Halomonas lysinitropha | 87.4 | 148 | 48% | 6.00E-14 | 35.97 | 1604 | [WP_151442986.1](https://www.ncbi.nlm.nih.gov/protein/WP_151442986.1?report=genbank&log$=prottop&blast_rank=9&RID=891PGCJ3013) |
| VWA domain-containing protein [Chromatium okenii] | Chromatium okenii | 107 | 107 | 31% | 5.00E-20 | 39.05 | 4243 | [MBV5310677.1](https://www.ncbi.nlm.nih.gov/protein/MBV5310677.1?report=genbank&log$=prottop&blast_rank=10&RID=891PGCJ3013) |
| DUF4214 domain-containing protein [Pseudomaricurvus alkylphenolicus] | Pseudomaricurvus alkylphenolicus | 51.6 | 51.6 | 22% | 0.005 | 31.68 | 4338 | [WP_166995439.1](https://www.ncbi.nlm.nih.gov/protein/WP_166995439.1?report=genbank&log$=prottop&blast_rank=11&RID=891PGCJ3013) |
| DUF4214 domain-containing protein [Halomonas sp.] | Halomonas sp. | 59.7 | 59.7 | 21% | 2.00E-05 | 36.56 | 8219 | [MBL1269843.1](https://www.ncbi.nlm.nih.gov/protein/MBL1269843.1?report=genbank&log$=prottop&blast_rank=12&RID=891PGCJ3013) |
| retention module-containing protein [Gammaproteobacteria bacterium] | Gammaproteobacteria bacterium | 53.9 | 53.9 | 19% | 0.001 | 41.4 | 3697 | [RYY04220.1](https://www.ncbi.nlm.nih.gov/protein/RYY04220.1?report=genbank&log$=prottop&blast_rank=13&RID=891PGCJ3013) |
| hypothetical protein [Marinobacterium georgiense] | Marinobacterium georgiense | 64.3 | 64.3 | 19% | 7.00E-07 | 43.37 | 7857 | [WP_206836748.1](https://www.ncbi.nlm.nih.gov/protein/WP_206836748.1?report=genbank&log$=prottop&blast_rank=14&RID=891PGCJ3013) |
| calcium-binding protein [Marinobacterium litorale] | Marinobacterium litorale | 58.5 | 58.5 | 15% | 4.00E-05 | 39.73 | 6883 | [WP_027852952.1](https://www.ncbi.nlm.nih.gov/protein/WP_027852952.1?report=genbank&log$=prottop&blast_rank=15&RID=891PGCJ3013) |
| DUF4214 domain-containing protein [Halomonas sp.] | Halomonas sp. | 53.9 | 53.9 | 12% | 0.001 | 37.96 | 4124 | [MBL1266195.1](https://www.ncbi.nlm.nih.gov/protein/MBL1266195.1?report=genbank&log$=prottop&blast_rank=16&RID=891PGCJ3013) |
| hypothetical protein [Halomonas sp.] | Halomonas sp. | 53.1 | 53.1 | 12% | 0.002 | 37.96 | 2115 | [MBL1269839.1](https://www.ncbi.nlm.nih.gov/protein/MBL1269839.1?report=genbank&log$=prottop&blast_rank=17&RID=891PGCJ3013) |
